# Myotropes unveil two myosin cycles with distinct kinetics and stroke size

**DOI:** 10.64898/2026.09.11.750962

**Authors:** Albin E. Berg, Tianbang Wang, Emrulla Spahiu, Lok Priya Velayuthan, Marlene Norrby, Sven Tågerud, Cristobal dos Remedios, Theresia Kraft, Arnab Nayak, Marko Usaj, Alf Månsson, Mamta Amrute-Nayak

## Abstract

Cardiac contraction relies on cyclic interactions between myosin II motors and actin filaments, powered by the free energy of ATP turnover^1^. Targeting of this fundamental process has emerged as a promising therapeutic strategy; Mavacamten, a cardiac myosin inhibitor was recently approved for treating obstructive hypertrophic cardiomyopathy whereas omecamtiv mecarbil was developed as a myosin activator for heart failure. Both myotropes bind to the same myosin pocket but their mechanisms of action remain incompletely understood. Here, we investigate these using optical tweezers mechanoenzymology and single-molecule fluorescence-based actomyosin kinetics applied to β-myosin from donor hearts and recombinant human β-myosin subfragment 1. Our findings reveal unexpected layers of complexity. Both mavacamten and omecamtiv mecarbil partially slow the ATP turnover, giving one fast (similar to physiological) and one drug-induced slow ATP turnover cycle at saturating drug concentrations with different rate-limiting steps for the two compounds. In one of the parallel cycles, mavacamten increases stroke size and appreciably delays the power-stroke, allowing direct visualization of a strongly bound pre-power-stroke state. No power-stroke delay is seen in the other cycle, but a reduced stroke size is observed. Our findings uncover a new paradigm in cardiac regulation at the single-molecule level, demonstrating partial inhibition of myosin as a powerful therapeutic strategy.

## Main

The force-generating interaction between myosin II and actin is an emerging drug target in several diseases ^2–6^ particularly cardiac conditions. Major emphasis has been on hypertrophic cardiomyopathy (HCM) the leading cause of sudden cardiac death in young people with the recent approval of the myosin inhibitor mavacamten (Mava) for subtypes of the disease ^7^. Heart failure, a leading cause of hospitalization with a poor prognosis has also attracted interest, first with the cardiac myosin activator omecamtiv mecarbil (OM) ^8,9^. More recently, additional myotropes have been developed including aficamten ^10,11^ showing inhibitory actions, targeting HCM, and danicamtiv ^12,13^ with pharmacodynamic similarities to OM.

Intriguingly, the pioneering compounds Mava and OM differently modulate cardiac contraction –either inhibiting or activating it - despite binding to the same pocket on myosin^14^. Ultrastructural analyses suggest different effects of Mava and OM on myosin dynamics as basis for their divergent effects ^14–16^. There is also consensus that Mava slows release of the ATP-hydrolysis product inorganic phosphate (Pi) from the active site of actin-bound myosin whereas OM has the opposite effect ^6,8,17–19^. However, the mechanistic basis for the major differences in myocardial effects between Mava and OM remains poorly understood. The insights are particularly limited for the clinically approved drug, Mava. For instance, although both structural and biophysical studies demonstrated that OM inhibits the motion-generating lever arm swing (the power-stroke) of myosin^14,15,19,20^ the situation with Mava is unclear. While structural data suggest an inhibiting effect of Mava^14,16^, supporting functional evidence is lacking.

Here, we employed a unique combination of single molecule methods, including optical tweezers mechanics and fluorescence based steady-state and transient kinetics assays, for new mechanistic insights. By studying both human ventricular myosin II isolated from donor hearts and expressed motor domains (e.g. as previously^16,20,21^) we ensure direct human physiological and clinical relevance. Strikingly, the effects of both Mava and OM can be interpreted within one unified explanatory framework: at saturating concentrations both compounds partially redirect the flux through the actin-myosin force-generating cycle from a physiological path to an alternative non-canonical slower ATP cycling route. This behaviour is characteristic of a “partial enzyme inhibitor” where a substantial fraction (>50 % under some conditions) of the myosin molecules continue to undergo normal ATP turnover at saturating drug concentrations. The two compounds differ in their inhibiting effects on the power-stroke, actin-myosin affinity, rate-limiting step of the strongly drug-bound pathway, modulation of the stroke size and the degree of shift to the non-canonical ATP turnover cycle. Together, our findings establish a comprehensive mechanistic framework for myosin-targeting therapeutics and reveal new twists of the chemo-mechanical cycle of cardiac myosin.

### Mava and OM reduce actin filament gliding speed

Actin gliding motility assays employing native human ventricular full-length myosin (βM-II) were used to compare the effects of OM and Mava across a range of concentrations. Both drugs produced a dose-dependent decrease in actin gliding speed (Fig. 1) with half-inhibitory concentration (IC_50_) of 379.16 ± 13.5 nM for Mava and 39.94 ± 2.1 nM for OM, i.e. 10-fold stronger inhibition with OM. Our motility results using native human cardiac myosin are compared with previous data from other myosin preparations^16,17,21^ in Extended Data Fig. 1.

**Fig. 1.**
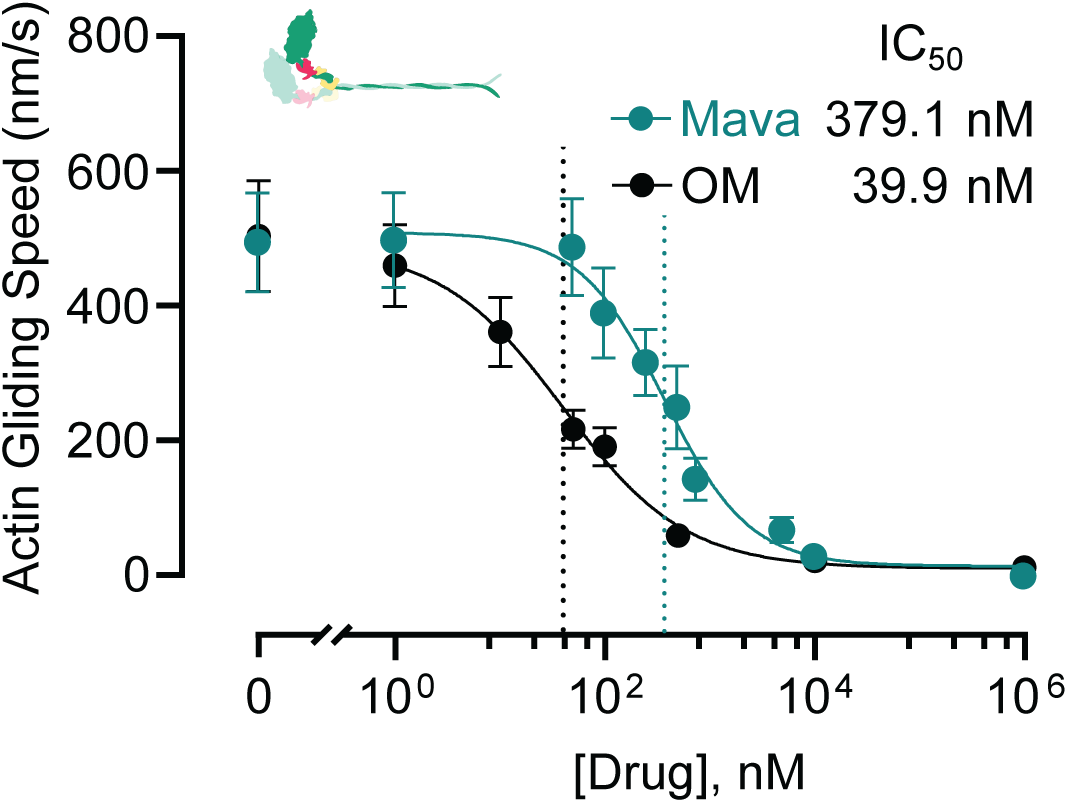
Effects of Mava and OM on the actin filament speed. Dose-response curve showing actin gliding speed of human βM-II in the presence of Mava and OM. Motility experiments were conducted at 2 mM [ATP] and 22°C. Movies were collected from two separate myosin preparations. For each data point, 100-110 filaments were manually tracked using the MTrackJ plugin in ImageJ.

To unravel mechanisms whereby Mava and OM influence myosin activity, we employed single-molecule ATPase and optical trapping experiments. These were applied to full-length human ventricular myosin (βM-II) or myosin subfragment 1 (S1) from donor hearts or human ventricular myosin S1 (AA 1-848) expressed in C2C12 cells. The latter (“S1^E^”) had the heavy chain fused to GFP and endogenous murine skeletal muscle light chains. To minimize unwanted heterogeneities in the single molecule measurements, we employed saturating concentrations of Mava (30 µM) and OM (100 µM) (cf. Fig. S1).

### Similar effects of OM and Mava on ATPase

First, the basal ATPase was evaluated after capturing myosin S1^E^ on anti eGFP antibodies adsorbed to nitrocellulose surfaces (Fig. 2). Total internal reflection fluorescence (TIRF) microscopy was used to record dwell-times (Fig. 2a) attributed to S1^E^-binding and turnover of fluorescent Alexa 647 ATP (Alexa-ATP). The turnover rate constant was greatly reduced from 0.029 ± 0.00075 s^−1^ (mean ± SEM; n=2) under control conditions to 0.007 ± 0.0002 s^−1^ and 0.0062 ± 0.0015 s^−1^ in the presence of either Mava or OM (Supplementary movies 1-3). This was suggested by the slowest phase in exponential fits to cumulative frequency distributions (Fig. 2a-d) constructed as described previously^22,23^. Presumably, the values with the drug are photobleaching-limited^23^, suggesting an underestimated reduction in basal ATPase. A dominating fast phase (rate constant >0.1 s^−1^) under control conditions, previously attributed to non-specific binding of Alexa-ATP to myosin (Fig. S2) ^23^, was not distinguished following drug addition, but its presence is indicated by high error of the fastest rate constant (Fig. 2c).

**Fig. 2.**
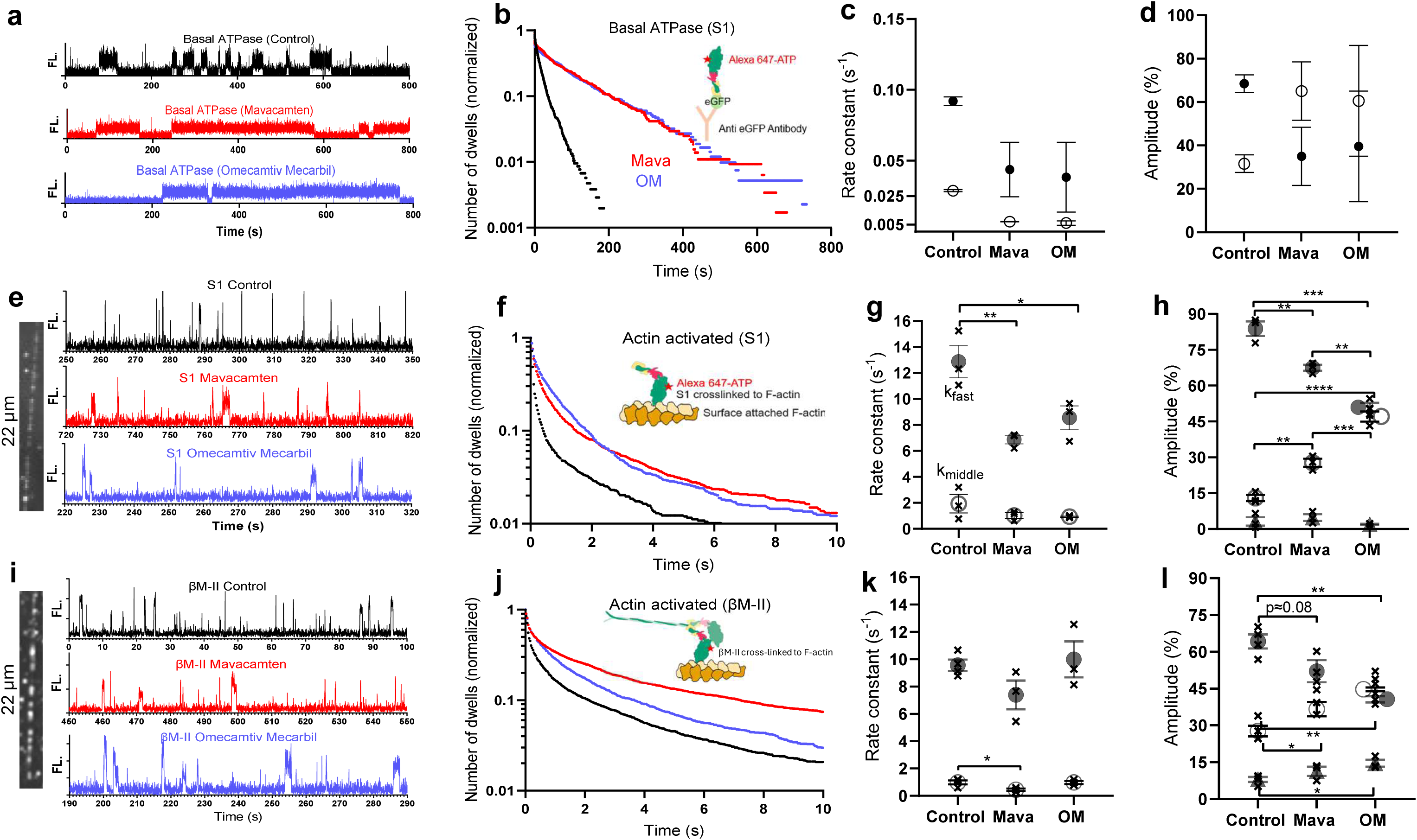
MAVA (30 µM) and OM (100 µM) promote slow basal ATPase and increase the amplitude of a slow process in the cumulative Alexa 647 (“Alexa”) nucleotide dwell time distributions for actin-activated ATP turnover. **a)** Typical time traces of Alexa-nucleotide dwell times with myosin S1^E^ alone, without drug (black), with Mava (red) and with OM (blue). b) Cumulative dwell time distribution of Alexa ATP binding and turnover events on S1^E^ alone (n=2). Control (N_dwell_ = 509 and 544), OM (N_dwell_ = 170 and 221), Mava (N_dwell_ = 171 and 294). Inset: schematic of experiment. c-d) Rate constants and amplitudes extracted from triple exponential fittings to cumulative dwell time distribution in b with the slowest processes (open circles) attributed to ATP turnover. The parameters representing the third unspecific process included in the fitting procedure (see Table S1 and Fig. S2) are not shown for clarity. e) Typical time traces of Alexa-nucleotide dwell times with myosin S1^E^ cross-linked to actin. Color coding as in a. Inset left: Typical Standard deviation time projection of Alexa ATP binding to S1 cross-linked along an actin filament. f) Cumulative dwell time distributions of Alexa ATP binding and turnover events (N_dwell_ ≈ 3000 for each condition, n=3) for S1^E^ crosslinked to actin. Inset, schematic of experiment. g, h) Rate constants and amplitudes from triple exponential fittings. Filled circles: fast process (k_fast_ 6.8-12.8 s^−1^), Open circles: intermediate process (k_middle_ 0.45-1.9 s^−1^). Filled triangles: slow process (k_slow_ 0.04-0.32 s^−1^; not included in g, Crosses: individual fitting results from 3 independent experiments (N_dwell_ ranging from 940 to 1032; mean ± SEM). i) Typical time traces showing Alexa-nucleotide dwell times with full length β-myosin (βM-II cross-linked to actin. Inset, left. Typical standard deviation time projection of Alexa ATP binding to βM-II cross-linked along actin filament. j) Cumulative dwell time distribution of Alexa ATP binding events to βM-II crosslinked to actin (N_dwell_ = 1005-1019 for each condition, n=3-4). Inset. Schematic of experiment. k-l) Rate constants and amplitudes from triple exponential fittings to distributions like those in g-h (k_fast_7.3-9.9 s^−1^ k_middle_ 0.5-0.98 s^−1^, k_slow_ 0.032-0.22 s^−1^; k_slow_ not included in k). Asterisks in g, h, k, l refer to p-values (**** p<0.0001; *** p<0.001, ** p<0.01, * p< 0.05) derived from unpaired ANOVA followed by Sidak’s post hoc test. P-value < 0.1 is also given by number. Data given as mean ± SEM; n = number of independent experiments.

The actin-activated ATPase (Fig. 2e-l) was investigated by the first systematic application of a recently developed single-molecule method^24^ with either S1^E^ or native myosin (βM-II) crosslinked to actin using the zero-length cross-linker 1-Ethyl-3-[3-dimethylaminopropyl]carbodiimide hydrochloride (EDC)^25–27^ (Supplementary movies 4-9). These experiments revealed two dominating turnover components, one fast (k_fast_ >≈10 s^−1^) and one slower (k_middle_ ≈ 1 s^−1^) both in the presence and absence of drug with drug-induced shift towards the slow component as described below. Cumulative frequency distributions (Fig. 2f, j) were generated from durations of Alexa-ATP binding and turnover events in fluorescence intensity time traces (Fig. 2e, i). The cumulative distributions (Fig. 2f) are best described by three exponentials^24^ following a brief lag (∼ 30 ms) (Extended Data Fig. 2) attributed to hydrolysis equilibration, ADP-release (75-120 s^−1^; e.g.^28^ ^29^) and limited camera frame rate (32 ms/frame). For S1^E^, the fastest exponential dominates under control conditions with amplitude A_fast_ ≈ 84 % and rate constant k_fast_ ≈ 13 s^−1^ (Fig. 2g, h). The fastest process also dominates for βM-II with A_fast_ ≈ 64% and k_fast_ ≈ 10 s^−1^ (Fig. 2k-l). Surprisingly, despite saturating drug concentrations, the fast exponential phase persisted (Fig. 2h) with a large amplitude (A_fast_>40 %; A_fast_>≈ A_middle_) upon adding OM or Mava. This striking feature was also observed for βM-II despite greater complexity (Fig. 2l). With both S1^E^ and βM-II (Fig. 2h, l), we observed a shift in the event-fractions towards a slower exponential process (k_middle_ range 0.5-1.9 s^−1^). For S1^E^ (Fig. 2h), the amplitude of the latter process (A_middle_) increased from 12.9 ± 1.3% (control; mean ± SEM) to 27.7 ± 1.7% with Mava and 47 ± 2.1% with OM.

The increase in A_middle_ that was larger with OM than Mava (p<0.01), occurred at the expense of A_fast_ which decreased to a corresponding degree (Fig. 2h, l). These drug-induced changes were associated with reduction of k_fast_ to approximately 50 % with both OM and Mava for S1^E^ (Fig. 2g) but not for βM-II (Fig. 2k). In separate experiments using S1^E^ (Fig. S1), we found that reduction of k_fast_ only occurred at the highest drug concentrations tested (>10 µM) (see also ^15,30^). Moreover, as it was only observed with S1^E^ (Fig. 2k), we do not consider it further below.

To clarify further aspects of drug effects on the actin-myosin interaction we monitored transient S1^E^ binding to, and dissociation from, actin filaments (Fig. 3a). For both control conditions and with 30 µM Mava, the dwell-time distributions for myosin binding to actin were well described by single exponential functions at all tested ATP concentrations (0.1-1µM; Extended Data Fig. 3). The detachment rate constant in these cases increased linearly with [ATP], yielding a second-order rate constant of 7.4 ± 1.2 (mean ± 95% CI) μM^−1^ s^−1^ and 7.8 ± 1.3 μM^−1^ s^−1^, with and without Mava, respectively. The adequate fitting by a single exponential function suggests that rate limiting transition of the actin-activated ATPase (Fig. 2) occurs before or is concomitant with attachment under both control conditions (10-15 s^−1^) and with Mava (1-2 s^−1^). In contrast, the detachment of OM is best described by a double exponential function, with one ATP-sensitive detachment rate constant (6.8 ± 1.9 μM^−1^ s^−1^) and another [ATP]-independent process with rate constant 10.8-17.9 s^−1^ (Fig. 3b). Notably, this latter rate constant is faster than the rate-limiting transition of the slow phase (1-2 s^−1^) with OM (Fig. 2) suggesting the existence of a slow process elsewhere in the cycle (see below). The small effects of Mava and OM on the second order rate constant for ATP-induced actomyosin dissociation is consistent with previous results^6,8,18,20,30,31^.

**Fig. 3.**
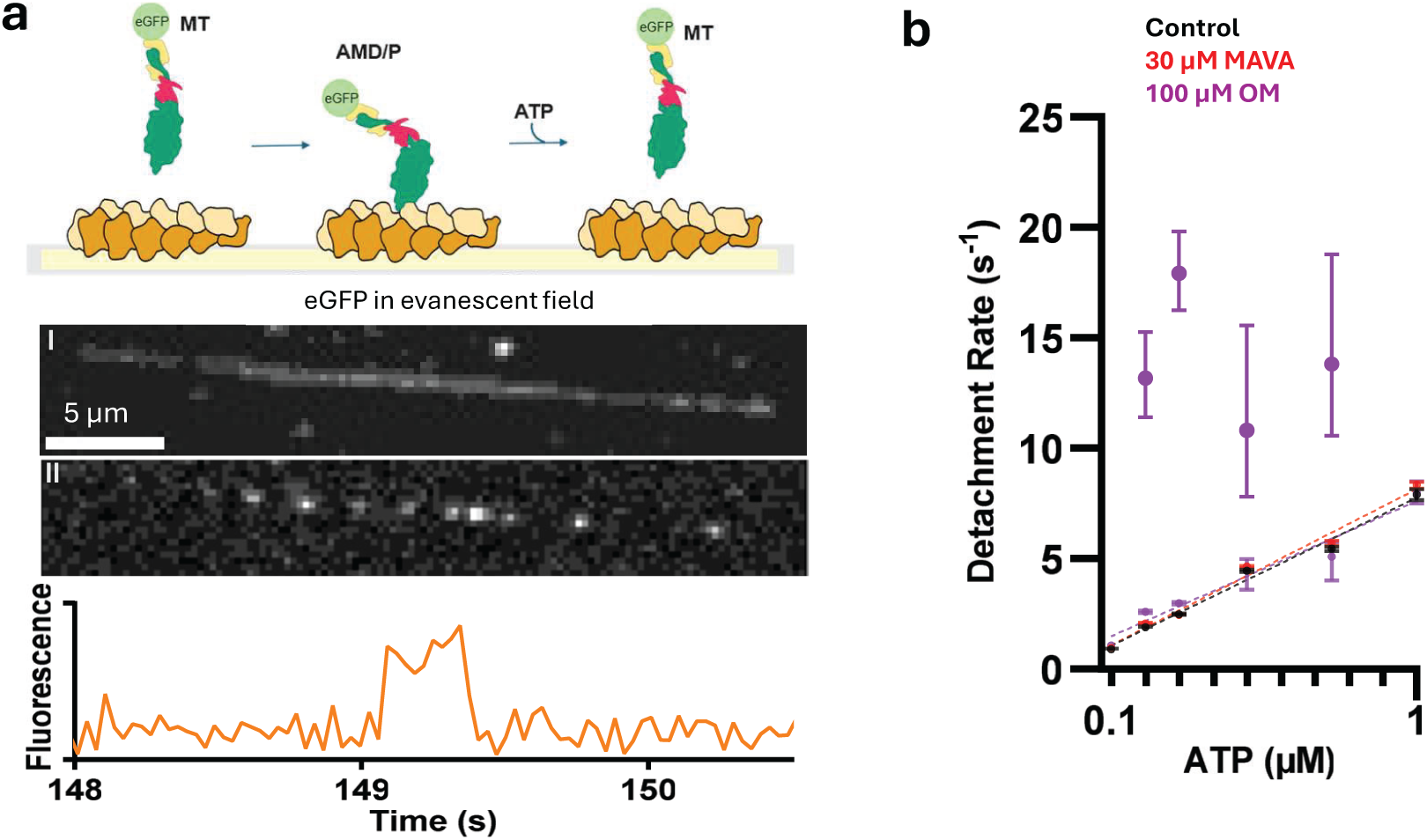
Effects of MAVA and OM on myosin S1^E^ attachment duration at low [ATP]. **a)** Schematic illustration of detachment kinetic assay together with experimental projection of myosin binding event to one actin filament. Images show average eGFP projection from 6100 frames (I) and from 50 frames (II). Bottom panel, a fluorescence intensity over time trace showing an actomyosin binding event. **b)** Actomyosin detachment rate as a function of ATP concentration (0.1-1µM) with either 100 µM OM or 30 µM Mava. Final concentration of DMSO ≤ 1.4%. Error bars represent the 95% CI of the fit. Dashed lines represent the linear regression of different conditions. Experiment was performed at 23±1 °C. Camera capturing time was limited to 32 ms. At the lowest [ATP] (0.1 µM) only one exponential can be fitted for all conditions. For some ATP concentrations, Akike information criteria (AICc) comparisons favor a double-exponential over a single-exponential model with Mava and for the control conditions. However, in these cases, one component contributes 83-99.7% of the total amplitude and the dominant rate constant closely matches the value obtained from the single-exponential model. For all ATP concentrations of 0.1µM < [ATP] < 1 µM the preferred model with OM is a double exponential with the amplitude of the fast and slow process of 20-47% and 53-80, respectively. For [ATP] = 1µM the rate constants of the two processes are too close for meaningful separation, See further Extended Data Fig. 3. N_dwells_ for each condition can be found in **Table S2.**

### Mava delays the power-stroke

For in depth elucidation of the mechanistic basis of Mava and OM-induced changes in the actin filament gliding speed (Fig. 1) and the actin activated ATP turnover kinetics (Fig. 2), we turned to single-molecule optical trapping (Fig. 4a). This technique enables direct measurement of the stroke size of individual motor molecules and the kinetics of actomyosin transitions beyond the rate limiting steps with high spatiotemporal resolution. The information has direct bearing on the myosin-driven actin filament gliding speed and sarcomere shortening velocity as well as the force-generating ability of myosin.

**Fig. 4.**
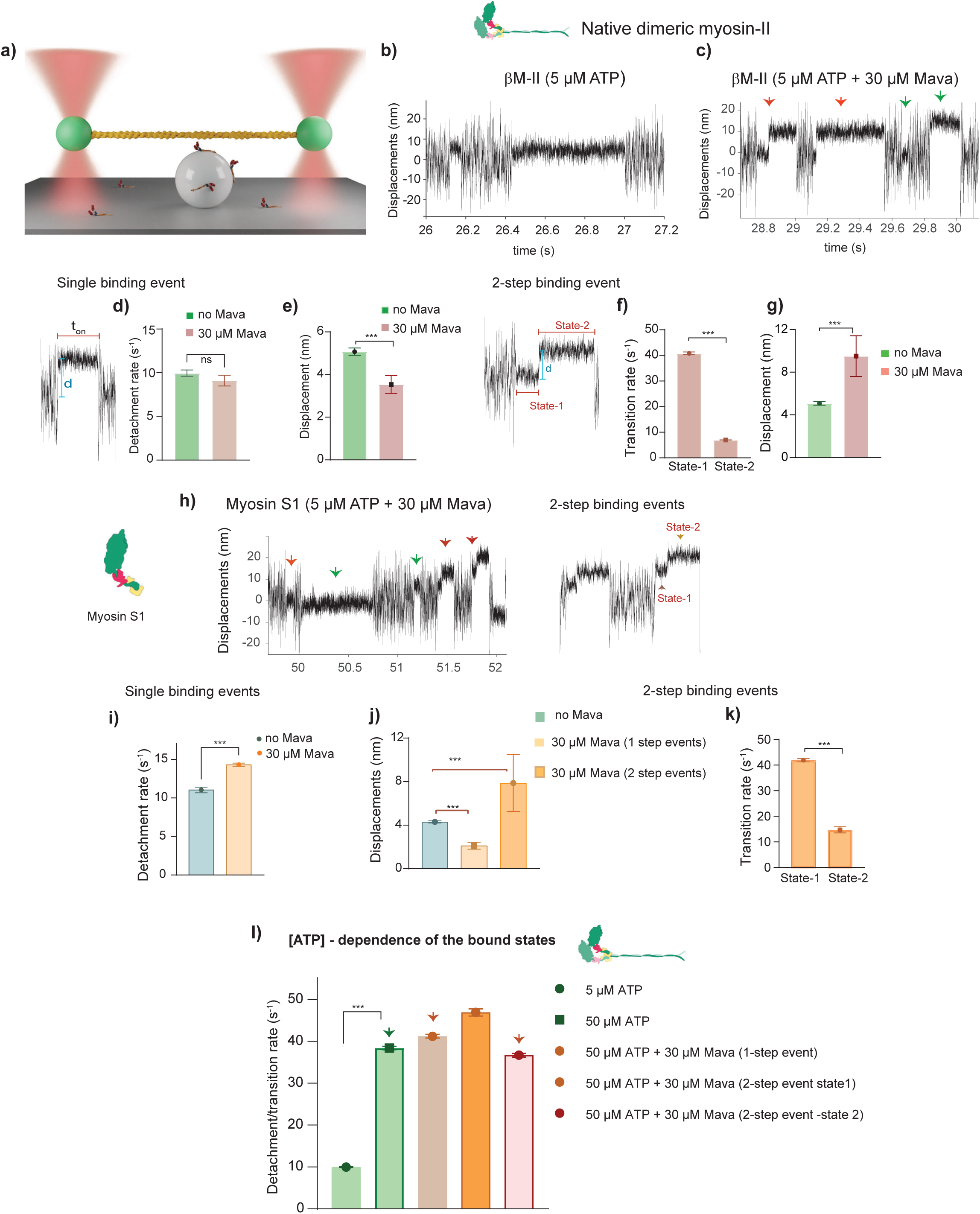
Effects of Mava on native beta cardiac myosin and myosin S1 interaction with actin studied using optical trap. **a)** Optical trap setup. Actin filaments are suspended between the two optically trapped beads; the motors are adsorbed on the nitrocellulose-coated glass bead immobilized on the surface. **b)** The original displacement over time trace at 5 µM ATP from a single βM-II molecule as it interacts with an actin filament. Reduction in the amplitude of the bead’s Brownian motion is indicative of actomyosin interaction events. **c)** A representative data trace from a single βM-II molecule in the presence of 5 µM ATP and 30 µM Mava. The binding events are indicated with green or orange arrows showing two distinct populations, with apparent single- and apparent 2-step binding events, respectively. **d)** Inset - representative apparent single actomyosin interaction event (last interaction event shown in c), t_on_ - the lifetime of actomyosin association, d – displacement from mean dumbbell position. Bar diagram with detachment rate of the single binding events at 5 µM ATP and with and without 30 µM Mava are compared. P = 0.26 (ns). Detachment rate = 1/τ, τ-average time constant estimated by fitting the interaction events to single exponential decay function (as shown in Fig. S4c) **e)** Bar diagram showing the stroke size obtained by using the ‘shift-of-histogram’ method. P = 0.006. Data fits are shown in Fig. S4b and d **f)** Inset - Representative apparent 2-step AM interaction event (first event shown in c), state-1 and state2 are shown, d – distance between the two states. Bar diagram shows transition rates for state-1 and state-2, P< 0.001. **g)** Stroke size at 5 µM ATP and the distance between the two states in the presence of 30 µM Mava are compared. P< 0.001. **h)** Effect of Mava on beta cardiac S1 interaction with actin. Representative original data trace with interaction events between single headed myosin S1 and actin filament showing apparent single and 2-step events are indicated with green and brown arrows, respectively. Inset highlights the 2-step events in the presence of 5 µM ATP and 30 µM Mava (last two events in the data trace). **i)** In a bar diagram for single binding events, actomyosin detachment rates in the presence under indicated conditions are compared. P < 0.001. **j)** Stroke sizes and, or distance between two states for 2-step events are compared between the indicated conditions. P < 0.001. **k)** Transitions rates for state-1 and state-2 for apparent 2-step actomyosin interaction event. P < 0.001. **l)** ATP concentration dependence of the interaction lifetime in the presence of Mava under indicated conditions. The histograms and data fits used to derive the mean values plotted in different bar diagrams are provided in the supplemental information (SI), together with number of analyzed myosin molecules (N) and number of events (n). Error bars in the bar diagrams are mean ± SEM. The mean and SEM from fitted data for indicated conditions were subjected to Welch’s t-test to probe the statistical significance for indicated pair. *** - P <0.001 indicated statistically significant difference. Ns – not significantly different.

We hypothesized that if Mava acts by binding and stabilizing the sequestered (OFF) state of the myosin heads as proposed earlier^14,16,32–34^, no actomyosin interaction events would be observed in our single-molecule setup. Alternatively, any binding event that does occur would represent Mava-independent actin-myosin association proceeding through a normal crossbridge cycle. Fig. 4b and c show displacement over time traces obtained from dimeric full length βM-II in the presence and absence of 30 µM Mava at 5 µM ATP demonstrating a characteristic example of intermittent single actomyosin binding events in the presence of ATP. Strikingly, in contrast to control conditions, two distinct populations of events were persistently observed in the presence of Mava. One of the populations was characterized by apparent single step events (Fig. 4b, Fig. S3a) whereas the other population showed two-step events (Fig. 4c; Fig. S3b). We treated the two populations separately and analyzed individual cardiac myosin molecules for their interaction with actin as described previously^35,36^. Actomyosin detachment/transition rates (1/τ) were determined from the average actomyosin-bound lifetimes (τ) (Fig. S4a) and the stroke size (d) was estimated as (i) the shift-of-the-histogram from the mean dumbbell position (Fig. S4b) or (ii) the distance between the two states in two-step events (indicated in Fig. 4f inset). The population with one-step events showed a similar actomyosin detachment rate compared to the no Mava condition at 5 µM ATP, i.e., ∼ 10 s^−1^ (Fig. 4d and FigS4c); however, the stroke size was reduced from 5 to 3.3 nm (Fig. 4e and S4d). For the population with 2-step events, the transition rate from state-1 to state-2 was ∼ 40 s^−1^ (Fig. 4f). The detachment rate from state-2 (∼7 s^−1^) was similar as without Mava and single-step events with Mava (Fig. 4f and S4e-f). A larger stroke size/displacement of about 9.5 ± 1.9 nm between the two states was observed for the two-step population (Fig. 4g). The 2-step events comprised nearly 36 ± 6.2% of the total actomyosin interaction events.

The two-step binding events occurred on the plus side of actin as shown in the data traces Figure 4c and h after initial association of actin and myosin. The polarity of the actin filament determines the direction of the working stroke, i.e., myosin II, a plus end directed motor binds the actin filament towards the plus end and generates a force-generating displacement/stroke towards the minus end. The initial actomyosin binding of 2-step events was observed at different positions along the actin filament throughout the bead or dumbbell fluctuation range of about 60 nm (i.e., ± 30 nm from mean dumbbell position, cf. Fig. 4c). However, strikingly, the stepping direction was the same for various binding events within a data record for a single myosin molecule. This consistency was also observed when the same actin dumbbell interacted with different myosin molecules. These results indicate an overall directional bias in stepping along the actin filament. Importantly, the observation that the stroke size of the single-step events was reduced by Mava suggests that the drug affects both the single-step and 2-step binding events.

One possible explanation for the 2-step binding events in the presence of Mava may be the sequential binding of two heads of native dimeric myosin employed in the experiments. To examine this hypothesis, we generated single headed myosin subfragment-1(S1) by proteolytic digestion of native cardiac myosin. Importantly, like native myosin, the data records of the actomyosin interactions for S1 revealed two populations, with 2-step events comprising 31.82 ± 9.2% of the total events (Figs. 4h-k, S3c), convincingly showing that the 2-step events represent features of single motor domains interacting with actin. Again, the population with 1-step events showed a rather similar actomyosin detachment rate to that of no Mava condition, i.e., 11 and 14 s^−1^, respectively (Fig. 4i, FigS4i), but reduced stroke size from 4.3 nm to 2 nm (Fig. 4j and S4j). Ensemble average analysis of the 1-step events with Mava suggest that the size of both the 1^st^ and 2^nd^ sub-strokes are reduced compared to no Mava control for both native and S1 (Extended Data Fig. 4a-d). For the 2-step population, the state-1 to state-2 transition rate constant was similar (∼42 s^−1^) as with native myosin, while state-2 showed a comparable dissociation rate to that of the apparent single events (Fig. 4k and Fig S4 k-l). Again, like for native myosin, a larger stroke size/displacement of about 7.9 ± 2.6 nm between the two states was observed for the S1 two-step population (Fig. 4j).

Following the transition of myosin from the weakly to the strongly actin-bound AM.ADP.Pi state under normal conditions (no drug), rapid Pi release and associated working stroke occur in < 2 ms^37^. Therefore, with temporal resolution of the trap in this range, the duration of the actomyosin interaction primarily reports the lifetimes of the ADP-bound and the nucleotide-free ‘rigor’ actomyosin state. The strongly bound AM.ADP.Pi state is too short-lived to be detected, typically giving apparent 1-step events under no drug conditions.

Biochemical studies have shown that Mava decreases the rate of Pi release^6,17^ but the temporal relationship of Pi-release to the working stroke is debated^38–40^. One possibility is that the rate constant of ∼40 s^−1^ found in the trapping data is associated with the delayed Pi-release. However, as further discussed below, a transition rate constant of this magnitude is too high to be rate-limiting (rate constant 1-2 s^−1^). Nonetheless, it is conceivable that state-1 in the 2-step events reflect a delayed Pi release, e.g. from the first stereo-specifically attached pre-power-stroke state to a Pi-release state^41–43^. To elucidate if the first state is a pre-power-stroke state and the second state in the 2-step events a post-power-stroke state, we tested the [ATP]-dependence of the lifetimes of the states. We found that raised [ATP] from 5 µM to 50 µM in the presence of Mava, reduced the actomyosin association duration of state-2 but did not affect state-1 (Figure Fig. 4l (fitted data in Fig. S4 and Fig. S5) in both native myosin and single-headed myosin. These results suggest that state-2 in 2-step events likely corresponds to the duration of ‘ADP-bound’ and ‘rigor’ states while state-1 corresponds to a state upstream of the force-generating stroke. Overall, Mava binding to myosin during its interaction with the actin filament increased the lifetime of the pre-power-stroke state, delayed the stroke, and led to two distinct stroke sizes, i.e., short (2-3 nm) in apparent single events and longer (7-10 nm) in 2-step events.

### OM inhibits power-stroke and slows detachment

Next, we probed the effects of OM on the kinetics and mechanics of the actomyosin interactions using either native myosin or single myosin S1 heads at either 5 or 50 µM ATP and 100 µM OM. As depicted in Fig. 5a-d, OM affected both the detachment rate and the stroke size of native cardiac myosin and S1. Consistent with earlier structural and functional studies indicating that OM restricts myosin in the pre-power-stroke state-despite the faster Pi release - the most pronounced effect was observed on the working stroke. While the actomyosin detachment rate was reduced by more than two-fold, the stroke size ranged between 0 and 1 nm (Fig. 5b, d, Fig. S6 and Extended Data Fig. 4e). Unlike with Mava, only single-step binding events were seen with OM. Notably, in the presence of OM, the actomyosin bound lifetime – and thus the actomyosin detachment rates - remained unchanged upon increasing [ATP] from 5 to 50 µM (Fig. 5a, c and Fig. S6), demonstrating [ATP]-independent detachment. This result is consistent with an escape detachment process in the presence of OM characterized by an ATP-insensitive rate constant of 10-20 s^−1^ as observed in our trapping and acto-myosin detachment rate measurements (Fig. 3) and suggested previously^20,30^.

**Fig. 5.**
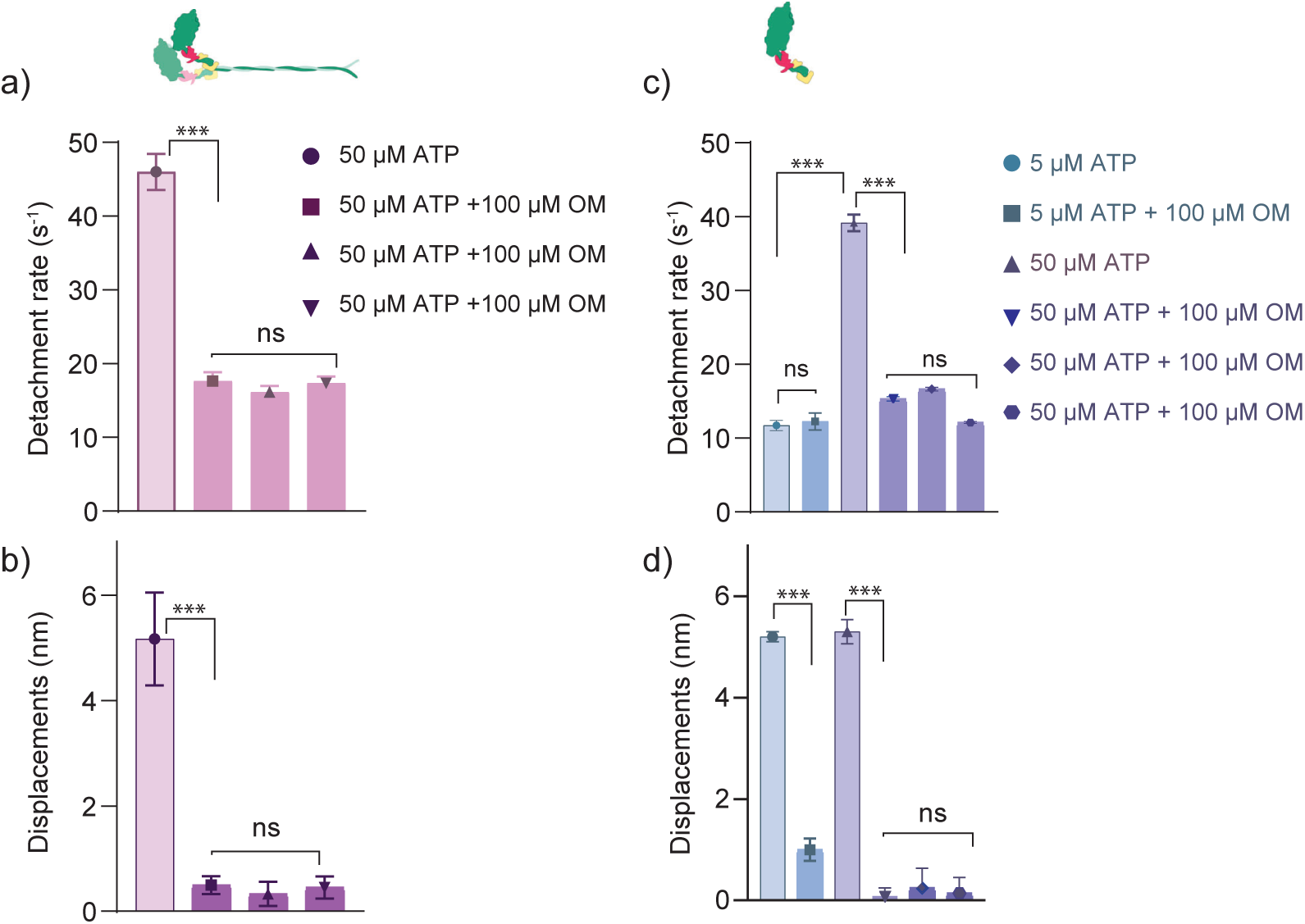
Effects of OM on native beta cardiac myosin and myosin S1 interaction with actin studied using optical trap. **a)** Full-length myosin II interaction with actin in the presence and absence of OM. Bar diagram shows AM detachment rates in the presence of 50 µM ATP and compared with the condition at 100 µM OM. Note that each individual bar in the presence of OM represents results from a single actin bead dumbbell and several myosin molecules. Three actin-bead dumbbells were used to record the interactions between dimeric myosins and actin in the presence of OM. Results from individual actin dumbbells allow us to reliably estimate the stroke size of myosins in the presence of OM, since the actin polarity is determined from the biased stroke size under normal conditions, p <0.0001. p = 0.13 for group comparison between three actin dumbbells for the detachment rate in the presence of OM. **b)** The displacements/stroke size for the corresponding actomyosin binding events indicated in (a), P <0.0001, for group comparison as indicated, p= 0.47. **c)** Myosin subfragment-1 interaction with actin in the presence and absence of OM. Actomyosin detachment rates compared for 5 µM and 50 µM ATP in the presence and absence of 100 µM OM, as indicated. For 50 µM ATP and 100 µM OM condition, results from three individual actin-bead dumbbells are presented, P = 0.37 between 5 µM ATP and 5 µM ATP + 100 µM OM. P<0.0001 between 50 µM ATP and 50 µM ATP + 100 µM OM. P = 0.68 for group comparison **d)** Displacements/stroke size for the corresponding AM interaction events and myosin molecules shown in (c), P <0.0001. P= 0.74 for group comparison. The error bars represent mean ± SD. For comparison between two mean values, Welch’s t-test was used. Group comparisons from three actin dumbbells for both full length myosin and S1 at 50 µM ATP + 100 µM OM using Brown-Forsythe and Welch Anova test. ns – not significant.’

### Modelling of Mava and OM effects

#### Partial myosin inhibition and two cycles

Our data suggest two parallel actomyosin cycles at saturating drug concentrations, one fast cycle (loosely bound drug; similar to physiological cycling) and one slow cycle (strongly bound drug). In our mechanokinetic model (Fig. 6a) the fractional occurrence of each cycle is determined by the values of the drug-binding rate constants (k_Dr+_, k_Dr-_, k_Dr2+_, k_Dr2-_, particularly k_Dr2+_) and the actomyosin attachment rate constant (k_on_(x)) (Fig. S7. S8).

**Fig. 6.**
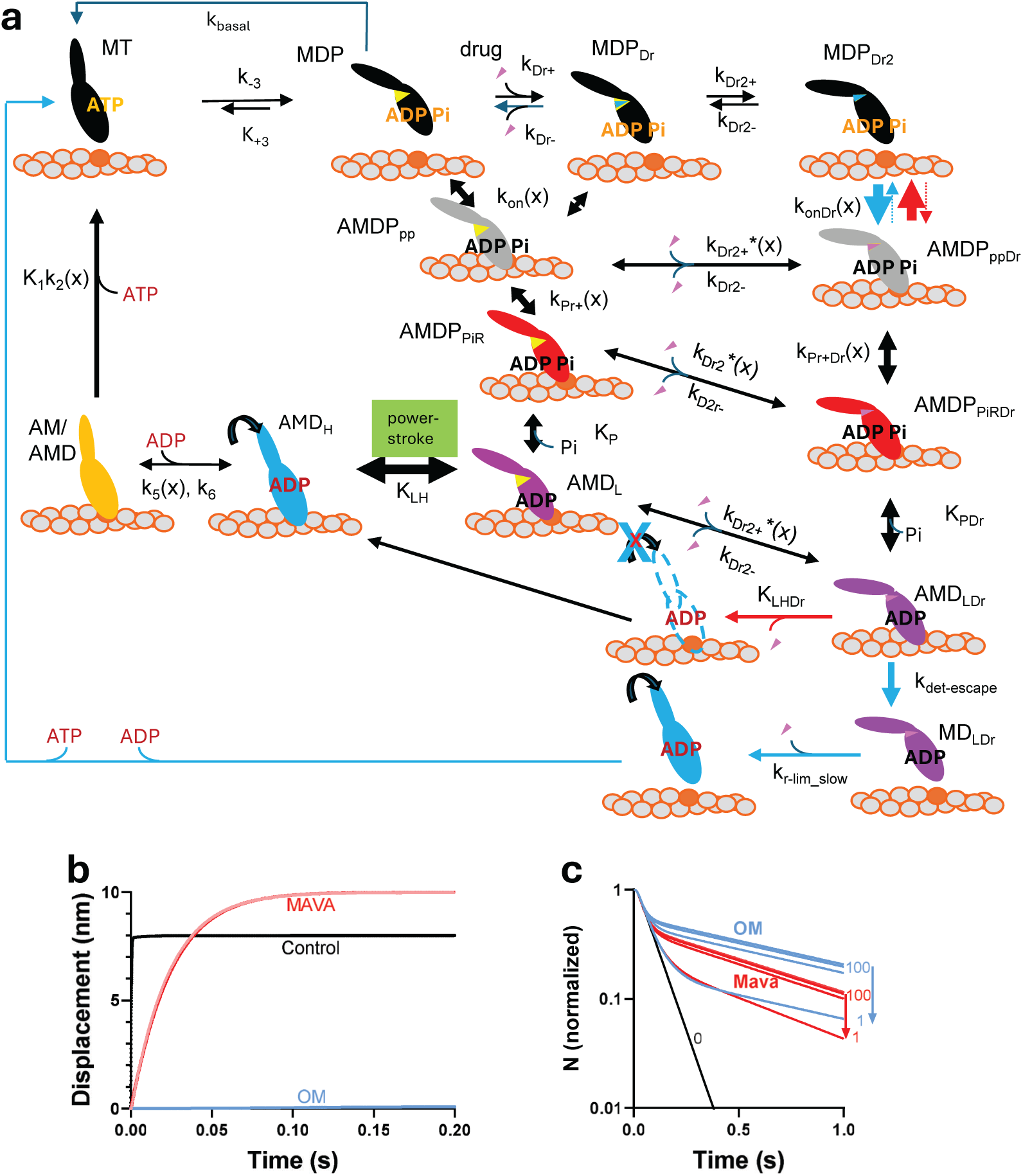
Mechanokinetic modelling. **a.** Model with different states of the myosin head and lever arm orientation (black and colours) denoted by capital letters with or without drug bound (subscript “Dr”). A: actin, M: myosin, D:ADP, P: inorganic phosphate (Pi), T: ATP. Strain-dependent rate constants indicated by argument (x). The sub-scripts “PP” and “PiR” refer to initial pre-power-stroke state and Pi-release state, respectively. The sub-scripts “L” and “H” denote low and high force respectively after Pi-release. The sub-script “Dr” denotes strongly drug-bound states. Other symbols are explained in the text or Tables S3-S4. Thickness of arrows in colour corresponding to drug (Mava: red; OM: blue) indicates increase or decrease by drug of most important transitions (see text). The left cycle (fast) represents both normal physiological cycling and cycling with loosely bound drug, the right cycle (slow) represents cycling with strongly bound drug to myosin with over-primed lever arm. **b.** Effects of drugs on power-stroke simulated by the model as averaged from many single events. Black: without drug. Red: simulated effect of Mava (slow path) assuming a reduction of k_Pr+_’ from 3000 to 40 s^−1^ without other changes. Faint red: simulated effect of Mava (slow path) assuming reduced ΔG_LH_ from 11 to 2 k_B_T and k_LH-_ from 5000 to 18 s^−1^ without other changes. Blue, simulated effect of OM (slow path) assuming a reduction of ΔG_LH_ from 11 to 0 k_B_T and k_LH-_ from 5000 to 0.1 s^−1^ without other changes. **c**. Simulated single molecule ATPase frequency distributions assuming that Mava affects the drug bound path (right) by reducing the attachment rate constant from 22 to 1 s^−1^, increasing the reversal of the attachment rate constant from 5 to 30 s^−1^, and reducing the rate-limiting constant for transition through attached states from 100 (ADP-release) to 40 s^−1^. The parameter values are as in Tables S3-S4 for transitions at the free energy minima (Fig. S10).

#### Pi-release rate, rate limiting steps of slow cycle and in vitro gliding velocity

Mava slows Pi-release and ATP turnover to similar degree^6,17^. Because our trapping data show that the slowest rate constant for transitions between strongly actin-bound states states (pre- to post power-stroke state) is ∼40 s^−1^ the rate limitation of 1-2 s^−1^ (Fig. 2) must reflect reduced actomyosin attachment rate (cf. ^8^). We implement this by reduced attachment rate constant (k_onDr_’) combined with increased rate of attachment reversal (see further SI). These effects also explain the Mava induced reduction of the maximum gliding velocity (Extended Data Fig. 1). The Mava-induced delay of the power-stroke (rate constant ∼40 s^−1^) suggests additional Mava-induced slowing of either of the AMDP_PP_ -> AMDP_PiR_ or the AMDP_L_ -> AMD_H_ transition (power-stroke). The model gives similar predictions for these alternatives (compared to control conditions and OM in Fig. 6b).

To explain an increased Pi-release rate with OM^8,18,19^ we assume increased rate constant for the actin attachment (weak-to-strong transition) in pre-power-stroke myosin states. An almost completely prevented power-stroke^20^ (Fig. 5) requires ATP-independent escape detachment from the AMD_LDr_ state in Fig. 6a (k_det-escape_ ≍ 10 s^−1^) (Fig. 3, 5, ^20,30^). This slow detachment and the inhibited power-stroke explain the OM-induced reduction of the in vitro gliding velocity (Extended Data Fig. 1). The appreciably slower ATP turnover through the slow cycle (1-2 s^−1^) suggests a separate rate-limiting transition, k_r-lim slow_ (Fig. 6a).

#### Multi-exponential frequency distributions of single-molecule actin-activated ATPase

Simulations, based on a simplified kinetic scheme (Fig. S9), predict (Fig. 6c; Fig. S8) double-exponential frequency distributions at saturating drug concentrations with the amplitude of the slow phase (1-2 s^−1^) markedly increased with raised drug concentration at the expense of the fast phase (>10 s^−1^). Importantly, like in experiments (Fig. 2), the amplitude of the fast phase exceeds 50 % at saturating drug concentrations due to a large fraction of the myosin motors cycling through the fast path with loosely bound drug. The lack of drug-induced change in the lag phase (Extended Data Fig. 2) agrees with previous findings^6,8,18,20,21,30,31^ that neither drug affects the ADP-release rate (Fig. S11a). In the simulations, we do not attempt to account for experimental complexities like one very slow phase (rate constant like basal ATPase) and a multiexponential distribution in the absence of drug (see SI and Discussion).

#### Flux through both cycles under steady-state conditions

The substantial flux through the fast cycle in the single-molecule actin-activated ATPase data (Fig. 2) is partly explainable by a competition between strong drug-binding (rate constant k_Dr2+_) and cross-bridge attachment into the normal cycle (rate constant k_on_(x)). However, the model also predicts myosin cycling (20 - 30 %) through a slow non-canonical path (Fig. 6a) at saturating [Mava], under conditions with equilibrium established between all MT and MDP states. This follows from the ratio of the fluxes between the non-canonical and the fast cycle k_onDr_[MDP_Dr2_]/k_on_([MDP]+[MDP_Dr_] (0.845/(0.155 x 20) ≈ 0.27) in the model. Interestingly, this prediction is similar to the flux of about 30 % through the two-step cycle with delayed power-stroke in our trapping data with Mava. With OM, on the other hand, the ratio is 18, suggesting that > 90 % of the myosin molecules would progress through the slow OM bound phase. This is consistent with our optical trapping data (Fig. 5) as well as similar data by others using expressed myosin S1^20^. However, it conflicts with our Fig. 3 and other trapping data using expressed myosin S1^30^ that both suggest an appreciably higher fractional flux through the near normal cycle at saturating OM. One possibility to account for the differences is if myosin heads escape to a greater extent into the fast cycle by partial drug-release from the pre-power-stroke AMD_LDr_ state under some conditions.

#### Model and previous experimental data

In addition to accounting for the observations in our Figs. 1-5, the mechanokinetic model in Fig. 6, with parameter values motivated above and in the Supplementary Information (Tables S3-S4, Fig. S10 for details) also satisfactorily account for OM and Mava effects on steady-state actin-activated ATPase^8,14,27,36^, isometric force and number of attached cross-bridges during isometric contraction (Extended Data Fig. 1, Figs. S12, S13).

### Physiological and pharmacological implications

Small-molecule myotropes, i.e. selective activators and inhibitors of myosin have emerged as promising therapeutics for modulation of the force-generating capacities in different diseases^2–5^. The present study uses a unique combination of optical trapping and recently developed^24^ fluorescence based actomyosin single molecule kinetics methodology to elucidate the mechanisms of two compounds of this type on actively cycling human cardiac myosin cross bridges. In our studies we use modelling to extrapolate the observed effects on actin-activated ATP turnover, myosin stroke size and delayed actomyosin power-stroke to effects of Mava and OM on actin filament gliding speed (Extended Data Fig. 1) and force development in cardiac muscle (Figs. S13). Key findings are summarized in the graphical illustration (Fig. S14).

For S1^E^ as well as full length native β-myosin II, both Mava and OM, at saturating concentrations, produce two parallel turnover cycles in the actin-activated ATPase with a substantial fraction of fast events of similar rate constant (about 10 s^−1^) as under control conditions. In contrast to the ATPase results, the optical trapping data showed more diverse Mava and OM effects. The main Mava effect was two populations of actomyosin binding events resulting in one population with delayed but larger power-stroke size and one with shorter power-stroke size but no apparent delay when compared to the no drug control. For OM, on the other hand, the optical trapping showed complete inhibition of the power-stroke and slowed actomyosin dissociation with a rate constant independent of [ATP], consistent with previous observations^20^. Modelling the effects of the two modulators, suggest that they are consistent with two different actomyosin cycles occurring in parallel at saturating drug concentration. This has been considered for OM^21,30^ previously but, to the best of our knowledge, not for Mava. The behaviour is consistent with partial inhibition of the force-generating cycle in analogy to the action of partial agonists/antagonists in a two-state model of receptor activation ^44,45^. The existence of two parallel cycles at saturating Mava is evident both from our actomyosin ATPase assay (Fig. 2) and trapping data (Fig. 4). Two cycles with OM are suggested both by the actin-activated ATPase data (Fig. 2) and the fluorescence-based actin-myosin binding assay (Fig. 3).

The effects of Mava and OM as well as normal cardiac regulation have increasingly been associated with modulation of the proportion of myosin in the super-relaxed (SRX) state, associated with low ATP turnover relative to disordered relaxed (DRX) state^34,46–50^. Particularly, the effect of Mava has primarily been attributed to an increased fraction of myosin heads in a folded back SRX state^34,50^. In partial contrast to this view, but seemingly consistent with recent findings^51,52^, our results show a range of striking modulatory effects of both Mava and OM on the single motor level, independent of inter-head interactions. Our mechanokinetic modelling suggests that these effects would extrapolate to the type of effects on force and shortening velocity observed in cardiomyocytes.

Both ATPase results (Fig. 2) and the trapping data (Fig. 4) showed two populations of interaction events with Mava. One of these constituted a two-step actomyosin interaction with a delayed power-stroke and longer stroke size of 7.5 - 10 nm compared to 4-5 nm without Mava. The 9° over-priming of the lever arm in the pre-power-stroke state found in structural studies^16^ when Mava occupies the pocket between the upper 50 kD domain and the converter region provides a plausible structural basis for the increased stroke length. Crucially, the latter stroke size could be directly measured due to the delay with a pre-power-stroke state of prolonged life-time that eliminates the need for estimation via histogram-shift analyses or ensemble averaging. A mechanistic basis for a shorter stroke size (2-3 nm) of Mava-associated one-step events is less evident from structural analysis. It seems likely that these events are associated with the fast process in the actin activated ATPase (Fig. 2). The latter is less affected by Mava (turn-over rate constant close to 10 s^−1^) and shows a fractional amplitude > 50 % at saturating concentration, like the flux through the one-step cycle in the trapping data. One possibility is that a weak-binding state of Mava, reduces the power-stroke amplitude with minimal effects on turnover kinetics. This would be consistent with the left cycle in the scheme in Fig. 6a if a substantial sub-population of weakly bound drug binds in the pocket with allosteric effects to reduce the power-stroke size but without appreciable effects on the kinetics of ATP turnover. The existence of different Mava binding states is consistent with molecular modelling^14^.

An intriguing aspect of our work is that it implies a mechanism for fine-tuning of cardiac contraction on a lower hierarchical level than interacting heads, by shuttling myosin through different actomyosin cycles to varying degree. This mechanism is of interest to explore in future drug discovery to obtain fine-tuned effects. Whether the mechanism has a physiological role e.g. mediated by post-translational modifications or endogenous ligands is unknown. In this context it is of interest to note, however, that the pocket is primarily accessible in certain functional states with excessive over-priming of the lever arm. It is also of interest to note that a non-drug dietary fatty acid^15^ exerts functional effects by binding to the Mava/OM pocket and that post-translational modifications have been noted in or near the pocket in slow human skeletal muscle myosin (identical sequence as cardiac myosin)^53^. Because our single molecule ATPase method directly quantifies the fractional contribution of the two cycles, it would be ideal for elucidating the issue further whether in drug screening processes or basic functional studies, facilitated by the need for only ng proteins for each assay.

### Conclusions

In this work, ATP turnover investigations and single-molecule optical trapping studies combined with mechanokinetic modelling provided new insights into the mechanism of Mava and OM modulation of the cardiac myosin chemo-mechanical cycle. Facilitated by the capacity of single molecule methodology to detect heterogeneities, our findings reveal previously unknown functional effects that may contribute to drug-induced modulations also within highly organized and integrated environment of the cardiac systems. The results underscore the importance of in-depth biochemical and biophysical studies on the single motor level for developing effective therapeutic interventions for cardiac diseases. Adding to their translational relevance, such studies advance our fundamental understanding of cardiac physiology with broader implications for understanding how cardiac performance is modulated under both physiological and pathophysiological conditions.

## Materials and Methods

### Materials and chemicals

The zero-length crosslinker 1-ethyl-3-(3-dimethylaminopropyl) carbodiimide hydrochloride (EDC) was obtained from Thermo Scientific (cat. no. 22980) and 2-(N-Morpholino) ethanesulfonic acid hydrate, MES Hydrate was from Sigma-Aldrich, (cat no. M8250). The following assay solution components were from Sigma-Aldrich: cyclooctatetraene (COT; cat. no. 138924), 4-nitrobenzyl alcohol (NBA; cat. no. N12821), pyranose oxidase (POX; cat. no. P4234), trolox (cat. no. 238813), creatine phosphate (CP; cat. no. P7936) and Creatine phosphokinase (CPK; cat.no. C3755). High purity bovine serum albumin (BSA; cat. no. A0281) and Phalloidin (cat. No 17466-45-4) were also from Sigma Aldrich whereas rhodamine phalloidin for actin filament fluorescence labeling was from Invitrogen (cat. No. R415). Mavacamten (MYK-461) and Omecamtiv mecarbil (CK-1827452) were purchased from Selleck Chemicals. All other chemicals were of analytical grade and obtained from Thermo Fisher or Sigma-Aldrich.

### Expression and purification of myosin subfragment in C2C12 cells

The myosin subfragment 1 (S1^E^) expression and purification were performed as described previously^24,54^. The expression plasmid consists of pcDNA 3.1 vector backbone that is fused to the MYH7 gene encoding truncated human β-cardiac myosin (1-848 amino acids) denoted as S1^E^ here, followed by TEV protease cleavage site, enhanced Green Fluorescent protein (eGFP) and FLAG tag (GenScript Biotech Corporation). The plasmid was transfected into ∼95% confluent C2C12 cells (ATCC CRL 1772) using a non-viral transfection method with JetPrime^®^ DNA transfection reagent kit (Polyplus-transfection S.A, Illkirch, France). The transfected cells were differentiated for 7 days, and the expressed myosin was isolated using affinity purification. The concentration of the purified S1^E^ was estimated using a Tecan Spark microplate reader with eGFP (MyBioSource, cat. No. MBS146512) as standard. Fluorescence measurements were performed at an excitation wavelength of 485/20 nm and emission at 535/20 nm.

### Human myocardium samples

Human myocardial sample from non-diseased, non-transplanted heart muscle tissue or donor samples were obtained from Sydney heart bank, Australia. Local ethical approval was obtained from Hannover Medical School ethics committees for use of tissue samples (Reference No. 10859_BO_K_2023). Post-death, harvested heart tissue samples were immediately flash-frozen in liquid nitrogen. The storage procedure was adjusted to minimize the influence of tissue handling on the protein quality and to preserve the native function of the biomolecules. In this study, cardiac left ventricular or interventricular septum tissues from three different donors, including a 42-year-old female and two 56-year-old males were used.

### Native dimeric myosin extraction from human cardiac tissue

Full-length human ventricular myosin was extracted from non-transplanted donor hearts as described before^55^. Briefly, tissue was mechanically ground into small pieces in liquid-nitrogen-cooled mortar and pestle and subsequently collected to an ultracentrifuge tube containing 1:3 (w/v) ratio of extraction buffer (500 mM NaCl, 10 mM HEPES pH 7.0, 5 mM MgCl_2_, 2.5 mM MgATP^1^, supplemented with 2 mM DTT and 1 mM AEBSF). After a 20 min extraction time, the sample was centrifuged to remove the debris (TLA 120.2, 150,000g, 60 min, 4°C). The resulting supernatant was diluted tenfold in ultrapure water containing 2 mM DTT and incubated for 40 min on ice, to promote myosin filament formation under low salt conditions. Myosin was collected by subsequent centrifugation (TLA 110, 70,000g, 30 min, 4°C) and resuspended in extraction buffer supplemented with 2 mM DTT and 0.5 mM AEBSF. The concentration of myosin was determined using the Bradford assay. Isolated myosin was aliquoted, flash frozen in liquid nitrogen and stored at −80°C in 50% glycerol. Cardiac tissue samples employed in this procedure were from human left ventricular tissue.

### Preparation of actin filaments for single molecule fluorescence experiments

Actin was purified from rabbit fast skeletal muscle as described previously^23,56^. The rabbit was euthanized according to the procedure approved by the Regional Ethical Committee for Animal Experiments in Linköping, Sweden (ref. 17088-2020). The rabbit was first anesthetized by an intramuscular injection of 0.25 ml Zoletil (active substances: Zolazepam, 6 mg/kg; Tiletamin, 6 mg/kg and Medetomidin, 0.6 mg/kg) followed by 2 ml of penthobarbital (100 mg/ml) in an ear vein. Euthanasia was performed by the Linnaeus University veterinarian. The actin filaments were isolated from the muscles of a single rabbit and were solely used for methodological purposes. In vivo experiments were not performed on the live animals. The laboratory animal use was in accordance with Animal Welfare act (Swedish law: SFS: 2018:1192) and the guidelines of Swedish Board of Agriculture. These regulations comply with EU-Directive 2010/63/EU on the use of animals for scientific purposes.

### Preparation of actin filaments for optical trapping studies and in vitro motility assays

Actin was purified from chicken pectoralis muscle as described^56^. To obtain sufficiently long biotinylated actin filaments (≥ 20 µm) for optical trapping experiments, chicken G actin and biotinylated G actin were mixed in equimolar ratios to a final concentration of 0.1 µg/µl each in polymerization-buffer (5 mM Na-phosphate, 50 mM K-acetate, and 2 mM Mg-acetate, 0.02% Na-Azide), containing 1 mM DTT, 1 mM ATP, and 0.5 mM AEBSF protease inhibitor (Cat No. 30827-99-7, PanReac Applichem ITW). The mixture was incubated overnight at 4°C and followed by addition of equimolar concentration of fluorescent (TMR, tertramethylrhodamine) phalloidin (cat no. P1951, Sigma-Aldrich) and biotin phalloidin (0.23 nM, Invitrogen/Thermofischer Scientific, B7474) to label the actin filaments. For *in vitro* motility assay, the unlabelled F-actin was polymerized using polymerization buffer containing 0.5 mM AEBSF for minimum 1 h on ice, while the labelled F-actin additionally contained fluorescent phalloidin.

### In vitro motility assays

Motility assay was performed as before^55^, with minor changes. Myosin was deposited on BSA-coated glass chambers for 5 min. After blocking non-functional myosins using unlabelled actin, Rhodamine-Phalloidin (Sigma-Aldrich; P1951) labeled actin was injected into the chamber and incubated for 1 min. Assay buffer (25 mM imidazole pH 7.2, 25 mM NaCl, 4 mM MgCl_2_, 1 mM EGTA, containing 2 mM DTT and 1 mM AEBSF, 0.5% methylcellulose, anti-bleaching system (10 mM DTT, 18 μg/ml catalase, 0.08 mg/ml glucose oxidase, 10 mg/ml D-Glucose), and the desired drug concentration was injected into the chamber before recording. Mavacamten (Hycultec, HY-109037) and Omecamtive Mecarbil (MedChemExpress, HY-14233) were dissolved in dimethyl sulfoxide (DMSO) at 10 mM concentration, briefly sonicated and stored at −80°C until used. In each chamber, 2-3 drug conditions were tested, starting with low and followed by higher drug concentrations. Movies were recorded using a custom-made TIRF microscope, with 158 nm pixel size and 0.2 s time resolution. MTrackJ plugin^57^ for ImageJ was used to manually track the filaments. Mean filament speeds were pooled for each condition to generate the histograms, which were fitted with Gaussian curves to obtain the mean ± SD values. Dose-response curves were made using the mean ± SD values and fitted using the “[Inhibitor] vs. response - Variable slope (four parameters)” model in Prism v9.5.1.

### TIRF-microscopy

Total internal reflection fluorescence (TIRF) microscopy was used for observation of S1^E^ with eGFP by excitation using a 488 nm laser 70 mW (experiment at 9.5-19 mW intensity) with a FITC filter cube (525 nm with 50 nm bandwidth). Alexa 647-ATP was excited using a 640 nm laser, 50 mW (experiment at 30-33 mW) with a Cy5 filter cube (706 nm with 95 nm bandwidth). Rhodamine phalloidin, finally, was excited using a 561 nm laser, 60 mW (experiment at 4.5 mW) with a Cy3 filter cube (595 nm with 44 nm bandwidth). All lasers were incorporated into Nikon LUNF laser unit. The lasers were coupled to the iLAS2 TIRF module with fast galvanometers to spin the laser in the back focal plane of the objective to average out interference artifacts and achieve uniform illumination. Microscope specification: Nikon Ti2-E coupled to a 60x oil immersion objective (NA 1.49). The videos were recorded using an electron-multiplication charged coupled device camera Andor iXon Ultra 897 controlled by NIS Elements software (Nikon, ver. 60.10.02) with gain set at 100 and exposure time at 30 ms).

### Cross-linking of F-actin to nitrocellulose surface

A volume of 10 µl of 10 nM Rhodamine-phalloidin or plain phalloidin labeled F-actin with actin prepared from rabbit skeletal muscle as described previously^23,56^ was crosslinked with 15 mM EDC in 50 mM MES pH 6.5 for 1 minute^24^. After crosslinking, the reaction was quenched by washing 3 times with either low ionic strength solution (LISS) (10 mM MOPS, 1 mM MgCl_2_, 0.1 mM K_2_EGTA, 1 mM DTT) or X-linking wash buffer (10 mM Imidazole, 5 mM KCl, 3 mM MgCl_2_, 1 mM DTT pH 7.5). Exposed nitrocellulose surface was blocked by addition of BSA solution (1mg/ml) in LISS or X-linking wash buffer.

### EDC crosslinking of myosin sub-fragment 1 to surface-immobilized actin filaments

ATP removal from the purified S1^E^ is crucial for effective rigor binding to F-actin before applying EDC cross-linking. To that end we dialyzed the purified S1^E^ in a Harvard apparatus Fast Spin Dialyzer (chamber volume 200 µl) with regenerated cellulose membrane with molecular weight cut-off of 10 kDa towards cross-linking (X-linking) wash buffer, with addition of BSA (1 mg/ml; final concentration). A volume of 5 µl of the single headed S1^E^ (≤12 nM) was then added to the surface-attached filaments. The attachment of the myosin construct to F-actin was followed live in the TIRF microscope by observing the fused eGFP. Subsequently, EDC cross-linking was initiated by incubation with 15 mM EDC in 50 mM MES pH 6.5 for 10 min following a recently developed procedure^24^. The reaction was quenched by three washes with X-linking wash buffer. Myosin that had not been crosslinked was washed away with 50 nM non-fluorescent ATP solution, containing a final concentration of 1mg/ml BSA in X-linking wash buffer. This process was followed by 2 washes with X-linking wash buffer and subsequently followed by addition of assay solution. The recording of data was initiated after 3 minutes.

### EDC crosslinking of full-length myosin to surface-immobilized actin filaments

The βM-II sample was diluted to a final volume of 200 µl (final concentration 33 nM) in high ionic strength solution (HISS) (300 mM KCl, 10 mM MOPS, 1 mM MgCl_2_, 0.1 mM K_2_EGTA, 1 mM DTT, pH 7.4) with addition of final concentration of 1mg/ml BSA. To remove ATP from previous purification steps the βM-II sample was dialyzed towards HISS in a Harvard apparatus Fast Spin Dialyzer as described for S1^E^ above. The sample was dialyzed towards the following volumes: 3x 125 ml for 1 hour each, 1x 500 ml overnight and finally 125 ml in the morning for 1hour before the start of assay. A volume of 5 µl of full-length myosin with a concentration of 3.3 nM in HISS was added to the surface-attached actin filaments. The attachment was followed by one wash with LISS. The subsequent crosslinking of full-length myosin (βM-II) to F-actin was performed essentially as with S1^E^. That is, first we incubated with 15 mM EDC in 50 mM MES (pH 6.5) for 10 min. The reaction was then quenched by three washes with HISS. Myosin that had not been crosslinked was washed away with 50 nM non-fluorescent ATP solution containing a final concentration of 1mg/ml BSA in HISS. To ensure satisfactory removal of any remaining unbound βM-II, we also applied 2 further washes with HISS followed by one wash with LISS. Assay solution was then infused, and the recording was initiated after 3 minutes.

### Single molecule fluorescence-based ATPase assay solution conditions

The optimized assay solution is based on that in ^23^, developed to minimize photobleaching and photo-blinking issues associated with the use of Alexa 647 ATP^58^. Briefly, the assay solution with 45 mM KCl, 10 mM DTT, 10 mM MOPS, 0.64% methylcellulose and Alexa 647-ATP (final concentration 5 nM) also contain an oxygen scavenger mixture (7.2 mg/mL glucose, 3 U/mL Pyranose oxidase, 0.01 mg/mL catalase), an ATP regenerating system (2.5 mM creatine phosphate, 0.2 mg/mL Creatine phosphokinase) and an anti-blinking/bleaching cocktail (2 mM COT, 2 mM NBA, ∼2 mM Trolox/Trolox-Quinone). Addition of 500 nM unlabeled ATP was included under single molecule actin activated ATPase studies to allow distinction of individual events despite close spacing of cross-linked myosin motor domains along the actin filaments^24^.

### Single molecule fluorescence-based binding assay

In another fluorescence based single molecule assay we did not cross-link the myosin motor fragments to actin filaments but only cross-linked the filaments to the nitrocellulose surface. We added 5-10 nM (final concentration) of single headed myosin construct (S1^E^) fused to eGFP, diluted in a binding-assay buffer. The latter contained 5 mM KCl, 10 mM Imidazole (pH 7.5), 3 mM MgCl_2_, 1 mM DTT and varying ATP concentrations (100-1000nM) supplemented with an oxygen scavenger and an ATP-regeneration cocktail (concentrations similar to what is described in the ATPase assay solution). Finally, 30 µM Mava or 100 µM OM (final concentration) or corresponding concentrations of DMSO (<1.4%) were included as desired. A volume of 10 µl of the solution was added to a chamber with surface attached actin filaments (as described previously). The binding under various ATP and drug conditions were recorded for 3 minutes at a temperature of 23 ± 1°C.

### Image processing and data extraction from single molecule recordings

After recording the videos were converted to 8-bit followed by background subtraction via rolling ball algorithm (rolling ball radius 5-pixels). Time-projections of the recorded videos are created either via standard deviation (SD) or average projection of all frames in a video. The following procedures are done in Image J (Fiji). For Alexa 647-ATP turnover experiments regions to analyze are picked based on observation of S1^E^ myosin positioned along actin filament (average projection based on eGFP) together with overlapping corresponding SD projection of Alexa 647-ATP. For β-MII, regions were picked based on Rhodamine phalloidin actin filament position overlapping with Alexa 647-ATP projection (SD). Detailed criteria for analysis of actin-activated ATP turnover is described further in ^59^.

### Data analysis in single molecule fluorescence studies

The dwell-on time from fluorescent myosin/Alexa 647-ATP binding events in a recorded time series were analyzed manually. In short, a dwell-on time was defined as the time from a one-step increase of intensity above a defined threshold until single-step drop in intensity back to baseline as described in greater detail previously^23,24^. Prior to fitting exponential functions to the data, the dwell-time frequency distribution was shifted by ∼ 30 ms to eliminate the time lag. The fitting was achieved using non-linear regression implemented in GraphPad prism Version 10.2.2 (Marquardt-Levenberg algorithm) giving R^2^ values of 0.993-0.998 for actin activated and 0.995-0.998 for basal ATPase activity. This non-linear regression based approach for fitting triple exponential functions to actin-activated ATPase data gave very similar rate constants and amplitudes of the exponential processes as when an algorithm based on Maximum Likelihood Estimation (MEMLET)^60^ was used in control analyses.

### Three-bead assay using optical trap

The optical trapping set up was described in detail previously^61,62^. For the assay, flow cells with approximately 15 µl chamber volumes were assembled using coverslips with nitrocellulose-coated beads. Glass microspheres (1.5 µm) suspended in 0.05 % nitrocellulose in amyl acetate were applied to 18×18 mm coverslips. All the dilutions of biotin-actin filaments were made in reaction buffer (KS buffer) containing 25 mM KCl, 25 mM HEPES (pH 7.4), 4 mM MgCl_2_, and 1 mM DTT. The full-length native myosin was diluted in high salt extraction buffer without MgATP. For the experiment, the chamber was prepared as follows, 1) flow cells were first incubated with 1 µg/ml native myosin for 1 min, 2) washed with high salt extraction buffer without ATP and thereafter with KS buffer, 3) followed by wash with 1 mg/ml BSA and incubated further for 2 min to block the surface, 4) finally, reaction mixture containing 0.8 µm neutravidin coated polystyrene beads (Polyscience, USA) and 1-2 nM biotinylated actin was flowed in with 5 µMATP (or with 50 µM ATP), ATP regenerating system (10 mM creatine phosphate, and 0.01 unit creatine kinase) and deoxygenating system (0.2 mg/ml catalase, 0.8 mg/ml glucose oxidase, 2 mg/ml glucose, and 20 mM DTT). The intended concentrations of Mava or OM was added to the chamber together with the reaction mixture. The assembled flow chamber was sealed with silicon and placed on an inverted microscope for imaging and trapping assay. An actin filament was suspended in between the two laser trapped beads (Fig. 4a), pre-stretched, and brought in contact with the platform bead immobilized on the chamber surface. Low-compliance links between the trapped beads and the filament were adjusted to about 0.2 pN/nm or higher^63^. The bead positions were precisely detected with two 4-quadrant photodetectors (QD), recorded and analysed. The acto-myosin interaction events were monitored as a reduction in free Brownian noise of the two trapped beads. Data traces were collected at a sampling rate of 10,000 Hz and low-pass filtered at 5000 Hz. All the experiments were carried out at room temperature of approximately 22°C.

### Analysis of optical trapping data

Data records were analyzed for acto-myosin interaction events by using the running variance and threshold method as described earlier^64,35^. This method enabled the actomyosin bound states (characterized by low variance in displacement) to be distinguished from the unbound states (high variance). Custom MATLAB routines were implemented to analyse the data records for ‘actomyosin’ interaction lifetime, ‘*t_on_*’ and to estimate the displacements during individual interactions as described earlier^65^.

### Single myosin molecule interaction with actin filaments

To ensure the reliability of single-molecule optical trapping data, rigorous quality control procedures were implemented to confirm that each recorded trace originated from an intermittent interaction between a single myosin molecule and an actin filament. Key criteria were applied to minimize the inclusion of multi-motor events: 1) Myosin density on the bead surface was carefully titrated by diluting the myosin solution to achieve low occupancy; 2) During bead screening, only approximately one out of 8-10 myosin-coated beads exhibited detectable interaction with the actin filament, indicating low occupancy; 3) Only traces displaying well-resolved, discrete actomyosin binding events –characteristic of single-molecule interactions - were included in the analysis; 4) Traces exhibiting closely-spaced or stepwise binding events – suggestive of multiple motors interacting simultaneously or consecutively with the dumbbell - were excluded. Based on Poisson statistics and the observed fraction of motor-free beads, the estimated probability of more than 1 motor per bead was about 5 %. After excluding traces with multi-event and or closely spaced events, the effective likelihood of multiple motors per bead was less than 5 %.

### Ensemble average analysis of optical trapping data

Ensemble averaging was used to estimate the first (δ1) and second power-stroke (δ2) during each individual actomyosin crossbridge cycle. The computational analysis was performed using tool SPASM (Software for Precise Analysis of Single Molecules) developed by Blackwell et al.^66^. To select the AM-binding events suitable for this analysis, following criteria were applied: (1) The displacement over time trace from one of the two trapped beads exhibited a high variance ratio (>4) between the bound and unbound state/free dumbbell noise; (2) Individual binding events were well separated in time, consistent with discrete single myosin-actin interactions; (3) The binding events – particularly the long-duration one-showed no evidence of nonspecific interference signal in either of the bead channels (4) the direction of the bead motion induced by actomyosin interaction should be synchronized across both channels; and (5) The channel with the higher signal-to-noise ratio was selected for analysis from myosin molecule

## Acknowledgements

Funding is acknowledged from The Swedish Research Council (grant # 2023-03453), The Crafoord Foundation (grant # 20240004) and the Linnaeus University, Faculty for Health and Life Sciences. This research was supported by a grant from Deutsche Forschungsgemeinschaft (DFG, German Research Foundation) to MA, Project number: 530881940 (AM 507/2-1).

## Author contributions

AM, MA, ST and AEB conceived the project. AM and MA supervised the project. ES performed and analysed in vitro motility assay experiments. TW performed and analysed optical tweezers experiments, AEB performed and analysed single molecule fluorescence experiments. LPV and AEB expressed and purified myosin fragments in C2C12 cells. AM and MA conducted the safety, security and ethics investigation. CDR provided the donor heart tissue samples. AN, MN and TK provided critical inputs to the conception and design of the study. AM, MA, AEB, LPV, ES and TW wrote the parts of first draft of the manuscript. AM and MA edited and finalized the manuscript together with all co-authors and all authors approved the final draft of the manuscript.

## Generative AI statement

The author declares that no Generative AI was used in the creation of this manuscript.

## Data Availability Statement

Most data supporting the findings of this study are available within the article, and the supplementary information files. Any additional data are available from the corresponding author upon reasonable request.

^1^When “ATP” is used as an abbreviation for Adenosine 5’-triphosphate in the following and above, it is always assumed to be complexed with Mg^2+^ because the solutions contain excess free Mg^2+^.

---

**Extended Data Fig. 1.**
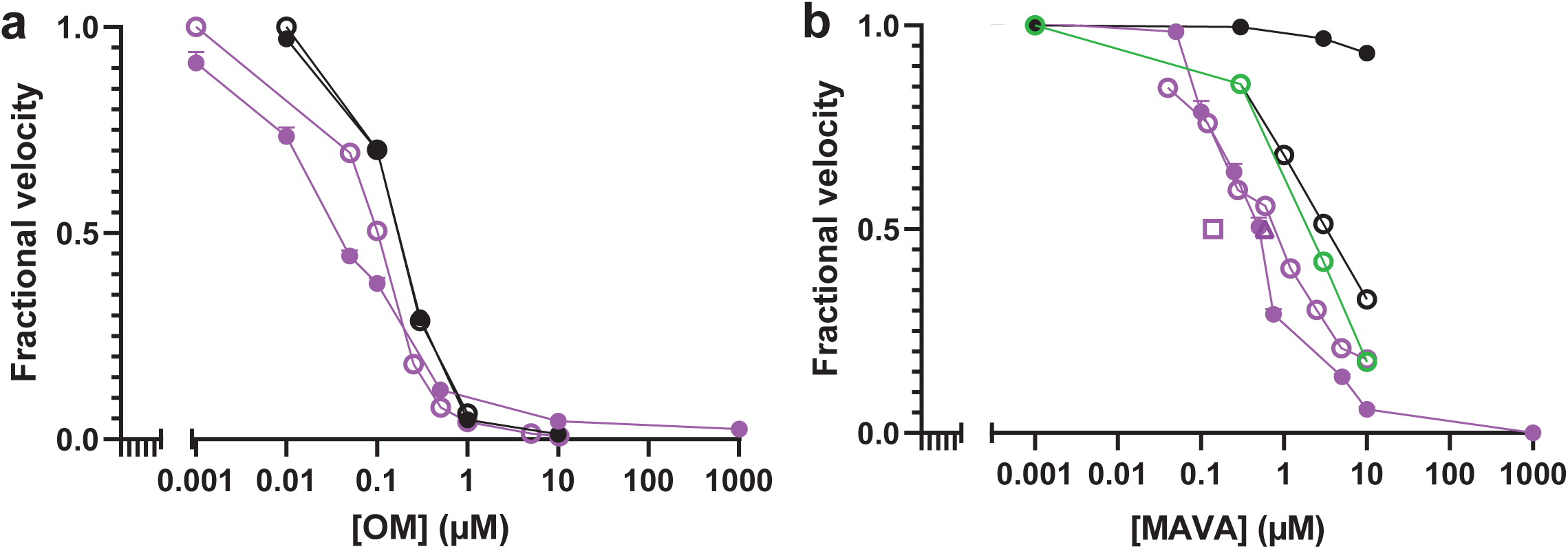
Experimental velocity data from main Fig. 1 (filled purple circles) compared to experimental data by others (open purple circles)^17,21^ and simulated data (black) based on mechanokinetic implementation of model in Fig. 6a. **a.** OM. Filled black symbols assume shortening against zero load. Open black symbols assume shortening against load corresponding to 2 % of the maximum isometric force in the absence of drug. **b.** Mava. Meaning of black symbols as in panel a. The effect of the load was somewhat greater with Mava if the drug-affinity was higher in attached pre-power-stroke states than in the detached states (c_att_=5 (green) instead of c_att_=1 (black)). Results with Mava for cardiac heavy meromyosin expressed in C2C12 cells with murine light chains^16^ (purple square) and expressed subfragment 1 with cardiac light chains^16^ (purple triangle) are also indicated by the IC_50_-values. All data simulated using parameter values in Tables S3-S4. Note, that the earlier cardiac sub-fragment 1 data for Mava (with cardiac light chains)^16^ and our Mava data for native myosin (Fig. 1) are reasonably well fitted on the assumption of a loaded motility assay with a load (due to non-functional “dead” heads always present and surface interactions) corresponding to 2 % of the isometric force. No similar effects were found with OM. The results are consistent with a true reduction in the maximum velocity of shortening against zero load by OM but a reduction in loaded velocity with Mava due to reduced force-generating capacity. The latter finding is supported by lack of apparent change in shortening velocity upon addition of Mava (0.5 µM) to rodent permeabilized cardiac myocytes^67^.

**Extended Data Fig. 2.**
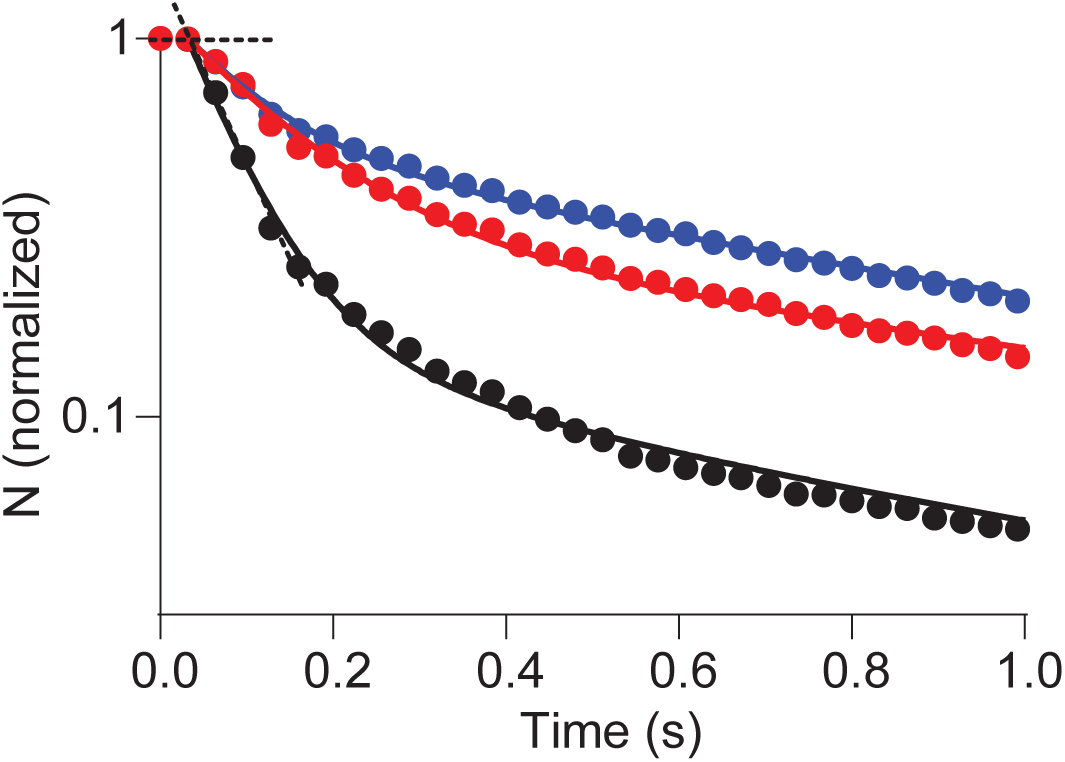
Frequency distributions in Fig. 2f for S1^E^ under control conditions and in the presence of Mava (red) and OM (blue) replotted to only show events with short dwell-times. Curves: Same triple exponential functions fitted to the data as in Fig. 2f after appropriate shifts to accommodate the time lag, estimated from the intersection of the dashed lines.

**Extended Data Fig. 3.**
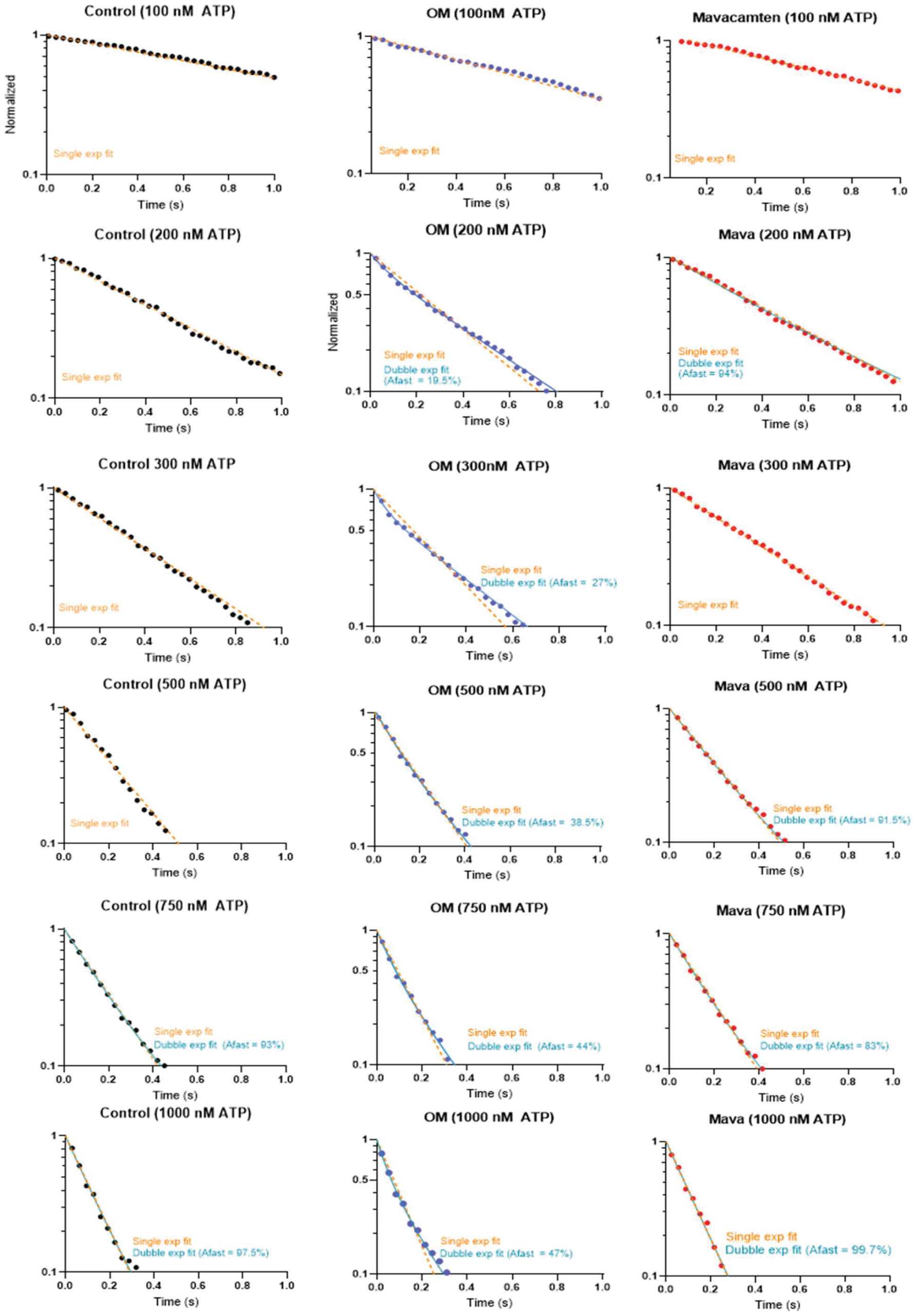
Cumulative dwell time distribution of for myosin binding to actin based on eGFP fluorescence terminated by ATP induced detachment at different ATP concentrations with either OM or Mava. In most cases the data are well described by a single exponential fit for both the control (black graph) and Mava (red graph) as suggested by AIC-values. In the control and Mava cases, where a double exponential is preferred, one process was greatly dominating (83-99%). Based on comparison of AICc values derived from the fit to OM data, the double-exponential model is preferred at all ATP concentrations with consistently more similar amplitudes of the fast and slow process. The n_dwells_ for each condition can be seen in supplementary **Table S2.**

**Extended Data Fig. 4.**
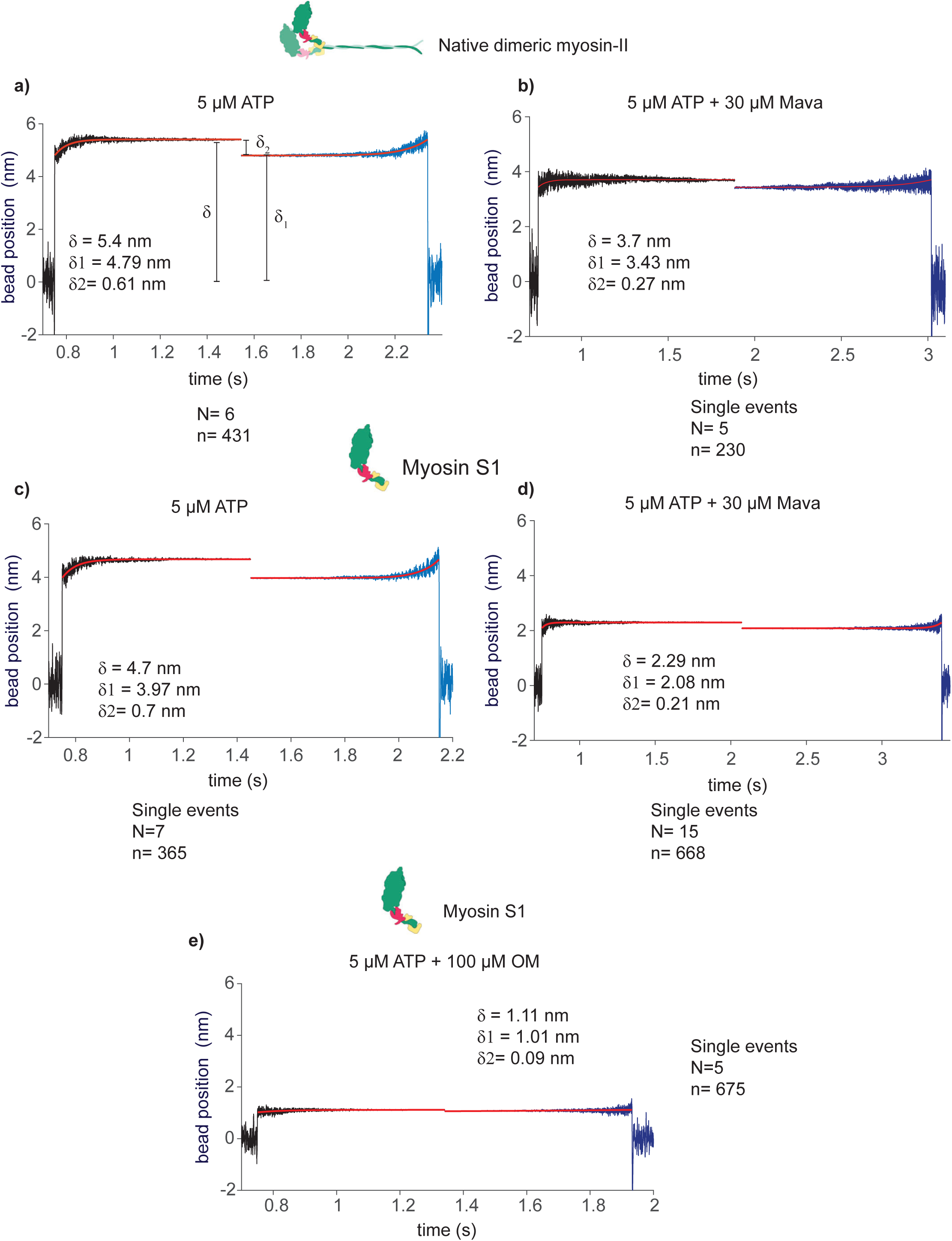
Ensemble average analysis of optical trapping data. The apparent single binding events were analysed. Actomyosin interactions were synchronized at the beginning and end of the events. Ensemble averaging used to estimate the first and second powerstroke size (δ_1_ and δ_2_) as indicated. a) Native full-length myosin, ensemble average of binding events at 5 µM ATP, N = 6, n=431. b) Ensemble average of single actomyosin interaction events at 5 µM ATP and 30 µM ATP, N = 5, n = 230. c and d) Myosin S1 interaction with actin at 5 µM ATP, N = 7, n=365 (c), and in the presence of 30 µM ATP (d), N = 15, n=668. e) Myosin S1interaction events with actin at 5 µM ATP and 100 µM OM analysed. N = 5, n = 675. N= Number of myosin molecules, n= number of binding events.

## References

1 Rassier, D. E. & Mansson, A. Mechanisms of myosin II force generation: insights from novel experimental techniques and approaches. Physiol Rev 105, 1–93, doi:10.1152/physrev.00014.2023 (2025).

2 Spudich, J. A. From amoeboid myosin to unique targeted medicines for a genetic cardiac disease. Front Physiol 15, 1496569, doi:10.3389/fphys.2024.1496569 (2024).

3 Trivedi, D. V., Nag, S., Spudich, A., Ruppel, K. M. & Spudich, J. A. The Myosin Family of Mechanoenzymes: From Mechanisms to Therapeutic Approaches. Annu Rev Biochem 89, 667–693, doi:10.1146/annurev-biochem-011520-105234 (2020).

4 Gyimesi, M. et al. Single Residue Variation in Skeletal Muscle Myosin Enables Direct and Selective Drug Targeting for Spasticity and Muscle Stiffness. Cell 183, 335–346 e313, doi:10.1016/j.cell.2020.08.050 (2020).

5 Miller, C. A., Quinones-Hinojosa, A. & Rosenfeld, S. S. Non-muscle myosin II is a promising therapeutic target. Trends Pharmacol Sci 46, 931–934, doi:10.1016/j.tips.2025.08.011 (2025).

6 Green, E. M. et al. A small-molecule inhibitor of sarcomere contractility suppresses hypertrophic cardiomyopathy in mice. Science 351, 617–621, doi:10.1126/science.aad3456 (2016).

7 Olivotto, I. et al. Mavacamten for treatment of symptomatic obstructive hypertrophic cardiomyopathy (EXPLORER-HCM): a randomised, double-blind, placebo-controlled, phase 3 trial. Lancet 396, 759–769, doi:10.1016/S0140-6736(20)31792-X (2020).

8 Malik, F. I. et al. Cardiac myosin activation: a potential therapeutic approach for systolic heart failure. Science 331, 1439–1443, doi:10.1126/science.1200113 (2011).

9 Cleland, J. G. et al. The effects of the cardiac myosin activator, omecamtiv mecarbil, on cardiac function in systolic heart failure: a double-blind, placebo-controlled, crossover, dose-ranging phase 2 trial. Lancet 378, 676–683, doi:10.1016/S0140-6736(11)61126-4 (2011).

10 Savsin, H. & Tokarek, T. Comprehensive Review: Mavacamten and Aficamten in Hypertrophic Cardiomyopathy. Biomedicines 13, doi:10.3390/biomedicines13071619 (2025).

11 Chuang, C. et al. Discovery of Aficamten (CK-274), a Next-Generation Cardiac Myosin Inhibitor for the Treatment of Hypertrophic Cardiomyopathy. J Med Chem 64, 14142–14152, doi:10.1021/acs.jmedchem.1c01290 (2021).

12 Shen, S., Sewanan, L. R., Jacoby, D. L. & Campbell, S. G. Danicamtiv Enhances Systolic Function and Frank-Starling Behavior at Minimal Diastolic Cost in Engineered Human Myocardium. J Am Heart Assoc 10, e020860, doi:10.1161/JAHA.121.020860 (2021).

13 Voors, A. A. et al. Effects of danicamtiv, a novel cardiac myosin activator, in heart failure with reduced ejection fraction: experimental data and clinical results from a phase 2a trial. Eur J Heart Fail 22, 1649–1658, doi:10.1002/ejhf.1933 (2020).

14 Auguin, D. et al. Omecamtiv mecarbil and Mavacamten target the same myosin pocket despite opposite effects in heart contraction. Nat Commun 15, 4885, doi:10.1038/s41467-024-47587-9 (2024).

15 Planelles-Herrero, V. J., Hartman, J. J., Robert-Paganin, J., Malik, F. I. & Houdusse, A. Mechanistic and structural basis for activation of cardiac myosin force production by omecamtiv mecarbil. Nat Commun 8, 190, doi:10.1038/s41467-017-00176-5 (2017).

16 McMillan, S. N., Pitts, J. R. T., Barua, B., Winkelmann, D. A. & Scarff, C. A. Mavacamten inhibits myosin activity by stabilizing the myosin interacting-heads motif and stalling motor force generation. Sci Adv 12, eaea9335, doi:10.1126/sciadv.aea9335 (2026).

17 Kawas, R. F. et al. A small-molecule modulator of cardiac myosin acts on multiple stages of the myosin chemomechanical cycle. J Biol Chem 292, 16571–16577, doi:10.1074/jbc.M117.776815 (2017).

18 Liu, Y., White, H. D., Belknap, B., Winkelmann, D. A. & Forgacs, E. Omecamtiv Mecarbil modulates the kinetic and motile properties of porcine beta-cardiac myosin. Biochemistry 54, 1963–1975, doi:10.1021/bi5015166 (2015).

19 Rohde, J. A., Thomas, D. D. & Muretta, J. M. Heart failure drug changes the mechanoenzymology of the cardiac myosin powerstroke. Proc Natl Acad Sci U S A 114, E1796–E1804, doi:10.1073/pnas.1611698114 (2017).

20 Woody, M. S. et al. Positive cardiac inotrope omecamtiv mecarbil activates muscle despite suppressing the myosin working stroke. Nat Commun 9, 3838, doi:10.1038/s41467-018-06193-2 (2018).

21 Swenson, A. M. et al. Omecamtiv Mecarbil Enhances the Duty Ratio of Human beta-Cardiac Myosin Resulting in Increased Calcium Sensitivity and Slowed Force Development in Cardiac Muscle. J Biol Chem 292, 3768–3778, doi:10.1074/jbc.M116.748780 (2017).

22 Amrute-Nayak, M. et al. ATP turnover by individual myosin molecules hints at two conformers of the myosin active site. Proc Natl Acad Sci U S A 111, 2536–2541, doi:10.1073/pnas.1316390111 (2014).

23 Usaj, M., Moretto, L., Vemula, V., Salhotra, A. & Mansson, A. Single molecule turnover of fluorescent ATP by myosin and actomyosin unveil elusive enzymatic mechanisms. Commun Biol 4, 64, doi:10.1038/s42003-020-01574-0 (2021).

24 Berg, A., Velayuthan, L. P., Tagerud, S., Usaj, M. & Månsson, A. Probing actin-activated ATP turnover kinetics of human cardiac myosin II by single molecule fluorescence. Cytoskeleton (Hoboken), doi:10.1002/cm.21858 (2024).

25 Mornet, D., Bertrand, R., Pantel, P., Audemard, E. & Kassab, R. Structure of the actin-myosin interface. Nature 292, 301–306, doi:10.1038/292301a0 (1981).

26 Stein, L. A., Greene, L. E., Chock, P. B. & Eisenberg, E. Rate-limiting step in the actomyosin adenosinetriphosphatase cycle: studies with myosin subfragment 1 cross-linked to actin. Biochemistry 24, 1357–1363 (1985).

27 Iwamoto, H., Oiwa, K., Suzuki, T. & Fujisawa, T. X-ray diffraction evidence for the lack of stereospecific protein interactions in highly activated actomyosin complex. J Mol Biol 305, 863–874, doi:10.1006/jmbi.2000.4334 (2001).

28 Bodt, S. M. L. et al. Dilated cardiomyopathy mutation in beta-cardiac myosin enhances actin activation of the power stroke and phosphate release. PNAS Nexus 3, pgae279, doi:10.1093/pnasnexus/pgae279 (2024).

29 Wang, T. et al. Single-molecule analysis sheds light on cardiac myosin dysfunction due to hypertrophic cardiomyopathy mutation A57D in ventricular myosin light chain-1 (MLC1v). bioRxiv, 2026.2003.2006.710098, doi:10.64898/2026.03.06.710098 (2026).

30 Liu, C., Kawana, M., Song, D., Ruppel, K. M. & Spudich, J. A. Controlling load-dependent kinetics of beta-cardiac myosin at the single-molecule level. Nat Struct Mol Biol 25, 505–514, doi:10.1038/s41594-018-0069-x (2018).

31 Rohde, J. A., Roopnarine, O., Thomas, D. D. & Muretta, J. M. Mavacamten stabilizes an autoinhibited state of two-headed cardiac myosin. Proc Natl Acad Sci U S A 115, E7486–E7494, doi:10.1073/pnas.1720342115 (2018).

32 Anderson, R. L. et al. Deciphering the super relaxed state of human beta-cardiac myosin and the mode of action of mavacamten from myosin molecules to muscle fibers. Proc. Natl. Acad. Sci. U. S. A. 115, E8143–E8152, doi:10.1073/pnas.1809540115 (2018).

33 Tamborrini, D. et al. Structure of the native myosin filament in the relaxed cardiac sarcomere. Nature 623, 863–871, doi:10.1038/s41586-023-06690-5 (2023).

34 Somavarapu, A. K., Ge, J., Yengo, C. M., Craig, R. & Padron, R. Cryo-EM reveals how cardiomyopathy therapeutic drugs modulate the myosin motors of the heart. Sci Adv 12, eaed6472, doi:10.1126/sciadv.aed6472 (2026).

35 Wang, T., Nayak, A., Kraft, T. & Amrute-Nayak, M. Single-Molecule Investigation of Load-Dependent Actomyosin Dissociation Kinetics for Cardiac and Slow Skeletal Myosin. Small 20, e2406865, doi:10.1002/smll.202406865 (2024).

36 Wang, T., Brenner, B., Nayak, A. & Amrute-Nayak, M. Acto-Myosin Cross-Bridge Stiffness Depends on the Nucleotide State of Myosin II. Nano Lett 20, 7506–7512, doi:10.1021/acs.nanolett.0c02960 (2020).

37 Woody, M. S., Winkelmann, D. A., Capitanio, M., Ostap, E. M. & Goldman, Y. E. Single molecule mechanics resolves the earliest events in force generation by cardiac myosin. Elife 8, doi:10.7554/eLife.49266 (2019).

38 Mansson, A. & Karatzaferi, C. Editorial: Release of inorganic phosphate from the myosin active site in actomyosin energy transduction. Front Physiol 17, 1823925, doi:10.3389/fphys.2026.1823925 (2026).

39 Debold, E. P., Marang, C. P. & Scott, B. D. The order of things: phosphate release or the power stroke, which does actomyosin do first? Front Physiol 16, 1692606, doi:10.3389/fphys.2025.1692606 (2025).

40 Caremani, M. et al. Multiple pathways of the actin-myosin cycle in energy transduction and the release of orthophosphate in muscle. Front Physiol 16, 1664568, doi:10.3389/fphys.2025.1664568 (2025).

41 Robert-Paganin, J., Pylypenko, O., Kikuti, C., Sweeney, H. L. & Houdusse, A. Force Generation by Myosin Motors: A Structural Perspective. Chem Rev 120, 5–35, doi:10.1021/acs.chemrev.9b00264 (2020).

42 Llinas, P. et al. How actin initiates the motor activity of Myosin. Dev Cell 33, 401–412, doi:10.1016/j.devcel.2015.03.025 (2015).

43 Rahman, M. A., Usaj, M., Rassier, D. E. & Mansson, A. Blebbistatin Effects Expose Hidden Secrets in the Force-Generating Cycle of Actin and Myosin. Biophys J 115, 386–397, doi:10.1016/j.bpj.2018.05.037 (2018).

44 Leff, P. The two-state model of receptor activation. Trends Pharmacol Sci 16, 89–97, doi:10.1016/s0165-6147(00)88989-0 (1995).

45 Berg, K. A. & Clarke, W. P. Making Sense of Pharmacology: Inverse Agonism and Functional Selectivity. Int J Neuropsychopharmacol 21, 962–977, doi:10.1093/ijnp/pyy071 (2018).

46 Pilagov, M. et al. Direct measurement of mavacamten and deoxyATP perturbation of the SRX/DRX ratio in porcine cardiac myofibrils using a simple, accessible and multiplexed approach. J Muscle Res Cell Motil 46, 407–414, doi:10.1007/s10974-025-09712-z (2025).

47 Ma, W. et al. Myosin in autoinhibited off state(s), stabilized by mavacamten, can be recruited in response to inotropic interventions. Proc Natl Acad Sci U S A 121, e2314914121, doi:10.1073/pnas.2314914121 (2024).

48 Brunello, E. & Fusi, L. Regulating Striated Muscle Contraction: Through Thick and Thin. Annu Rev Physiol 86, 255–275, doi:10.1146/annurev-physiol-042222-022728 (2024).

49 Irving, M. Functional control of myosin motors in the cardiac cycle. Nat Rev Cardiol 22, 9–19, doi:10.1038/s41569-024-01063-5 (2025).

50 Nag, S., Gollapudi, S. K., Del Rio, C. L., Spudich, J. A. & McDowell, R. Mavacamten, a precision medicine for hypertrophic cardiomyopathy: From a motor protein to patients. Sci Adv 9, eabo7622, doi:10.1126/sciadv.abo7622 (2023).

51 Mohran, S. et al. The biochemically defined super relaxed state of myosin-A paradox. J Biol Chem 300, 105565, doi:10.1016/j.jbc.2023.105565 (2024).

52 Chu, S., Muretta, J. M. & Thomas, D. D. Direct detection of the myosin super-relaxed state and interacting-heads motif in solution. J Biol Chem 297, 101157, doi:10.1016/j.jbc.2021.101157 (2021).

53 Ribeiro, F. et al. Myosin Post-Translational Modifications Associated With Critical Illness Myopathy. Acta Physiol (Oxf) 242, e70240, doi:10.1111/apha.70240 (2026).

54 Velayuthan, L. P., Moretto, L., Tagerud, S., Usaj, M. & Mansson, A. Virus-free transfection, transient expression, and purification of human cardiac myosin in mammalian muscle cells for biochemical and biophysical assays. Sci Rep 13, 4101, doi:10.1038/s41598-023-30576-1 (2023).

55 Spahiu, E., Uta, P., Kraft, T., Nayak, A. & Amrute-Nayak, M. Influence of native thin filament type on the regulation of atrial and ventricular myosin motor activity. J Biol Chem 300, 107854, doi:10.1016/j.jbc.2024.107854 (2024).

56 Pardee, J. D. & Spudich, J. A. Purification of muscle actin. Methods Cell Biol. 24, 271–289 (1982).

57 Meijering, E., Dzyubachyk, O. & Smal, I. Methods for cell and particle tracking. Methods Enzymol 504, 183–200, doi:10.1016/B978-0-12-391857-4.00009-4 (2012).

58 Balaz, M., Sundberg, M., Persson, M., Kvassman, J. & Månsson, A. Effects of Surface Adsorption on Catalytic Activity of Heavy Meromyosin Studied using Fluorescent ATP Analogue. Biochemistry (Mosc). 46, 7233–7251 (2007).

59 Berg, A., Velayuthan, L. P., Tagerud, S., Usaj, M. & Mansson, A. Probing actin-activated ATP turnover kinetics of human cardiac myosin II by single molecule fluorescence. Cytoskeleton (Hoboken) 81, 883–901, doi:10.1002/cm.21858 (2024).

60 Woody, M. S., Lewis, J. H., Greenberg, M. J., Goldman, Y. E. & Ostap, E. M. MEMLET: An Easy-to-Use Tool for Data Fitting and Model Comparison Using Maximum-Likelihood Estimation. Biophys J 111, 273–282, doi:10.1016/j.bpj.2016.06.019 (2016).

61 Steffen, W. & Sleep, J. Using optical tweezers to relate the chemical and mechanical cross-bridge cycles. Philos Trans R Soc Lond B Biol Sci 359, 1857–1865 (2004).

62 Steffen, W., Smith, D., Simmons, R. & Sleep, J. Mapping the actin filament with myosin. Proc. Natl. Acad. Sci. U. S. A. 98, 14949–14954 (2001).

63 Lewalle, A., Steffen, W., Stevenson, O., Ouyang, Z. & Sleep, J. Single-molecule measurement of the stiffness of the rigor myosin head. Biophys J 94, 2160–2169, doi:10.1529/biophysj.107.119396 (2008).

64 Molloy, J. E., Burns, J. E., Kendrick-Jones, J., Tregear, R. T. & White, D. C. Movement and force produced by a single myosin head. Nature 378, 209–212, doi:10.1038/378209a0 (1995).

65 Amrute-Nayak, M. et al. Transformation of the Nonprocessive Fast Skeletal Myosin II into a Processive Motor. Small 15, e1804313, doi:10.1002/smll.201804313 (2019).

66 Blackwell, T., Stump, W. T., Clippinger, S. R. & Greenberg, M. J. Computational Tool for Ensemble Averaging of Single-Molecule Data. Biophys J 120, 10–20, doi:10.1016/j.bpj.2020.10.047 (2021).

67 McDonald, K. S., Kalogeris, T. J., Veteto, A. B., Davis, D. J. & Hanft, L. M. Myosin binding protein-C modulates loaded sarcomere shortening in rodent permeabilized cardiac myocytes. J Gen Physiol 157, doi:10.1085/jgp.202413678 (2025).

