## Supplemental information and figures for "Myotropes unveil two myosin cycles with distinct kinetics and stroke size"

### Model

*Overall description and justification of model*

The individual model states and transitions follow from previous work[1-4] with control parameter values (Tables S3-S4; Fig. S10) primarily obtained in [4] (based on experimental results; see references in [4]). Our single molecule actin-activated ATPase data (Fig. 2) suggest two different cycles in parallel at saturating OM and Mava concentrations (Fig. 6a) with the drug acting as partial inhibitor that only partly shifts the cycling from a fast ("weakly drug-binding)) to a slow (strongly drug-binding) path. The rate constants associated with drug-binding to myosin were assigned values based on key constraints with minor differences between Mava and OM as motivated below (Figs. S7-S8). Rapid decrease in force in contracting myofibrils (Mava-transients) upon Mava addition[5] suggests strong drug-binding also to actin-attached pre-power-stroke states (Fig. 6a). Notably, such binding is likely to occur primarily to cross-bridges with a positively strained elastic element (cf. effects of varied [Pi])[6,7]. This is because negatively strained actomyosin cross-bridges, 1. would have difficulty of undergoing over-priming of the lever arm required for formation of the pocket for strong drug-binding (Equation S10), and 2. very short lifetimes of pre-power-stroke states due to fast downstream events. In accordance with reversible strong drug-binding to actin-attached states we also expect the reverse transition to weak-binding. The latter may occur at faster rate with Mava for which tumbling of the drug in its pocket has been demonstrated[8].

With the scheme in Fig. 6a as starting point we asked which individual transitions rate-limit the strongly drug-bound paths. For OM, our results from human cardiac myosin and available data [8,9] suggest a nearly completely inhibited power stroke but increased Pi-release rate[10-12]. We implement the increased Pi-release rate by increased attachment rate constant. While we also assume (cf. [4]) increased free energy difference for the $AMDP_{PP}$ -> $AMDP_{PiR}$ transition the functional effects (on turnover rate, gliding velocity, force etc) of the latter change is minimal. With complete power stroke inhibition, an escape pathway must be introduced to account for the observed (Fig. 3, Fig. 5) ATP-independent detachment ($k_{det\text{-}escape} \approx 10\ s^{-1}$) (see also [9,13]). This slow detachment together with the inhibited power stroke is sufficient to explain the OM-induced reduction of the in vitro gliding velocity as experimentally observed and shown in Extended Data Fig. 1. We propose that the slow detachment is followed by a rate limiting transition with rate constant $k_{r\text{-}limit\_slow}$=1-2 $s^{-1}$ presumably associated with completion of the lever arm swing in detached cross-bridges (Fig. 6a; cf. [12]). As mentioned above, we suggest that myosin heads may also escape into the drug-free cycle by drug-release from the pre-power-stroke $AMD_{LDr}$ state under some conditions. Such an effect with OM, with the heads spending prolonged times in the $AMD_{LDr}$ state, may explain why a high-amplitude ATP-dependent detachment process is seen in some experimental studies at saturating drug concentration (e.g. our Fig. 3 and [13]). Otherwise, the kinetic effects of OM predict > 90 % of the flux through the slow strongly drug-bound cycle following a $k_{onDr}[MDP_{Dr2}]/k_{on}([MDP]+[MDP_{Dr}])$ ratio of ~18 based on the rate constants in Tables S3-S4. The dominance of the strongly drug-bound cycle is consistent with our optical trapping data (Fig. 5; see also [9]) but, yet again not with our binding data in Fig. 3 and other optical trapping data[13].

Mava, in contrast to OM, slows Pi-release and actin-activated ATP turnover to similar degree[14,15]. Our trapping data show that the slowest rate constant for transitions between strongly actin-bound pre-power-stroke states is ~40 $s^{-1}$. Therefore, the observed rate limitation (1-2 $s^{-1}$) must reflect reduced rate of actomyosin attachment, consistent with Mava-induced inhibition of the weak-to-strong actomyosin transition[14]. This idea accords with substantial myosin cycling (~ 60 %) through a fast path with weakly bound Mava at saturating [Mava] with equilibrium established between all MT and MDP states as suggested by our ATP turnover

(Fig. 2) and optical trapping (Fig. 4) data. We implement this Mava effect by reduced attachment rate constant ($k_{on}'$) and increased value of its reversal, to also account for data from isometrically contracting myofibrils[5], e.g. with little Mava-induced change in the rate of force development but a very large reduction in steady-state isometric force. Simulations in Extended Data Fig. 1 show that these mechanisms alone could also explain the Mava induced reduction of the maximum in vitro gliding velocity because the gliding in an in vitro motility assay is not fully unloaded. The latter is clear from the presence of stationary filaments and stops and pauses of motile filaments.

The indications from structural analysis[8],[16] that Mava like OM hinders the power stroke is consistent with a prolonged actin bound waiting phase (rate constant ~40 $s^{-1}$) before the power stroke in the two-phase binding events in optical trapping. The mechanism may involve Mava-induced slowing of either of the following transitions: 1. $AMDP_{PP}$ -> $AMDP_{PiR}$, 2. $AMDP_{PP}$ -> $AMD_L$ or 3. $AMDP_L$ -> $AMD_H$ (power stroke). Model predictions of averaged optical trapping data (assuming rate constant 40 $s^{-1}$) for the mechanisms 1 and 3 are similar. The effects are compared in Fig. 6c to simulated conditions without drug (rate constant 1000 $s^{-1}$) or with OM (assumed effective rate constant < 1 $s^{-1}$).

Simulations, incorporating the above ideas into a simple kinetic scheme (Fig. S9), neglecting drug-binding in actin-attached states, predict (Fig. 6c; Fig. S8) double-exponential frequency distributions under saturating drug concentrations for both drugs. The amplitude of a slow phase (1-2 $s^{-1}$) is greatly increased with raised drug concentration in the simulations at the expense of the fast phase (>10 $s^{-1}$). Importantly, like in experiments (Fig. 2), the amplitude of the fast phase exceeds 50 % even at saturating drug concentrations due to a large fraction of the myosin motors cycling through the path with weakly bound drug. The lack of drug-induced change in the lag phase (Extended Data Fig. 2) agrees with previous findings[9-11,13,15,17,18] that neither OM nor Mava affects the ADP-release rate (Fig. S11a). In the simulations, we do not attempt to account for all experimental complexities like one very slow phase (rate constant like basal ATPase) and a multiexponential distribution in the absence of drug. This is justified by dominance of the fast phase (usually 80-90 %) (see also [19]). The quite similar value of two different rate limiting transitions for OM and Mava might seem fortuitous. Importantly, however, the parameter values that account for our single-molecule data quite faithfully reproduce previous conventional steady-state actin-activated ATPase data obtained in solution[12,14,15,17,18] (Fig. S12).

The drug-binding mechanism, involving over-priming of the lever arm for strong drug-binding suggests a mechanism for increased step length of the 2-step events with Mava in the trapping data. Thus, myosin attachment to actin with an over-primed lever arm (locked in place by Mava) is expected to result in a larger swing if the end-point in the post-power-stroke state is the same as without drug. This drug effect was taken into account in the simulations by a right-ward shift of the position of the free energy minima of the $AMDP_{PP}$, $AMDP_{PiR}$ and $AMD_L$ states ($x_1$; Fig. S10, Table S3). The shorter stroke length for the one-step events in optical trapping data could be associated with Mava binding to another site or a weaker binding within the same pocket to shorten the stroke but with minimal effects on the ATP-turnover kinetics. We did not include this effect in the simulations (e.g. by left-ward shift of $x_1$) due to uncertainties about the exact mechanism.

*Rate constants for drug-binding to myosin*

We assume strong binding of the drug only to cross-bridges in pre-power-stroke states with a primed lever arm. This is a slight simplification but generally consistent with structural studies as well as binding studies suggesting about 10-fold higher affinity in pre-power-stroke than post-power-stroke states[8,16,20]. Different properties constrain the drug-binding rate constants: $k_{Dr+}$, $k_{Dr-}$, $k_{Dr2+}$ and $k_{Dr2-}$: 1. $k_{Dr+}$ and $k_{Dr-}$ are assumed to be fast consistent with a diffusion limited process when the drug enters the open drug-binding pocket. 2. The subsequent stereospecific strong binding of the drug would require formation of several bonds with myosin residues whose appropriate location depend on over-priming of the lever arm. Therefore, we assume that this second binding step is slow as well as its reversal that now requires breakage of all bonds. 3. The values of the rate constants used, need to account for the concentration dependence of the drug effects on the basal myosin ATPase and measured drug affinity in pre-power-stroke states. 4. The ratio $k_{Dr2+}/k_{on}$ for actin-dissociated myosin must not be far from 1 to allow a substantial fraction of actomyosin cycles to progress through the drug-free path at saturating drug concentration, 5. The values should account for the observed OM and Mava concentration dependence of the actin-activated ATPase. The values assigned to the rate constants based on these constraints (Fig. 6a, Tables S3-S4) allow good fits of our model to most experimentally observed phenomena. However, there is strong evidence for drug-binding also to actin-attached pre-power-stroke states. The associated rate constants are used in our simulation of gliding velocities and isometric force development but could also play roles in other phenomena as we discuss in the main paper. Essentially, the rate of drug-binding to the actin-attached pre-power-stroke states are governed by the same parameters $k_{Dr+}$, $k_{Dr-}$, $k_{Dr2+}$ and $k_{Dr2-}$.as for actin-detached states but are assumed to be non-zero only at strain-values (details below) that may be associated with over-priming of the lever arm.

*Drug-affinity in detached pre-power-stroke states*

Assuming that either of the drugs can only bind to actin-detached pre-power-stroke states then, based on the scheme in Fig. 6a, it is straightforward to show that the fraction of bound drug would depend on the drug-concentrations according to the following equation:

$$\text{Fraction bound} = \frac{[\text{Drug}]}{\frac{k_{Dr2-}K_D}{k_{Dr2+}}+[\text{Drug}](1+\frac{k_{Dr2-}}{k_{Dr2+}})} \quad (1a)$$

where $K_D = k_{Dr-}/k_{Dr+}$. This relationship can be approximated by a hyperbolic equation if $k_{Dr2-}$ =2.5 $s^{-1}$ and $k_{Dr2+} \geq 15$ $s^{-1}$ as assumed in our modelling:

$$\text{Fraction bound} \approx \frac{[\text{Drug}]}{\frac{k_{Dr2-}K_D}{k_{Dr2+}}+[\text{Drug}]} \quad (1b)$$

If we insert the values for the rate constants $k_{Dr+}$, $k_{Dr-}$, $k_{Dr2+}$ and $k_{Dr2-}$ that we arrive at from our arguments above and our analysis below (Figs. S7-S8) we find a dissociation constant for binding to myosin pre-power-stroke states of 0.3 µM and 0.5 µM for Mava and OM, respectively. The value for OM is in reasonable agreement with that of 0.3 µM measured by isothermal titration calorimetry[20] for the affinity of OM to an analogue of the pre-power-stroke state (myosin with vanadate locked into the active site).

*Strain-dependence of rate functions for transitions between actomyosin states in mechanokinetic model*

The rate constants with argument (x) in Fig. 6a depend on the elastic strain in the myosin head. Here, x is a position coordinate representing a distance between the myosin molecule and its nearest binding site on the actin filament (simplified as 1 per 36 nm). The variable x is defined such that x=0 nm gives zero force in the $AMDP_{PP}$ state under control conditions. The strain dependence of the rate constants (rate functions) are given below as further justified in [4]. Cross-bridge stiffness $k_s(x)$, is assumed to be 2.8 pN/nm except for the AM-state at $x<x_3$ where where the value 0.2 pN/nm[4] is used.

The transition from the detached or weakly bound MDP state to the first stereo-specifically bound pre-power-stroke ($AMDP_{PP}$) state in the drug-free path is governed by:

$k_{on}(x) = k_{on}' \exp(\frac{1}{\gamma} (\Delta G_{on} - (k_s(x)/2)(x-x_1)^2/(k_B T)))$ (S2)

with values for control conditions for $k_{on}'$ and $\Delta$Gon from Tables S3 and S4 and γ set to 2 (cf. [7]) unless otherwise specified.

The reversal of this transition (1) depends on x as:

$k_{on-}(x) = k_{on}' \exp(((1 - \frac{1}{\gamma})(-\Delta G_{on} + k_s(x)/2)(x-x_1)^2/(k_B T)))$ (S3)

The subsequent transition into the Pi-release state, followed by Pi-release, in the drug-free path ($AMDP_{PiR}$) [1,21] and its reversal are governed by:

$k_{Pr+}(x) = k_{Pr+}' \exp(\Delta G_{PiR}/2)$ (S4)

and

$k_{Pr-}(x) = k_{Pr+}' \exp(-\Delta G_{PiR}/2)$ (S5)

The quantity, $\Delta G_{PiR}$ is the difference between the free energy minima of the $AMDP_{PP}$ and the $AMDP_{PiR}$ states. We here lump this step together with the rapid equilibrium for Pi-release and re-binding as follows:

$k_{Pr-}(x) = ([Pi]/(K_C+[Pi]))\, k_{Pr+}' \exp(-\Delta G_{PiR}/2)$ (S6)

This procedure is approximately valid energetically, under the assumption of a constant, low Pi-concentration (<< $K_C$) and very high actual Pi-binding and unbinding rate constants (cf. [22]). It is not strictly valid for modelling effects of large [Pi].

The next transition in the cycle is a structural change associated with the lever-arm swing[23] generally termed the power-stroke; see also [24]. The forward transition rate constant is given by:

$k_{LH+}(x) = k_{LH-} \exp(\Delta G_{LH} + (k_s(x)/2)(x-x_1)^2/(k_B T) - (k_s(x)/2)(x-x_2)^2/(k_B T))$
if $k_{LH+}(x) < 300\,000\ s^{-1}$ else $k_{LH+}(x) = 300\,000\ s^{-1}$ (S7)

whereas the reverse rate constant $k_{LH-}$ (Table S4) is assumed to be independent of x.

Strain dependent rate constants in the presence of drugs ($k_{onDr}(x)$, $k_{on-Dr}(x)$ $k_{Pr-Dr}(x)$, $k_{Pr-Dr}(x)$ and $k_{LH+Dr}(x)$) are defined as for the drug free path (Eqs. S2 – S7 above) but with other appropriate parameter values from Tables S3-S4.

The transition from the $AMD_H$ to the AMD state is governed by:

$$k_5(x) = k_{-5} \exp(\Delta G_{HD} + \frac{k_s(x)(x-x_2)^2}{2k_B T} - \frac{k_s(x)(x-x_3)^2}{2k_B T}) \quad (S8)$$

where $\Delta G_{HD}$ is the free energy difference between the states. The rate constant for the reverse transition is independent of x and equal to $k_{-5}$

We assume that the subsequent ADP-dissociation is effectively irreversible (due to very low ADP concentration) with a rate constant $k_6$ independent of x (Table S4). The rate constant $k_{off}$ (likewise x-independent) for the following detachment reaction from the AM to the MT state is given by (cf. [1]):

$$k_{off} = \frac{k_2[MgATP]}{\frac{1}{K_1}+[MgATP]} \qquad \text{(S9)}$$

where $K_1$ is the equilibrium constant for MgATP binding to the AM state.

Drug binding is either assumed to occur in detached myosin states according to the scheme in Fig. 6a or to pre-power-stroke $AMDP_{PP}$, $AMDP_{PiR}$ or $AMD_L$ states. In the latter case the drug-binding rate constant $k_{Dr2+}*(x)$ is assumed to exhibit a very simple strain dependence:

$k_{Dr2+}*(x) = 0\ s^{-1}$ for $x< x_1$ and

$$k_{Dr2+}*(x) = \frac{k_{Dr2+}[Drug]}{{k_{Dr-}}/{k_{Dr+}}+[Drug]}$$

for $x \geq x_1$ (S10)

The reverse rate constant is taken as:

$k_{Dr2-}*= k_{Dr2-}$ (S11)

The rate constant for escape detachment, $k_{det\text{-}escape}(x)$ was set to zero for all x-values with Mava but was assumed to exhibit a simple strain-dependence as follows with OM:

$k_{det\text{-}escape}(x) = 0\ s^{-1}$ for $x>x_{10}$

but

$k_{det\text{-}escape}(x) = k_{ADP\text{-}escape}\ (x_{10}-x)$ for $x\leq x_{10}$ (S12)

*Force-velocity data from state probabilities obtained by solution of differential equations*

State probabilities for solving steady-state force-velocity relationships, were derived by solving a set of ordinal differential equations in state probabilities indicated by [] for each model state. Note, that the transitions governed by $k_{Pir}$ and $k_{PirDr}$ are lumped together with the subsequent Pi-release as defined in Eqs. S4-S6. This leaves the Pi-release states $AMD_{PiR}$ and $AMD_{PiRDr}$ from Fig. 6a outside the equations below.

$$\frac{d[MT]}{dt}=(-k_{+3}\ [MT]+k_{-3}[MDP]+k_{off}\ [AM]+k_{det\text{-}escape}(x)([AMD_{LDr}])/v$$
(S13)

$$\frac{d[MDP]}{dt}=(k_{+3}[MT]+k_{on-}(x)[AMDP_{PP}]+k_{Dr2-}*[MDP_{Dr}]-(k_{Dr+}[Drug]+k_{on}(x)+k_{-3}) \qquad [MDP])/v$$
(S14)

$\frac{d[AMDP_{PP}]}{dt}$=(k$_{on}$(x)([MDP]+[MDP$_{DR}$])+k$_{Pr-}$(x)[AMD$_L$]+k$_{Dr2-}$*[AMD$_{PPDr}$]- (k$_{on-}$(x)+k$_{Pr}$(x)+k$_{Dr2+}$*(x)[Drug])[AMDP$_{PP}$])/v (S15)

$\frac{d[AMD_L]}{dt}$=(k$_{LH-}$(x)$[AMD_H]$+k$_{Pr+}$(x)[AMDP$_{PP}$]+k$_{Dr2-}$*[AMD$_{LDr}$]-(k$_{LH+}$(x)+ k$_{Pr-}$(x)+ k$_{Dr2+}$*(x)[Drug]) $[AMD_L]$)/v (S16)

$\frac{d[AMD_H]}{dt}$=(k$_{LH+}$(x)$[AMD_L]$+k$_{LH+Dr}$(x)$[AMD_L Dr]$+k-$_5$(x)[AMD]-(k$_{LH-}$(x) +k$_5$(x)))$[AMD_L]$)/v (S17)

$\frac{d[AMD]}{dt}$=(k$_5$(x)$[AMD_L]$-(k$_6$+k-$_5$(x))[AMD])/v (S18)

$\frac{d[AM]}{dt}$=(k$_6$[AMD]-k$_{off}$[AM])/v (S19)

$\frac{d[MDP_{Dr}]}{dt}$=(k$_{Dr+}$[Drug][MDP]+k$_{Dr2-}$[MDP$_{Dr2}$]-(k$_{Dr-}$+k$_{Dr2+}$)[MDP$_{Dr}$])/v (S20)

$\frac{d[MDP_{Dr2}]}{dt}$=-(k$_{Dr2+}$ [MDP$_{Dr}$] + k$_{on-Dr}$[AMDP$_{Dr}$] - (k$_{on-Dr}$ + k$_{Dr2-}$)[MDP$_{Dr2}$])/v (S21)

$\frac{d[AMDP_{PPDr}]}{dt}$=(k$_{on+Dr}$(x)[MDP$_{Dr2}$]+k$_{Pr-Dr}$(x)[AMD$_{LDr}$]+ k$_{Dr2+}$*(x)[Drug][AMDP$_{PP}$]-(k$_{on-Dr}$(x)+k$_{Pr+Dr}$(x)+k$_{Dr2-}$)[AMDP$_{PPDr}$])/v (S22)

$\frac{d[AMD_{LDr}]}{dt}$=(k$_{LH-}$(x)$[AMD_H]$+k$_{Pr+-DR}$(x)[AMDP$_{PPDr}$]+k$_{Dr2+}$*(x)[Drug]$[AMD_L]$-(k$_{LH+Dr}$(x)+k$_{Pr-}$(x)+ k$_{Dr2-}$*+k$_{det-escape}$) $[AMD_{LDr}]$)/v (S23)

The initial values in the numerical simulations were set to zero for all actin-attached states when the myosin heads are instead assumed to equilibrate between the detached states MT, MDP, MDP$_{Dr}$ and MDP$_{Dr2}$. The initial values at different concentrations of OM were calculated from the steady-state probabilities under these assumptions using the rate constants in Tables S3-S4. These calculated values, exemplified for a standard set of parameter values consistent with the observed effects of OM on the hydrolysis equilibrium are given in Table S1. The values were obtained by solving the linear equation system for the steady-state distribution using Wolfram Mathematica.

Observable variables were calculated from state probabilities [25] by averaging over the distance 36 nm assumed to exist between myosin binding sites along the actin filament. Average force <*F*> (in pN) divided by the total number of myosin heads is given by:

$$< F > \quad \sum_1^{natt} \int_{-91}^{14} k_s \; a_j(x)(x - x_j) dx \Big/ \sum_1^{ntot} \int_{-22}^{14} a_j(x) dx \qquad \text{(S24)}$$

Where “natt” is the total number of attached cross-bridge states and ntot is the total number of states (whether the cross-bridges are attached or detached). Thus, the denominator sums over all states and x-values.

For computational stability, the values of the rate function from Eqs. S2-S10 were limited to the range between $r_{min} = 1\ 10^{-6}\ s^{-1}$ and $r_{max} = 300\ 000\ s^{-1}$. The parameter value was set to $r_{max}$ or $r_{min}$ if either limit was exceeded.

As previously described[4] the simulated maximum isometric force under control conditions was approximated as the force at a velocity of 0.5 nm/s (< 1/1000 of the unloaded shortening velocity).

We solved the differential equations numerically using a Runge-Kutta Fehlberg (4/5) algorithm implemented in the program Simnon[26] (see also [3,27-29]. The code is reproduced below with commenting that would make it readily translatable to other programming languages.

*Simulations of power-stroke*

We simulated displacement traces reflecting averages of many binding and power-stroke events by solution of ordinary differential equations in state probabilities as described previously[3]. This procedure is consistent with averaging approaches that can be applied to single-molecule data[9]. In the simulations, myosin heads were always assumed to be clamped to zero force and attachment of heads to actin and detachment could be neglected because of large separation of relevant time domains. Initially, as in single molecule optical tweezers experiments, the myosin heads were assumed to be in an early pre-power-stroke state simulated by setting $[AMDP_{pp}] = 1$ (fractional population) whereas other state probabilities were set to zero. We simulated the power stroke by shifting the cross-bridges along the x-coordinate from $x=x_1$ to $x=x_2$ under the assumption that tension is clamped to zero. We assumed compliant optical traps[9],[30] in the simulations implemented by setting the cross-bridge stiffness to 0.07 pN/nm (similar to that inferred from the stiffness of the trap acting in series with the cross-bridge) instead of 2.8 pN/nm.

The displacement time ($t$) course (power stroke progression) of the myosin head strain $\Delta L(t)$, initially in the $AMDP_{PP}$ state at $x=x_1$ (i.e. with force clamped to zero) is given by:

$$\Delta L(t)=([AMD_L](t)*(x_1-x_2)+([AMD]+[AMD])(t)*(x_1-x_3))/\ ([MDP](t)+[AMDP_T](t)+[AMD_L](t) + [AM] + [AMD]) \qquad (S25)$$

where (t) indicates a functional dependence on $t$ whereas [ ] indicate state probability.

The power-stroke was simulated essentially as in ref. [3] for either the path without (left path in Fig. 6a) or with drug (right path in Fig. 6a).

*Kinetic scheme simulating effects of varied drug concentration on maximum actin activated ATPase*

In contrast to mechanical events such as power-stroke, force development and gliding velocity, we use a simple kinetic scheme (Fig. S9) that does not consider strain-dependence of parameter values, to simulate steady-state actin-activated ATPase in solution. We also simulate cumulative frequency distributions for single turnover actin-activated ATPase in the single molecule assay (Fig. 2) using this kinetic scheme.

The use of a simple kinetic scheme is possible because no elastic forces act between myosin and actin in the mentioned experiments and all rate constants correspond to those at the minimum of the free energies of the model states. For this purpose, the rate constants are denoted by the values indicated in Fig. 6a but without the (x)-arguments. The scheme is further simplified for our purposes by assuming that the attachment of myosin to actin is irreversible in the case of investigating OM effects. This is justified by the fast subsequent transition from the $AMDP_{PP}$ state into the $AMDP_{PiR}$ state.

The maximum actin-activated ATPase ($k_{cat}$) as well as flux through different paths was obtained by first calculating the steady state probability of the different states in the scheme in Fig. S9. This was done by analytically solving the corresponding system of linear equations using Wolfram Mathematica. Subsequently the actin-activated ATPase was calculated as $k_{cat} = [AM] \times k_2 [ATP]/(1/K_1+[ATP]$, taking [ATP]≡[MgATP] as 5 mM.

To simulate the frequency distributions arising from many single molecule actin-activated ATP turnovers (experiments in Fig. 2), we solved the system of differential equations governing the scheme in Fig. S9 using Simnon (commented version of code given below).

**Supplementary Discussion**

*Drug's effects on different myosin preparation*

Many previous studies of Mava and OM effects on β-myosin have utilized either porcine, bovine or human β-myosin heavy chains co-expressed in C2C12 cells with mouse skeletal muscle light chains. Importantly, our study mitigates concerns about the translational relevance of our studies by utilizing human native cardiac myosins. It avoids the discrepancies arising from species- and isoform-specific differences in the motor components. Notably, our key findings are consistent across both expressed myosin and full-length native myosin from donor hearts (with human ventricular light chains), suggesting two different cycles of ATP-turnover at saturating drug concentrations. Both expressed and native myosins reveal similar drug-induced changes in the relative contribution of the two ATP turnover processes. Our in vitro motility data for full length myosin across varying drug concentrations (our Fig. 1) show qualitative agreement with previous studies using expressed myosin[18],[16] (e.g. Extended Data Fig. 1). Moreover, our model provides a reasonable account of the drugs effects on isometric force and number of attached cross-bridges in muscle sarcomeres[5,31-33].

*On the validity of our single molecule actin-activated ATPase assay*

As we considered in some detail previously[19] there is convincing evidence from several types of studies[34-43] that the EDC-based actomyosin cross-linking via a flexible myosin surface loop[44,45] does not interfere with the myosin dynamics critical for a normal actin-activated ATP turnover. Specifically, $k_{cat}$ of the actin-activated ATP turnover rate for human cardiac $S1^E$ in the absence of drug is similar whether it is obtained using NADH-based steady-state solution studies ([46] and references therein) or the fastest rate constant in triple exponential fits as in the main Fig. 2f-h. A somewhat lower value in previous solution based studies (about 8 $s^{-1}$) than in Fig. 2 (about 12 $s^{-1}$) is expected from a fraction of inactive heads[46] in the solution assays. Such heads do not influence results of the single molecule assay.

The idea that myosin cross-linked via EDC to actin appropriately reflects the actin-activated ATPase without cross-linking is substantiated by comparisons between fast skeletal muscle and human cardiac myosin. Previous solution studies[40] suggested $k_{cat}$ around 50 $s^{-1}$ for fast skeletal muscle myosin at 25 °C appreciably higher than our solution value[46] for human cardiac myosin $S1^E$ of 8 $s^{-1}$ at 23 °C. In accordance with these earlier findings, we found a fastest rate constant of ~50 $s^{-1}$ at 23 °C in a triple exponential fit to cumulative distribution functions derived from EDC-cross-linked skeletal muscle heavy meromyosin in preliminary studies (with camera frame rate 100 $s^{-1}$). The higher value than for human cardiac myosin (~10 $s^{-1}$) accords with the previous solution assays (e.g. [46] vs. [40]).

Further supporting the validity of the assay with EDC cross-linked actomyosin, several effects of OM and Mava that can be compared to previous solution studies are consistent between our single molecule data and results from the earlier work. This includes lack of evidence for changes in the ADP-release rate and the rate constant of the hydrolysis equilibrium suggested by the lag in the frequency distributions in the main Fig. 2 (see also Extended Data Fig. 2 and Fig. S11). Moreover, the effects of OM and Mava on $k_{cat}$ in previous solution-based steady-ATPase assays (Fig. S12) are reasonably consistent with the changes observed in the amplitudes and rate constants in the main Fig. 2.

A further issue that deserves consideration is the amplitudes of the different exponential processes in Fig. 2 that are derived directly from the exponential fits. Under some experimental conditions, correction for different rate constants of the associated process[47] has been claimed to be necessary. Importantly, however, in the present case such corrections are not required. This is because the time associated with each cycle from ATP-binding to ADP release is not rate limited by the rate-limiting step of the ATP turnover. Under our experimental conditions, the latter is fast compared to the waiting time for ATP-binding.

### Supplementary Figures

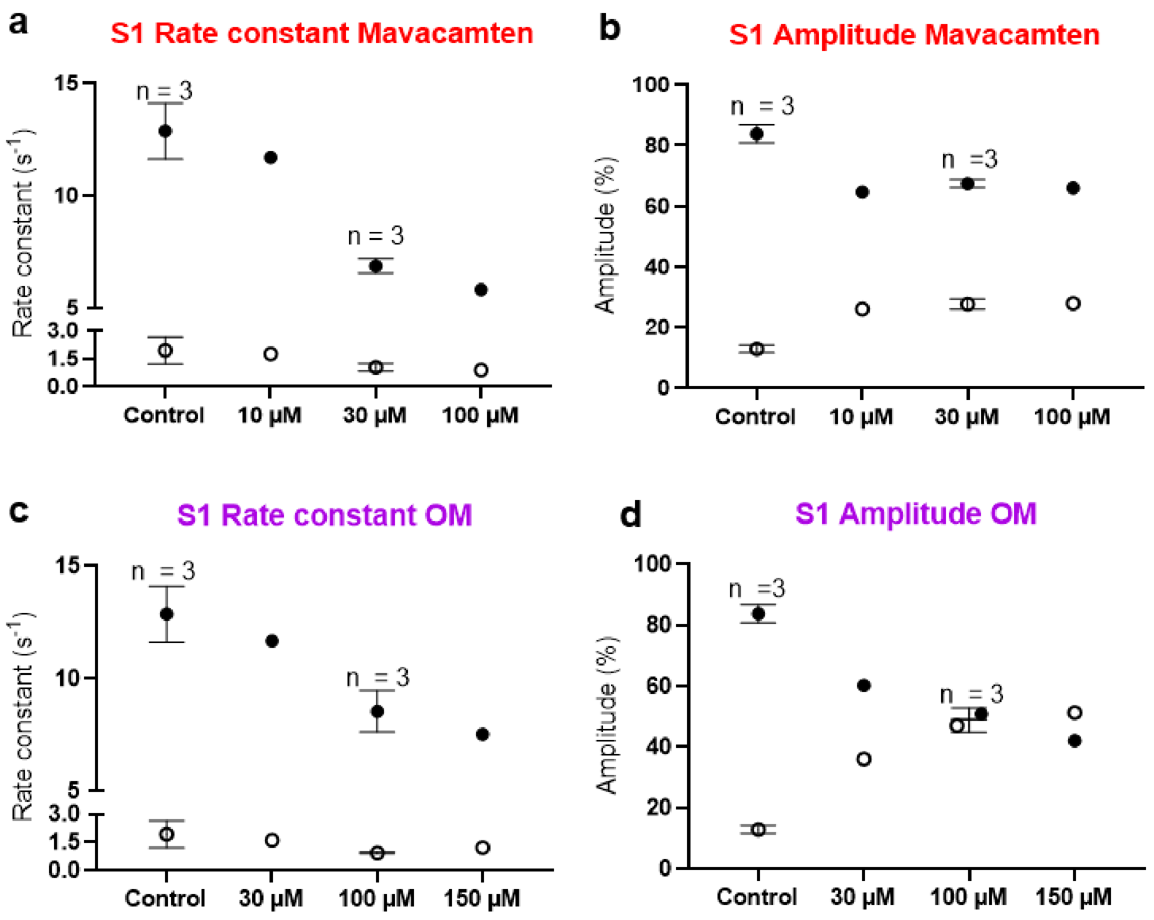


**Figure S1 (Related to Figure 2): Different concentrations of either Mava (10-100 µM) or OM (30-150 µM) and their effects on the rate constants and amplitudes in actin activated ATPase of single headed myosin construct S1$^{E}$. a-b**) Rate constants and amplitudes obtained from fits of cumulative dwell time distributions with different concentrations of Mava. **c-d**) Rate constants and amplitudes obtained from fits of cumulative dwell time distributions with different concentrations of OM. At observed saturation the experiments were repeated at 3 individual occasions using different myosin preparations (same data as in main Fig. 2). Error bars = SEM. $n_{dwells} \approx 1000$ for each repetition. Closed circles: fast process ($k_{fast}$), Open circles: intermediate process ($k_{middle}$). We view the findings reliable that addition of Mava and OM change the rate constants, only at the highest concentrations (30 and 100 µM) tested, considering good reproducibility between experiments (cf. Fig. 2g, h).

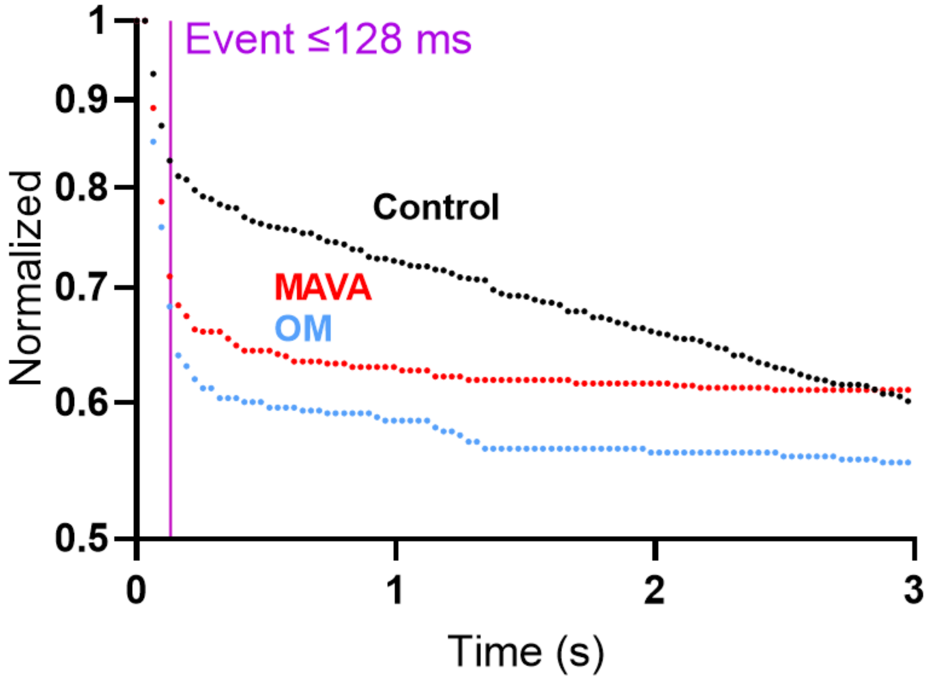


**Figure S2 (Related to Figure 2). Contribution of unspecific Alexa 647-ATP surface interaction to the overall cumulative dwell time distribution of basal ATPase.** Left side of purple line (<0.128 ms) represents the overall population of unspecific events collected under basal conditions. The events are referred to as unspecific because they are observed at similar frequency in regions of interest without myosin molecules. Few specific (myosin-interacting) dwell times were observed with Mava ($N_{dwell}$ = 465) and OM ($N_{dwell}$ = 391) compared to control conditions ($N_{dwell}$=1053) due to long lifetimes relative to the total sampling time.

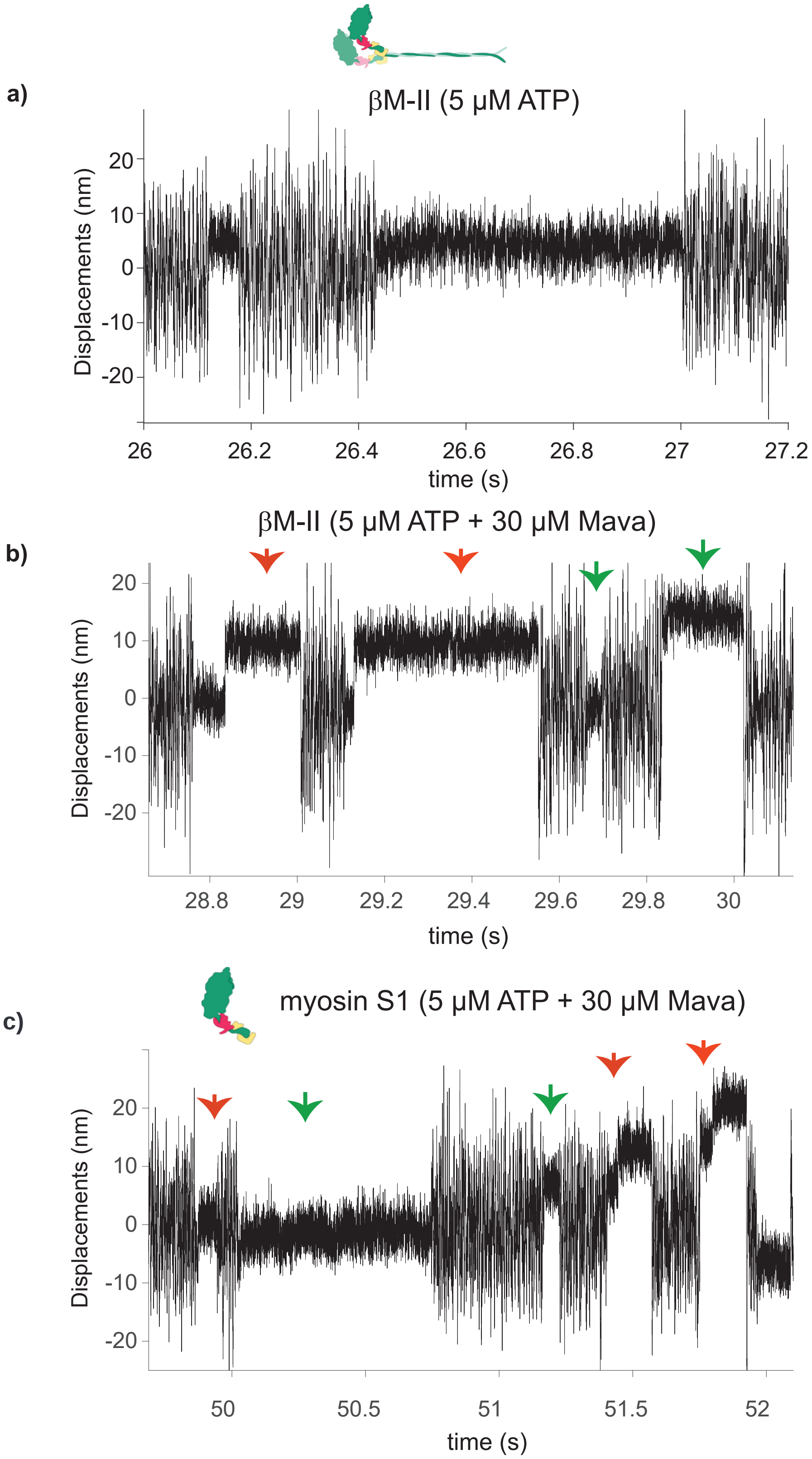


**Figure S3 (Related to Fig. 4)**. Original data traces (displacement over time records) from optical trapping measurements with and without Mava to clearly demonstrate the presence of apparent single and 2-step actomyosin interaction events indicated with green and red arrows, respectively within the indicated duration. Note that only single interaction events are detected in no drug control and in the presence of OM.

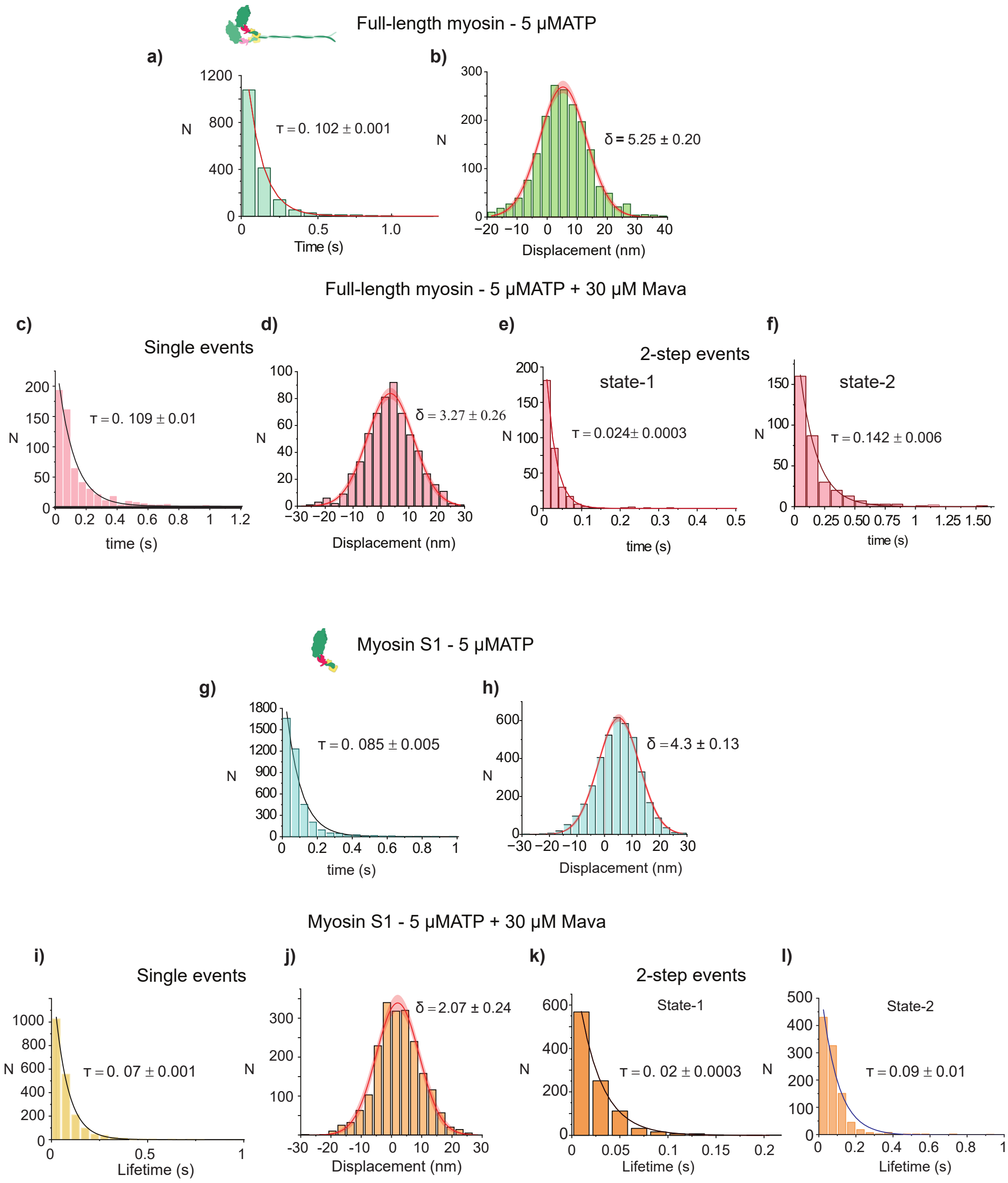


**Figure S4 (related to Fig. 4). Analyses of effects of Mava (30 µM) at 5 µM ATP on native βM-II and Myosin S1 prepared from native βM-II. a)** Full-length cardiac myosin probed for its interaction with actin at 5 µM [ATP]. Histogram showing lifetime of actomyosin binding events fitted with a single exponential decay function yielding an average lifetime of 0.102 ± 0.001 sec. **b)** The displacements from individual actomyosin interaction events plotted in a histogram is fitted with a Gaussian function to derive the average stroke size using 'histogram-shift' method, yielding a stroke size of 5.25 ± 0.2 nm. N = 17, n= 1791. **c - f)** Trapping Measurements at 5 µM ATP in the presence of 30 µM Mava. **c)** Event lifetimes from apparent single step binding events fitted to a single exponential decay function. **d)** Stroke size estimation for single step events. N = 9, n = 592. **e and f)** Event lifetimes from state-1 and state-2 of apparent two-step binding events fitted to get an average time constant. N = 9, n = 333 each for state-1 and state-2. **g-h)** Cardiac myosin S1 probed for its interaction with actin at 5 µM [ATP], to get an average lifetime **(g)** and stroke size **(h)**. N = 12, n = 3205. **i-l)** Myosin-S1 interactions with actin in the presence of 30 µM Mava. Event lifetimes from apparent single step binding events fitted to a single exponential decay function **(i)** and stroke size estimation for single step events **(j)**. **k-l)** Event lifetimes from state-1 and state-2 of apparent two-step binding events fitted to get an average time constant. N = 21, n = 2054. N = 21, n = 1000. N= number of myosin molecules, n = number of actomyosin binding events.

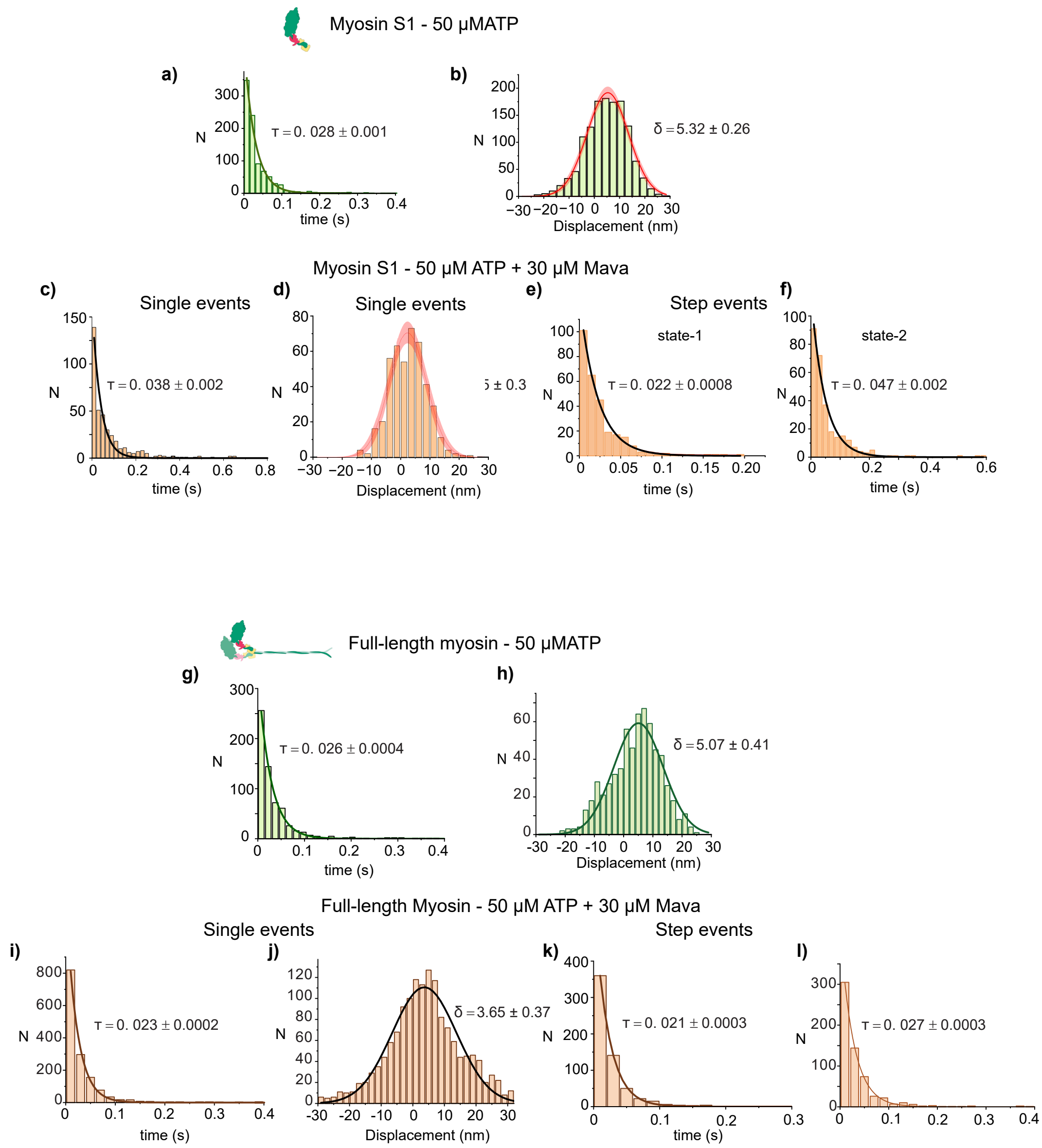


**Figure S5 (Related to Figure 4). Analyses of effects of Mava (30 µM) at 50 µM ATP on native βM-II. a)** Cardiac myosin S1 probed for its interaction with actin at 50 µM [ATP], and average lifetime (a) and stroke size (b) was estimated. N = 7, n = 918. **c-f)** Measurements at 50 µM ATP in the presence of 30 µM Mava. Apparent single events fitted to derive the average interaction duration (c) and stroke size (d) N = 8, n = 388. **e-f)** 2-step events – average lifetime of state-1 and state-2 estimated from data fitted to single exponential decay functions. N = 8, n = 288 for state-1 and state-2. **g-h)** Full-length myosin probed for its interaction with actin at 50 µM [ATP], and an average lifetime (g) and stroke size (h) were estimated. N = 5, n = 633. **i-l)** Myosin interactions with actin in the presence of 50 µM ATP and 30 µM Mava analyzed. Event lifetimes from apparent single step binding events fitted to a single exponential decay function (i) and stroke size estimation for single step events (j). **k-l)** Event lifetimes from state-1 and state-2 of apparent two-step binding events fitted to get an average time constant. N = 21, n = 2054. N = 21, n = 1000 each for state-1 and state-2. N = number of myosin molecules, n = number of actomyosin binding events.

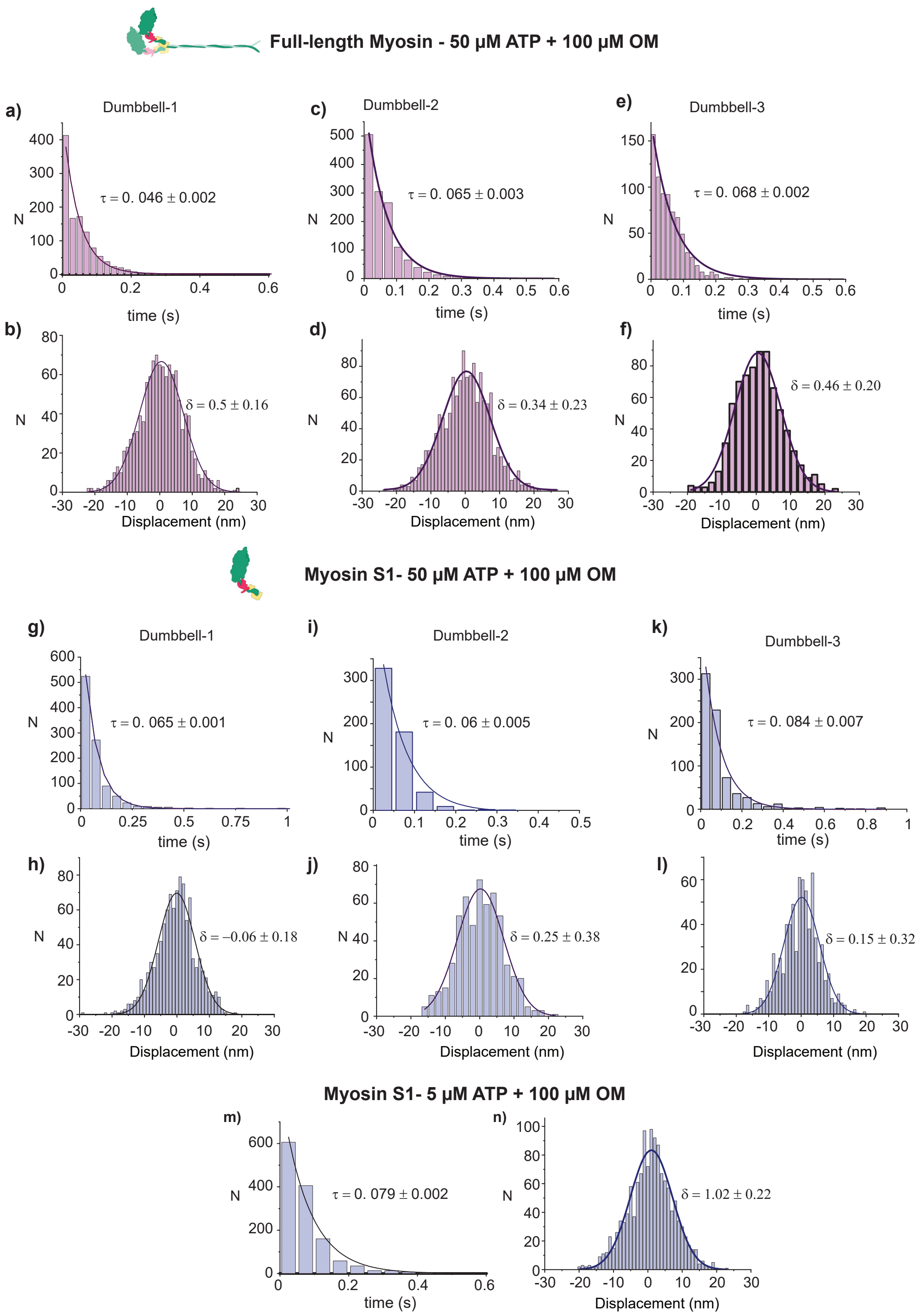


**Figure S6 (Related to Figure 5). Analyses of effects of OM (100 µM) on native βM-II and Myosin S1 prepared from native βM-II. a-f)** Full-length cardiac myosin probed for its interaction with actin for event lifetimes and displacement at 50 µM [ATP] in the presence and absence of 100 µM OM. **(a-b)** Actin dumbell 1- 50 µM ATP + 100 µM OM - N= 4, n= 1138, **(c-d)** Actin dumbell 2- 50 µM ATP + 100 µM OM - N= 5, n= 1355, **(e-f)** Actin dumbell 3 - 50 µM ATP + 100 µM OM - N= 3, n= 744. **g-n)** Myosin S1-actin interactions at 5 and 50 µM [ATP] in the presence and absence of 100 µM OM. **(g-h)** Actin dumbell 1- 50 µM ATP + 100 µM OM - N= 5, n= 1010, **(i-j)** Actin dumbell 2- 50 µM ATP + 100 µM OM - N= 3, n= 565, **(k-l)** Actin dumbell 3- 50 µM ATP + 100 µM OM - N= 4, n= 725, **(m-n)** 5 µM ATP + 100 µM OM - N= 5, n= 1314. N= Number of myosin molecules, n= number of binding events.

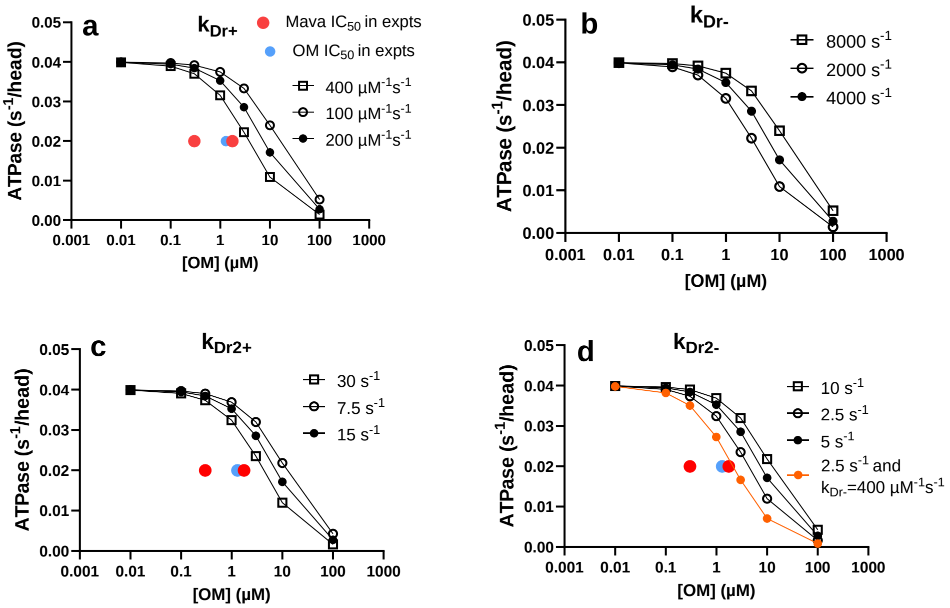


**Figure. S7. Effects of limited changes in the rate constants $k_{Dr+}$, $k_{Dr-}$, $k_{Dr2+}$ $k_{Dr2-}$ on basal myosin ATPase vs drug concentration according to scheme in main Fig. 6a. a.** Changes in $k_{Dr+}$ between 100 and 400 µM$^{-1}$ s$^{-1}$ assuming diffusion limited weak binding of either OM or Mava. OM and Mava concentrations ($IC_{50}$) for 50 % inhibition of the basal ATPase in experiments are indicated. **b.** Changes in $k_{Dr-}$ between 2000 and 8000 s$^{-1}$. **c.** Changes in $k_{Dr2+}$ between 7.5 and 30 s$^{-1}$. **d.** Changes in $k_{Dr2-}$ between 2.5 and 10 s$^{-1}$.

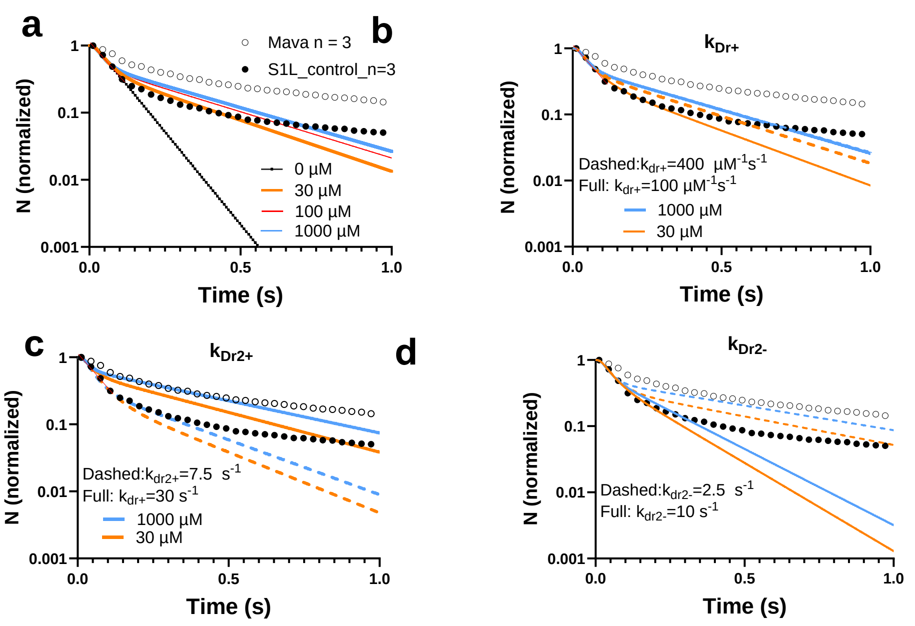


**Figure. S8. Effects of changes in the rate constants $k_{Dr+}$, $k_{Dr-}$, $k_{Dr2+}$ $k_{Dr2-}$ on double exponential frequency distributions reflecting single-molecule ATPase.** **a.** Comparison of simulated frequency distributions (lines) at different Mava concentrations to experimental distributions under control conditions (filled black circles) and with 30 µM Mava (open black circles). The simulated data are obtained using standard parameter values from Tables S3-S4 but with $k_{Dr+}$= 200 µM$^{-1}$s$^{-1}$ $k_{Dr-}$ = 2000 s$^{-1}$, $k_{Dr2+}$ = 15 s$^{-1}$ and $k_{Dr2-}$ = 5 s$^{-1}$. Note, remaining fast exponential component at close to saturating drug concentrations of 30 – 1000 µM. The rather appreciable difference from experimental data is due to the additional slowing of the fast rate constant and the presence of a third slow exponential process not included in the simulations. **b.** Effects of changes in $k_{Dr+}$ on simulated frequency distributions at 30 and 1000 µM Mava. Note that dashed ($k_{Dr+}$=400 µM$^{-1}$s$^{1}$) lines show appreciably smaller difference between orange (30 µM Mava) and blue (1000 µM) lines suggesting increased affinity. The situation is opposite for the full lines ($k_{Dr+}$=100 µM$^{-1}$s$^{1}$). There are only minimal changes in slope of the slow phase and its fractional amplitude at 1000 µM with varied $k_{Dr+}$ suggesting unchanged slow rate constant and unchanged fractional contribution of the slow process. **c** Effects of changes in $k_{Dr2+}$. Note that dashed ($k_{Dr2+}$=7.5 s$^{1}$) lines show similar difference between orange (30 µM Mava) and blue (1000 µM Mava) as full lines ($k_{Dr2+}$=7.5 s$^{1}$), i.e. only small changes in affinity. However, increased value of $k_{Dr2+}$ both reduces the slope of the slow phase and its fractional amplitude. **d.** Changes in $k_{Dr2-}$. In analogy with changes in c, dashed ($k_{Dr2-}$=2.5 s$^{1}$) lines show similar difference between orange (30 µM Mava) and blue (1000 µM) as full lines ($k_{Dr2-}$=10 s$^{1}$), i.e. only small changes in affinity. However, reduced value of $k_{Dr2-}$ reduces the slope of the slow phase with minimal effect on its fractional amplitude.

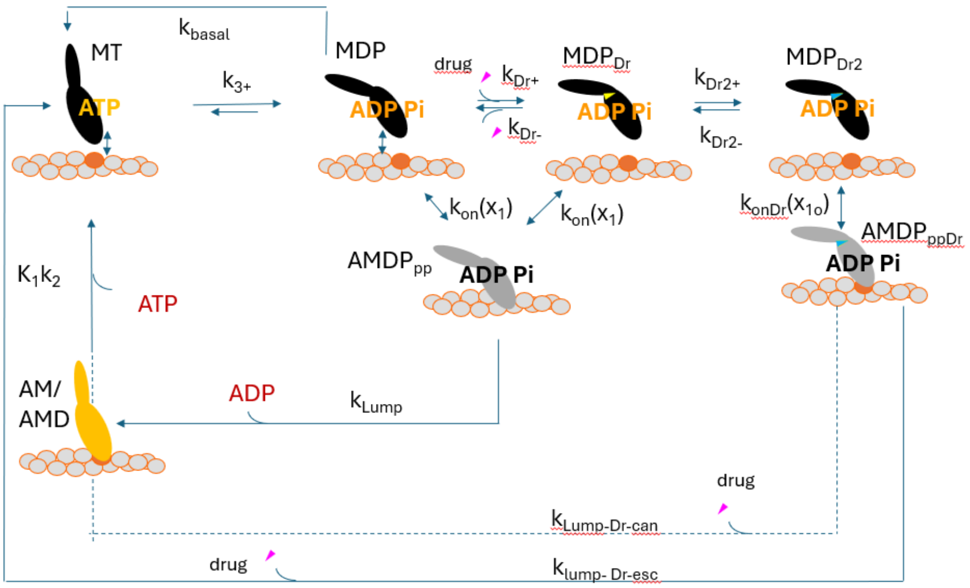


**Figure S9. Simplified version of model for simulation of frequency distributions in single molecule data and steady-state ATPase results vs drug concentration.** In these simulations we made approximations of parameter values as in Fig. 6a and Tables S3-S4: $k_{lump}$= $k_6$, $k_{lump\text{-}Dr\text{-}can}$= 40 $s^{-1}$ for Mava and 0 $s^{-1}$ for OM. Finally, $k_{lump\text{-}Dr\text{-}esc}$ = $k_{r\text{-}lim\text{-}slow}$ (0 $s^{-1}$ for Mava assuming no escape pathway and 1 $s^{-1}$for OM). In the fast (drug-free) cycle and in the slow cycle with OM the reversal of the myosin-actin attachment transition was assumed to be irreversible in the calculations due to very fast down-stream transitions after attachment. In the subscripts, “can” and “esc” refer to “canonical” and “escape”, respectively.

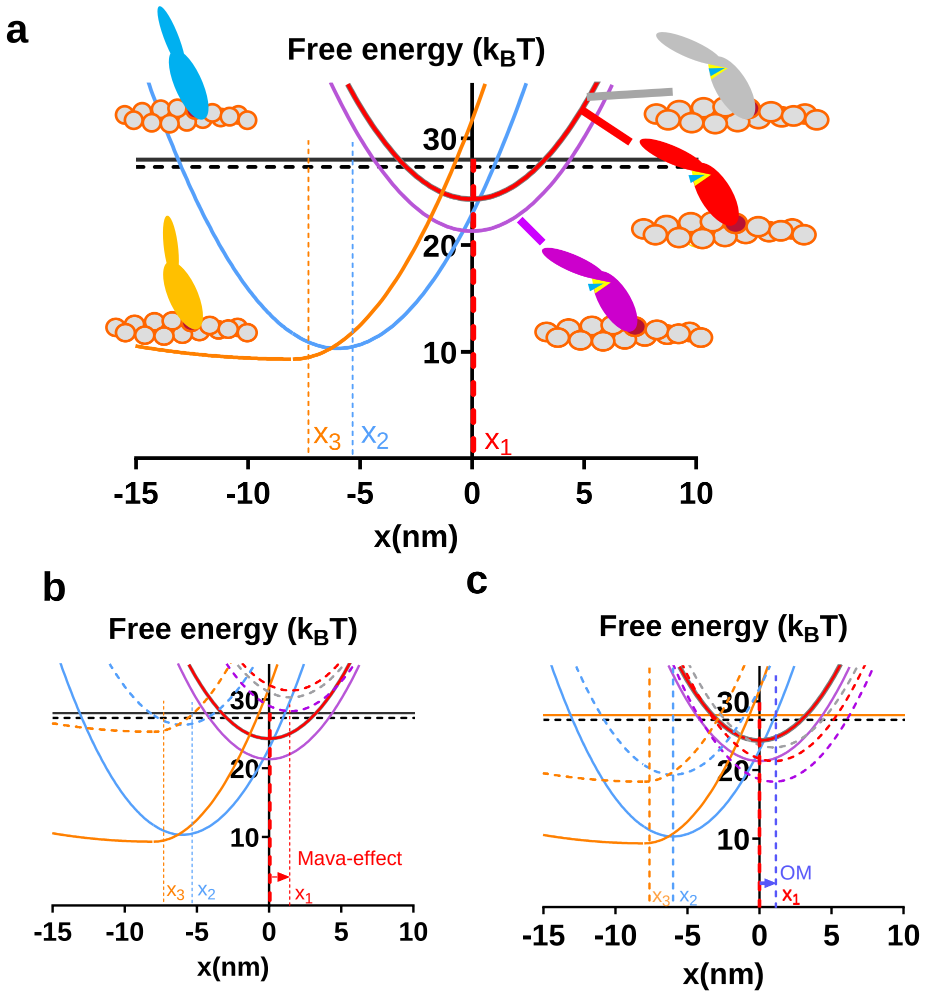


**Fig. S10. Free energy diagrams for modelling of control conditions and conditions with the presence of Mava and OM. a.** Diagrams for control conditions with positions for free energy minima, $x_1$, $x_2$ and $x_3$ indicated. Association of diagrams with schematic illustrations of states in main Fig. 6a also indicated. **b.** Diagrams for simulating effects of Mava. The diagrams for the fast path (full lines and dashed black line) are assumed similar to those for the control conditions without any changes, e.g. in position of free energy minima. The diagrams for the slow (strongly drug-bound) path are indicated by dashed lines (and full black line) and are associated with shift of the pre-power-stroke states to larger x-values. No shift to lower x-values were assumed for the fast phase. **c.** Diagrams for simulating effects of OM. The diagrams for the fast path (full lines) are assumed similar to those for the control conditions without any changes, e.g. in position of free energy minima. The diagrams for the slow (strongly drug-bound) path are indicated by dashed lines and are also for OM assumed to be associated with shift of the pre-power-stroke states to larger x-values.

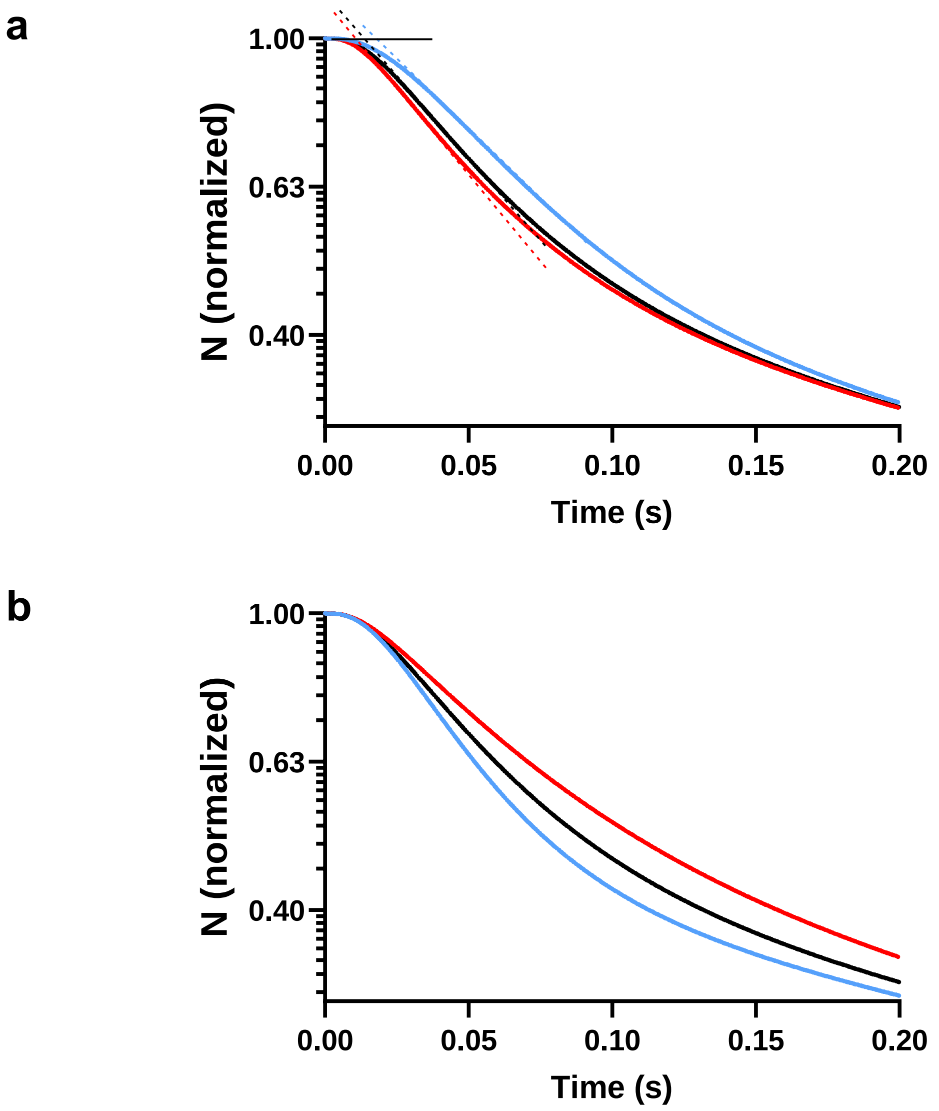


**Figure. S11. Information from early phase of frequency distribution from single molecule actin-activated ATPase**. **a.** Different values of $k_{lump}$ from Fig. S9 (100 $s^{-1}$: black, 150 $s^{-1}$: red and 50 $s^{-1}$: blue), governing the ADP release rate in the fast, presumably drug free path. Note, clearly prolonged lag phase with reduction in $k_{lump}$. **b.** Different values of $k_{3-}$ (50 $s^{-1}$: black, 100 $s^{-1}$: red and 17 $s^{-1}$: blue).

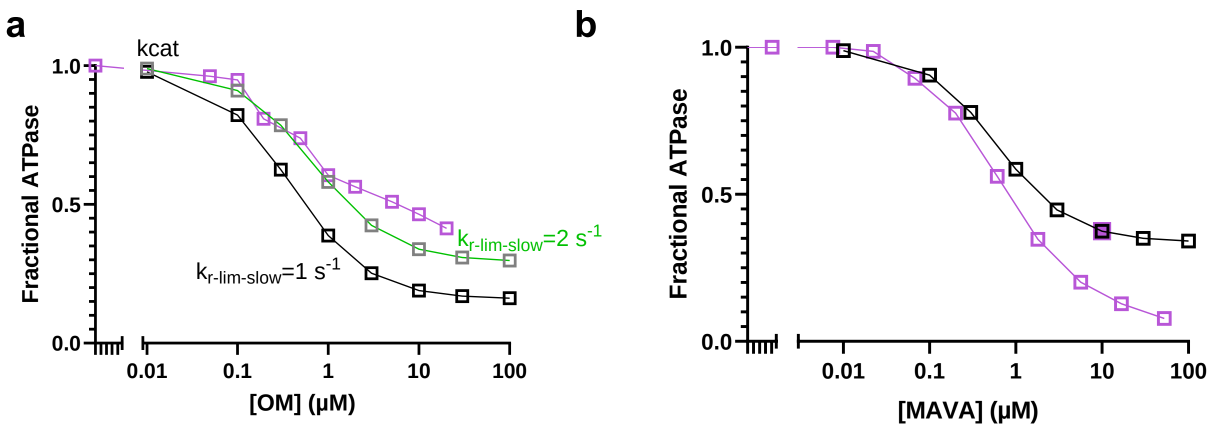


**Fig. S12. Predictions of model in Fig. 6a of steady-state actin-activated myosin ATPase in solution assays in comparison to experimental data. a)** Comparison of experimental data (purple) from Swenson et al and Fig. 3 for steady-state actin-activated ATPase (open squares) to model results for different OM concentrations obtained using two different sets of parameter values (black and green) as indicated. **b)** Comparison of simulated data (black) to experimental data (purple) from Kawas et al.[14](open symbols) for a series of experiments at different Mava concentrations using 14 µM actin. An experimental $k_{cat}$ value at 10 µM Mava (filled purple square) is also shown for an experiment with extrapolation to infinite actin concentration from ATPase data at a range of actin concentrations. The data at 14 µM actin are likely to be underestimated at high Mava-concentrations due to reduced actin affinity with Mava. All data simulated using parameter values in Tables S3-S4.

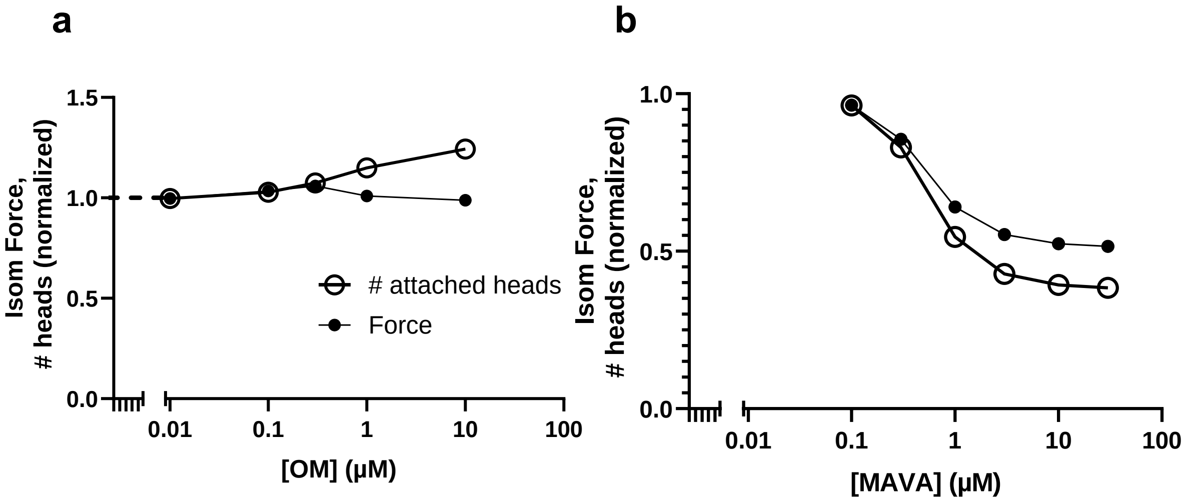


**Figure S13. Predictions of model in Fig. 6a of force and number of attached cross-bridges during isometric contraction. a)** OM effects on isometric force (filled symbols) and the number of attached cross-bridges (open symbols). **b)** Mava effects on isometric force (filled symbols) and the number of attached cross-bridges (open symbols). All data simulated using parameter values in Tables S3-S4. Note, the simulated isometric force for OM was quite sensitive to small changes in model parameter values, giving either small decrease or small increase of force with increased drug concentration. This is reminiscent of similar findings in experiments. For all different model parameters tested with OM we always found an increased number of attached cross-bridges with increased drug concentration.

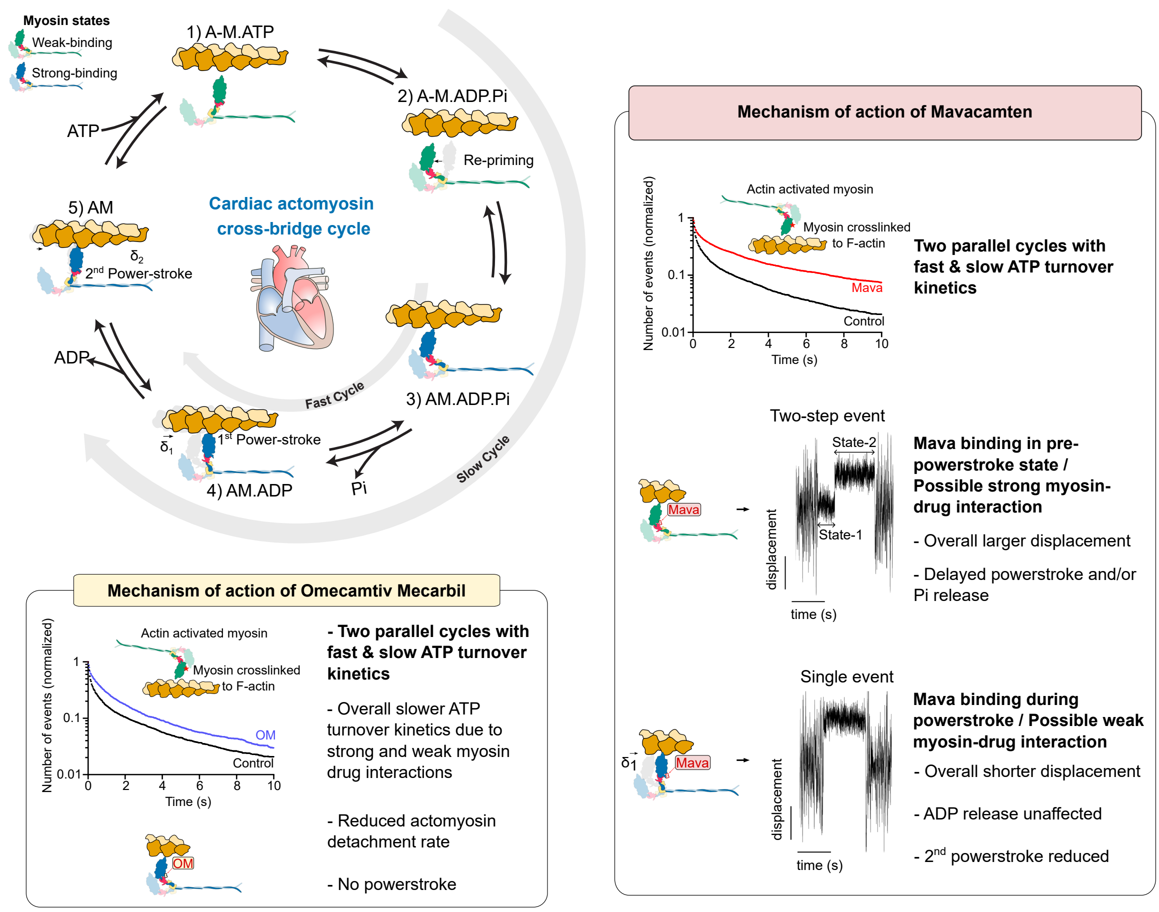


**Figure S14. Simplified graphical summary of mechanism of action of Mavacamten and Omecamtive mecarbil**. The top left panel illustrates the canonical actomyosin crossbridge cycle in cardiac muscle. The two gray arrows indicate the distinct parallel pathways induced by Mava and OM, corresponding to slow and fast cycles depending on the transition state to which each drug binds on the myosin head. The right and bottom panel summarize the effects of these drugs on ATP turnover kinetics, stroke size and actomyosin detachment kinetics, all of which can directly impact the cardiac contractility.

### Supplementary Tables

#### Table S1. Basis for triple-exponential fitting of Basal ATPase data accounting for unspecific Alexa 647-ATP-surface interactions unrelated to myosin

| Conditions | Range of number of events that are ≤128 ms/ region | Average number of events | Number of regions analysed |
|---|---|---|---|
| Mava 1, 2 | 0-4, 0-6 | 1.8, 2.4 | 15 |
| Mava 1 , 2 (background) | 0-5, 0-12 | 1.9, 3.1 | 15 |
| OM 1, 2 | 0-4, 0-6 | 1.8, 2.3 | 15 |
| OM 1, 2 (background) | 0-8, 1-10 | 3.8, 4.3 | 15 |
| Control 1, 2 | 0-6, 0-4 | 3, 3.2 | 15 |
| Control 1, 2 (background) | 0-15, 0-14 | 7, 3.2 | 15 |

Number of fast unspecific Alexa 647-ATP interaction/region/15 minute on S1 myosin and in a background region of interest without myosin. Black: repetition 1, Green: repetition 2. In **Figure 2** in the main manuscript only two exponential components out of 3 are presented. The exclusion of the 3rd exponential process is based on the long lifetimes of the basal ATP turnover under the influence of OM or Mava, which result in sparse sampling over the measured time interval. In contrast, the fast non-specific events occur at a relatively constant rate and makes a larger relative contribution to the overall collected data in the presence of drugs. The contribution is visualized in **Fig S2.** The remaining two processes are normalized towards each other based of their amplitude in the 3-exponential fit.

**Table S2**
**ATP induced detachment, Number of collected events per condition**

| ATP (nM) | $n_{dwells\ (100\ \mu M\ OM)}$ | Repetition* | $n_{dwells\ (Control)}$ | Repetition* | $n_{dwells\ (30\ \mu M\ Mava)}$ | rep |
|---|---|---|---|---|---|---|
| 100 | 309 | 1 | 645 | 2 | 314 | 1 |
| 200 | 967 | 2 | 607 | 2 | 516 | 2 |
| 300 | 1272 | 2 | 653 | 2 | 565 | 2 |
| 500 | 755 | 2 | 653 | 2 | 597 | 2 |
| 750 | 609 | 1 | 318 | 1 | 290 | 1 |
| 1000 | 321 | 1 | 313 | 1 | 317 | 1 |

*Repetition at different occasions with different myosin purifications.

**Table S3.** Parameter values[a] for model with and without OM and MAVA determining shape of free energy diagrams of (human) cardiac actomyosin

| **Parameter** | **Control, (fast)** | **OM (slow)** | **MAVA (slow)** |
|---|---|---|---|
| $x_1$ [b] | 0 nm | 1.5 nm | 1.5 nm |
| $x_{11}$ [b] | 0 nm | 1.5 nm | 1.5 nm |
| $x_2$ [b] | -6.0 nm [c] | -6.0 nm | -6.0 nm |
| $x_3$ [b] | -8.0 nm [c] | -8.0 nm | -8.0 nm |
| $\Delta G_{on}$ (Free energy difference btw MDP and $AMDP_{PP}$) | 3 $k_BT$ | 4 $k_BT$ | -3 $k_BT$ |
| $\Delta G_{PiR}$ (difference btw $AMDP_{PP}$ and $AMDP_{PiR}$) | 0 $k_BT$ | 2 $k_BT$ | -1 $k_BT$ |
| $\Delta G_{PiRL}$ (difference btw $AMDP_{PiR}$ and $AMD_L$) | ln([Pi]/$K_P$) ~3 $k_BT$ | 3 $k_BT$ | 3 $k_BT$ |
| $\Delta G_{LH}$ ($AMD_L$-$AMD_H$) | 11 $k_BT$[e] | -1 $k_BT$ | 2 $k_BT$ |
| $\Delta G_{HD}$ ($AMD_H$-AMD) | 1 $k_BT$[f] | 1 $k_BT$ | 1 $k_BT$ |
| $\Delta G_{ATP}$ | 13.1 + ln ([MgATP]/ ([MgADP][Pi]) = 28.5 $k_BT$ with current parameters | | |
| $k_s$ | 2.8 pN/nm (for positive x)<br>0.2 pN/nm (for negative x for AM/AMD state)<br>2.8 pN/nm (for negative x for all other states)[g] | | |

[a]These parameter values modified from those in [4] referring to two-headed myosin motor fragments at 26 °C. The data from [48] are used instead of human data because they were obtained with full length myosin generally giving higher (and presumably more realistic) [49,50] single molecule mechanics results than data obtained in single motor domains (S1). The free energy difference $\Delta G_{HD}$ has also been modified to be more close to that in ref. [48] . The OM parameter values are developed from those in [4] as starting points.

[b] See Fig. 6a

**Table S4.** Parameter values[a] for model with and without OM and MAVA determining rate functions and equilibrium constants

| **Para-meter** | **Control, (fast)** | **OM (slow)** | **Mava (slow)** |
|---|---|---|---|
| $k_{+3} + k_{-3}$ | 150 $s^{-1}$ | 150 $s^{-1}$ | 150 $s^{-1}$ |
| $K_3$ | 2 | 2 | 2 |
| $k_{-5}$ | 1000 $s^{-1}$ | 1000 $s^{-1}$ | 1000 $s^{-1}$ |
| Kc | 10 mM | 10 mM | 10 mM |
| $k_{on}'$ | 5 $s^{-1}$ | 5 $s^{-1}$ | 5 $s^{-1}$ |
| $k_{Pr+}'$ | 3000 $s^{-1}$ | 3000 $s^{-1}$ | 3000$s^{-1}$ |
| $k_{LH+}$ | 5000 $s^{-1}$ | 0.01 $s^{-1}$ | 5 $s^{-1}$ |
| $k_6$ | 120 $s^{-1}$ | 120 $s^{-1}$ | 120 $s^{-1}$ |
| $k_{det\text{-}escape}$ | NA | 10 $s^{-1}$ | NA |
| $k_{r\text{-}lim\_slow}$ | NA | 1-2 $s^{-1}$ | NA |
| [Pi] | 0.5 mM | 0.5 mM | 0.5 mM |
| [MgATP] | 5 mM | 5 mM | 5 mM |
| $K_1$ | 1.6 $mM^{-1}$ | 1.6 $mM^{-1}$ | 1.6 $mM^{-1}$ |
| $k_2$ | 1400 $s^{-1}$ | 1400 $s^{-1}$ | 1400 $s^{-1}$ |
| $k_{Dr+}$ | NA | 400 $\mu M^{-1}$ $s^{-1}$ | 1000 $\mu M^{-1}$ $s^{-1}$ |
| $k_{Dr-}$ | NA | 2000 $s^{-1}$ | 2000 $s^{-1}$ |
| $k_{Dr2+}$ | - | 25 $s^{-1}$ | 15 $s^{-1}$ |
| $k_{Dr2-}$ | - | 2.5 $s^{-1}$ | 2.5 $s^{-1}$ |
| $k_{basal}$ | 0.06 $s^{-1}$ | 0.06 $s^{-1}$ | 0.06 $s^{-1}$ |

[a] The parameter values primarily originate from myosin motor fragments with heavy chains from human β-cardiac myosin II (but in several cases with murine light chains) at 25°C, ionic strength: ~30 mM - ~120 mM, pH 7-7.5. They are then modified based on simulations in [4].

**Table S5. Initial values for large x, in numerical solutions of differential equations for the standard set of rate constants[a]**

| **[OM] (μM)** | **0** | **0.1** | **1** | **10** | **100** |
|---|---|---|---|---|---|
| **[MT]** | 0.333 | 0.291 | 0.135 | 0.021 | 0.002 |
| **[MDP]** | 0.667 | 0.581 | 0.270 | 0.043 | 0.005 |
| **[MDPDr]** | 0 | 0.012 | 0.054 | 0.085 | 0.090 |
| **[MDPDr2]** | 0 | 0.116 | 0.5405 | 0.851 | 0.903 |
| **Kh[b]** | 2.00 | 2.44 | 6.4 | 46 | 442 |

[a]The numerical values are obtained with $k_{Dr+}$=400 $\mu M^{-1}s^{-1}$, $k_{Dr-}$=2000$s^{-1}$, $k_{Dr2+}$=25$s^{-1}$, $k_{Dr2-}$=2.5$s^{-1}$

[b]This is the calculated hydrolysis equilibrium constant ([MDP]+[$MDP_{Dr}$]+[$MDP_{Dr2}$])/[MT].

**Programming Code - Simnon**

*Numerical solution of systems of differential equations*

The numerical solutions of systems of non-linear ordinary differential equations were obtained using the program Simnon by applying the Runge-Kutta Fehlberg 4/5 algorithm. Simnon was originally developed by the Department of Automatic control at Lund University [26] and was for a period (but no longer) commercially available via SSPA Maritime Consulting AB Gothenbourg, Sweden. We have previously compared the program performance against analytical solutions of differential equations in simple cases [29] and against Monte-Carlo simulations implemented in Matlab for more complex models similar to those used here [28]. Below we copy the Simnon code that we have used with extensive commenting. Whereas there is no commercially available program to run it directly, it should be readily transformed into other programming languages including Matlab, OpenModelica etc. e.g. by using large language models.

In Simnon, generally, a continuous system is first defined with states ("STATE") and their derivatives ("DER") named with "d" followed by the state name. An independent variable or integration variable ("time") is also defined possibly related to another variable (e.g. "x" in the present case). The system of differential equations is written as lines of code starting with "dSTATE" for all states. The initial values of the states are given by the state name followed by colon anywhere in the code withint the Simnon Continuous system. Also, model parameter values are given anywhere by the name of the parameter value followed by colon and the appropriate numerical value. Variables are defined by combining parameter values and independent variables in mathematical expressions. In running a simulation, parameter values and initial values are first set either directly in the program code or from a command dialog window in the Simnon program. Also the integration range is selected before the code is executed with output as requested in the command dialog.

***1. Simnon code for simulating ensemble averages of single turnover ATPase corresponding to frequency distributions in single molecule ATP turnover assays***

*" stands for comment*

```
"declaration of continuous system
continuous system atpase
"declaration of states and derivatives
state a00 a0 a0d a0dr a1 a1dr a4 a5
der da00 da0 da0d da0dr da1 da1dr da4 da5
"a00:MT; a0;MDP; a0d:MDPdr; a0dr:MDPdr2
"a1: strongly attached state without drug before AM
"a1dr: strongly attached state with drug before AM
"a4:AM starting state for single turnover
"a5:AM end state for single turnover
"DR:drug
"Please note, no AM states with drug!!

"definition of integration variable
time t

"System of ordinary differential equations
da4=-koff*a4
```

```
da00=koff*a4-k3p*a00+k3m*a0
da0=k3p*a00-(kon+kdr*DR+k3m)*a0+kdrm*a0d+konm*a1
da0d=kdr*DR*a0+kdr2m*a0dr-(kon+kdr2+kdrm)*a0d
da0dr=kdr2*a0d+kondrm*a1dr-(kdr2m+kondr)*a0dr
da1=kon*(a0+a0d)-(klump+konm)*a1
da1dr=kondr*a0dr-(klumpdr+kondrm)*a1dr
da5=klump*a1+klumpdr*a1dr

"Initial values
a4:1
a00:0
a0:0
a0d:0
a0dr:0
a1:0
a1dr:0
a5:0

"ATP induced detachment at saturating ATP
koff:500
"Reduced from appropriate value of 1200 s-1 to 500 s-1 to increase
"stability in numerical computations.
"No apparent effect on any aspect of the simulated frequency distributions

"Drug concentration in  M
DR:100

"hydrolysis
k3p:100
k3m:50

"Attachment and its reversal without drug
kon:22

"Inverse attachment rate, sometimes approximated to zero
konm:0

"Attachment and its reversal with drug bound. Different for different drugs
kondr:40

"Inverse attachment rate w drug, sometimes approximated to zero
kondrm:0

"transition between attached states btw attachment step and ATP-induced
 "detachment
klump:120

"klumpDr is different for different drugs
klumpdr:1
```

```
"Rate constants associated with drug binding
kdr:400
kdrm:2000
kdr2:25
kdr2m:2.5

a51=1-a5
a55=ln(1-a5)
"The variables a51 and a55 are usually taken as outputs

end
"End of program
```

_Solving continuous system:_ Parameter values and initial values are first set in the command dialog followed by requested output and integration range. The running of the continuous system is then initiated

***2. Simnon code for simulating power strokes, corresponding to averages of a large number of single molecule mechanics data***

```
continuous system pstrok
"Simulation of either fast weakly drug bound path or slow strongly drug
"bound path in single molecule mechanics depending on parameter values.
"NOTE! In this simulation we consider each cycle SEPARATELY and do not
" include effects of drug-binding and drug-release

"declaration of states and derivatives below
state a00 a0 a11 a1 a2 a1t a3 a4
der da00 da0 da11 da1 da2 da1t da3 da4

"Please NOTE! Different naming of states and rate constants than in the
" ATPase program
"a00:MT; a0:MDP; a1:AMDPPP
"a1t:AMDPiR;  a11:AMDL   a2:AMDH;   a3:AMD   a4:AM

"integration variable
time t

"Differential equations below
da00=-k3*a00+k3m*a0+koff*a4+krlim*a11
da0=k3*a00+konm*a1-(kon+k3m)*a0
da1=kon*a0+kprm*a1t-(konm+kprp)*a1
da1t=kprp*a1+kprel*(Pi/Kp)*a11-(kprm+kprel)*a1t
da11=kLHm*a2+kprel*a1t-(kLH+krlim+kprel*(Pi/Kp))*a11
da2=kLH*a11+k5m*a3-(k5+kLHm)*a2
da3=k5*a2-(k6+k5m)*a3
da4=k6*a3-koff*a4
```

```
"den: denominator for normalisation
den_l=(a1+a11+a1t+a2+a3+a4)
"l:average power stroke distance
l=((a11+a1t)*h0+a2*(h0+h)+(a3+a4)*(h0+h+h1))/den_l
h0=x1-x11
h=x11-x2
h1=x2-x3

"average attachment coordinate
x=F_set/k            "F_set=0 in all present calculations

"Initial values
a00:0
a0:0
a1:1
a1t:0
a11:0
a2:0
a22:0
a3:0
a4:0
"Definition of strain-dependent rate constants
"Translation to main Fig. 6 and Tables S3-S4 terminology
"attachment transitions between a0 and a1 (MDP and AMDPPP)
"kon0: kon ; kon0m: kon  (but   kon:kon+(x); konm: kon-(x); Gon: Gon;
kon0:5  "only used for reversal of attachment
kon00:0  "used to ingnore attachment
exp_s0=(1/2)*F_set*F_set/(k*4)   "F_set=0 in the present calculations
kon_l00=kon00*exp(Gon-exp_s0/2)
kof_l00=kon0*exp(exp_s0/2)
kon=if kon_l00<0.0001 then 0 else kon_l00
konm=if kof_l00>fc then fc else kof_l00

"a1 to a1t and back; AMDPPP and AMDPPiR
"kprp:kpr+(x); kprm: kpr-(x); GPiR: GPiR; F_set and h0 defined below
kp_plus:3000
"Parameter values vary depending on drug
exp_ps0=GPiR/2+(1/2)*F_set*F_set/(k*4)-(k/2)*(x+h0)*(x+h0)/4
exp_ps0m=GPiR/2+(k/2)*(x-h0)*(x-h0)/4-(1/2)*F_set*F_set/(k*4)
"F_set=0 always in present calculations
kp_plur0=kp_plus*exp(exp_ps0)
kp_min0=kp_plus*exp(-exp_ps0m)
kprp=if kp_plur0<0.0001 then 0 else kp_plur0
kprm=if kp_min0>fc then fc else kp_min0

"Pi release and re-binding (btw a1t and a11)
kprel:10000
Pi:0.5
Kp:10
"Dissociation constant of Pi
```

```
"transitions between a11 and a2 (Huxley-Simmons transition;AMDL "-
>AMDH)
kLH0:5000
exp_12=(1/2)*F_set*F_set/(k*4)-(k/2)*(x+h+h0)*(x+h+h0)/4
"forward
exp_12m=(k/2)*(x-(h+h0))*(x-(h+h0))/4-(1/2)*F_set*F_set/(k*4)
"backward
"F_set=0 always in present calculations
kplusf=kLH0*exp(GLH+exp_12)
kLH=if (kplusf>fc1) then fc1 else kplusf

kpfh=kLH0*exp(GLH+exp_12m)
kvot1=kpfh/kLH0
Keq12=if kvot1>fc1 then fc1 else if kvot1<0.00001 then 0.00001 else kvot1
kLHm=kLH0/Keq12

"Transition AMDH <-> AMD
kmin5:1000
xh=h1+F_set/k
xhh1=(x-h-h1-x3)
exp5=(1/2)*F_set*F_set/(4*k)-(k/2)*xh*xh/4
kplusf5=if kmin5*exp(GHD+exp5)>fc1 then fc1 else kmin5*exp(GHD+exp5)
kvot5=kmin5/kplusf5
k5=if (kplusf5>fc) then fc else kplusf5
k5m=if (kplusf5>(fc-1)) then kvot5*kplusf5 else kmin5

"a4 -> a0  (AM -> MT)
"Detachment rate constant koff(x)=k2[ATP]/((1/K1)+[ATP])
"xcrit:0, critical strain-parameter for x-dependent detachment rate.
" Set "to 0 "when cross-bridge stiffness is non-linear (km=0.2 pn/Nm)
"for AM state at "x<x3
"fb=exp(abs(F_set)*xcrit/4)
"fb=1 when xcrit=0

fb=1
koff0=fb*k2*ATP/((1/(K1))+ATP)
koff=if x<(x1+4*xlimit) then koff0 else 0
"Remaining parameter values below
"Clamping force. Clamping to zero force is used throughout the paper.
F_set:0

"Hydrolysis equilibrium
k3:100
k3m:50

"ATP binding constant and ATP-induced dissociation constant
k2:1400
K1:1.6  "(mM-1)
"Very low ATP concentration (mM) to ignore ATP-induced detachment generally
```

```
ATP:0.0001

"ADP release rate constant
k6:120
"Escape route with OM or other drug
krlim:10

"DEFINITION OF FREE ENERGY DIAGRAMS BELOW
"Differences in free energy between neigboring states
" - change between ctrl "and between different drugs
Gon:1.5
GPir:1
GLH:11
GHD:3

"x-positions of minima of free energy diagrams
x1:0
x11:0
x2:-6
x3:-8

"constants rmax
fc:50000
fc1:300000
"constant to limit calculations to certain x-range for stability
xlimit:3

"Cross-bridge stiffness (only range of positive cross-bridge strains
" considered in these calculations eliminating need to consider
" non-linearity of cross-bridge elasticity)
k:2.8
end
“end of program
```

Solving continuous system: As in the case 1 above, parameter values and initial values are first set in the command dialog followed by requested output and integration range. The running of the continuous system is then initiated. Here we focus on the output l as a measure of the displacement during the power stroke.

#### 3. Simnon code for simulating force-velocity relationships including approximation of maximum isometric force.

```
continuous system pvheart
"Declaration of program start above
"Declaration of states below
state a00 a0 a11 a1 a2 a3 a4
state a0d a0dr a1dr a11dr
"a00:MT; a0:MDP;
"a1:AMDP  a11:AMDL   a2:AMDH;  a3:AMD  a4:AM
```

```
"a0d:MDPDr  a0dr: MDPDr2  a1dr: AMDPPPDr  a11dr: AMDLDr

state i atpas Na
"i: total force vs x
"atpas: ATPase vs x. Na: Number of attached x-bridges vs x

"Declaration of derivatives below
der da00 da0 da11 da1 da2 da3 da4
der da0d da0dr da1dr da11dr
der di datpas dNa
"di: derivative of total force vs x
"datpas: derivative of ATPase vs x. dNa: derivative of Na vs x

"Definition of integration variable
time t

"Definition of position-variable in terms of t
x=10-t

"Differential equations d[state]/dx
da00=(-k3*a00+krlim*a11dr+k3m*a0+koff*a4)/v
da0=(k3*a00+konm*a1+kdrm*a0d-(kdr*DR+kon+k3m)*a0)/v
da1=(kon*(a0+a0d)+kprm*a11+kdr2m*a1dr-(konm+kprp+kdr20*DR)*a1)/v
da11=(kLHm*a2+kprp*a1+kdr2m*a11dr-(kLH+kprm+kdr20*DR)*a11)/v
da2=(kLH*a11+kLHdr*a11dr+kmin5*a3-(kLHm+k5)*a2)/v
da3=(k5*a2-(k6+kmin5)*a3)/v
da4=(k6*a3-koff*a4)/v
"With drug bound below
da0d=(kdr*DR*a0+kdr2m*a0dr-(kdrm+kdr2+kon)*a0d)/v
da0dr=(kdr2*a0d+konmdr*a1dr-(kondr+kdr2m)*a0dr)/v
pa1dr=(konmdr+kprpdr+kdr2m)
da1dr=(kondr*a0dr+kprmdr*a11dr+kdr20*DR*a1-(pa1dr)*a1dr)/v
da11dr=(kprpdr*a1dr+kdr20*DR*a11-(kLHdr+kprmdr+kdr2m+krlim)*a11dr)/v
"implementation with non-linear cross-bridge elasticity
"with lower stiffness, km for x<x3
ksdr=if x>x3 then k else km
ks=if x>x3 then k else km
"ksdr is set to ksdr=k in the present simulations
"but it allows different stiffness of certain states
 "in the presence of drug.
"ks relates to non-linear cross-bridge elasticity for states
"a3 and a4


"Differential equations below performing integration to obtain
" average Na, force "and ATPase, along x starting at x=10 nm.
"Argument (x) for states not shown.

dNa=(1/36)*(a1+a11+a2+a3+a4+a1dr+a11dr)
f0drL=(k/36)*(a2*(x-x2)+(a1+a11)*(x-x1))
```

```
f0drH=(ks/36)*(a3+a4)*(x-x3)
f0dr=f0drL+f0drH
fdr=(ksdr/36) *(a11dr*(x-x11d)+a1dr*(x-x1d))
di=f0dr+fdr
datpas=(1/36)*koff*a4

"Setting velocity (the value is changed to obtain entire force-velocity relation)
v:0.5

"Initial values for x=10 nm where integration starts.
"These initial values are changed in the presence of drug,
"being calculated as described in the Methods by solving
" the corresponding system of linear "equations in Mathematica
a00:0.33
a0:0.67
a0d:0.
a0dr:0
a1:0
a1dr:0
a11:0
a11dr:0
a2:0
a3:0
a4:0

"Definition of strain-dependent rate constants
"attachment transitions between a0 and a1 (MDP and AMDPPP)
"kon0: kon ; kon:kon+(x); konm: kon-(x); Gon: Gon; gamma:gamma
"see SI methods description and Tables S3-S4 for
"parameter values. Division by 4 here and below because
"1kBT 4 pN nm with k in units of pN/nm
gamma:2
exp_s0=(k/2)*(x-x1)*(x-x1)/4
kon1=if x>2*x3 then kon0*exp((1/gamma)*(Gon-exp_s0)) else 0
kofl=kon0*exp((1-1/gamma)*(-Gon+exp_s0))
kon=if kon1<0.0001 then 0 else kon1
konm=if kofl>fc then fc else kofl

"Attachment transitions between a0 and a1 (MDPDr and AMDPPPDr)
"with drug strongly bound
"kon0dr: konDr ; kondr:konDr+(x); konmdr: konDr-(x);
"Gondr: GonDr   allows shift of max attachment probability
"away from minimum of free  energy of AMDPPdr-state
exp_s0o=(ksdr/2)*(x-x1d)*(x-x1d)/4
kon1dr=if x>2*x3 then kon0dr*exp((1/gamma)*(Gondr-exp_s0o)) else 0
kofldr= kon0dr*exp((1-1/gamma)*(-Gondr+exp_s0o))
kondr=if kon1dr<0.0001 then 0 else kon1dr
konmdr=if kofldr>fc then fc else kofldr

"kprp:kpr+(x); kprm: kpr-(x); GPiR: GPiR
```

```
exp_ps0=GPiR/2-(k/16)*(x-x11)*(x-x11)+(k/16)*(x-x1)*(x-x1)
"becomes exp_ps0=GPiR/2 if x1=x11 as here
kp_plur0=kp_plus*exp(exp_ps0)
kp_min0=kp_plus*exp(-exp_ps0)*Pi/(Kp+Pi)
kprp=if kp_plur0<0.0001 then 0 else kp_plur0
kprm=if kp_min0>fc then fc else kp_min0

"With drug
"kprpdr:kprDr+(x); kprmdr: kprDr-(x); GPiR: GPiRDr
exp_s0d=GPiRDr/2-(k/16)*(x-x11d)*(x-x11d)+(k/16)*(x-x1d)*(x-x1d)
"becomes exp_s0=GPiRDr/2 if x1=x11 as here
kp_pludr=kp_plud0*exp(exp_s0d)
kp_m0dr=kp_plud0*exp(-exp_s0d)*Pi/(Kpo+Pi)
kprpdr=if kp_pludr<0.0001 then 0 else kp_pludr
kprmdr=if kp_m0dr>fc then fc else kp_m0dr

"transitions between a11 and a2 (Huxley-Simmons
"AMDL->AMDH) without strongly bound drug
exp12=(k/2)*(x-x11)*(x-x11)/4-(k/2)*(x-x2)*(x-x2)/4
kplusf=KLH0*exp(GLH+exp12)
kLH=if (kplusf>fc1) then fc1 else kplusf
kLHm=kLH0

"Transitions between a11dr and a2 (Huxley-Simmons
"AMDL->AMDH) with strongly bound drug
"kLHdr- and kLHdr+ differ between drugs
exp12dr=(k/2)*(x-x11d)*(x-x11d)/4-(k/2)*(x-x2)*(x-x2)/4
kplusfd= kLHdr0p*exp(GLHdr+exp12dr)
kplusd0=if (kplusfd>fc1) then fc1 else kplusfd
kLHdr=if kplusd0<0.00001 then 0 else kplusd0

"a2 -> a3  (AMDH  -> AMD) transition
"k5:k5(x); kmin5:k5-(x)
exp5=(k/2)*(x-x2)*(x-x2)/4-(ks/2)*(x-x3)*(x-x3)/4
kplusf5=if kmin5*exp(GHD+exp5)>fc1 then fc1 else kmin5*exp(GHD+exp5)
k5=if (kplusf5>fc) then fc else kplusf5

"a4 -> a0  (AM -> MT)
"Detachment rate constant koff(x)=k2[ATP]/((1/K1)+[ATP])
"xcrit:0 (critical strain-parameter for x-dependent
"detachment rate. Set to 0 "when cross-bridge stiffness is
"non-linear (km=0.2 pn/Nm) for AM state at "x<x3
xcrit:0
fb=exp(ks*abs(x-x3)*xcrit/4)
"fb=1 when xcrit=0 as is the case here
"ATP=[MgATP]
koff0=fb*k2*ATP/((1/(K1))+ATP)
koff=if x<(x1+4*xlimit) then koff0 else 0

"Parameter values below These values are easily modified
```

```
"from command window when running program
"Rate constants (s-1)
"Hydrolysis equlibrium: k3:k3+; k3m: k3-;
k3:100
k3m:50

"Drug binding
"kdr:kdr+; kdrm:kdr-;"kdr2:kdr2+; kdr2m:kdr2-;
kdr:1000
kdrm:2000
kdr2:15
kdr2m:2.5
kdr20=if x<x1+deltaxo then 0 else kdr200
kdr200=catt*DR*kdr2/((kdrm/kdr)+DR)
catt:1
deltaxo:0
"Drug concentration( M)
DR:0

"Strong cross-bridge attachment and its reversal
"kon0dr different between OM and Mava
kon0:5
kon0dr:25
"Differences in free energy between neigboring states
"- change between ctrl "and between different drugs
Gon:3
Gondr:-3

"Transitions AMDPPP ->AMDL "including Pi dissociation
"Rate constant of transition
kp_plus:3000
kp_plud0:3000
"Pi dissociation constant (mM)
Kp:10
Kpo:10
"Pi concentration (mM)
Pi:0.5
"Difference between free energy levels
GPir:0
GPiRDr:-1
"Power-stroke and its reversal
kLH0:5000
kLHdr0p:5
"Difference between free energy levels
GLH:11
"GLHdr is different for different drugs
GLHdr:2

"Strain-dependent transition before ADP-release associated with second “stroke
kmin5:1000
```

```
GHD:1

"ADP release rate constant
k6:120
"Escape route with OM or other drug
krlim=if x>x1d+0.5 then 0 else krlim0*(x1d+0.5-x)
krlim0:0

"ATP binding constant (K1; mM^-1) and ATP-induced dissociation
"rate constant (k2)
K1:1.6
k2:1400

"Substrate concentration (mM)
ATP:5

"x-positions of minima of free energy diagrams
x1:0
x11:0
x1d:1.5
x11d:1.5
x2:-6
x3:-8

"Cross-bridge stiffness
"km: stiffness for a3 and a4-states for x<x3
"k:stiffness under all other conditions than listed above
k:2.8
km:0.2

"constants rmax
fc:50000
fc1:300000
kona:50000

"constant to limit calculations to certain x-range for stability
xlimit:3

"End of program
end
```

As in the cases 1-2 above, parameter values and initial values (Excel table below) are first set in the command dialog followed by requested output and integration range. The running of the continuous system is then initiated. Here we focus on the outputs "i" (force) and "Na" (number of attached cross-bridges). Integration was limited to x=10 nm to x = -15 to -17 nm for velocity v = 0.5 nm/s (approximating isometric contraction) with integration running along the negative x-axis. The integration was limited to the range x=10 nm to x = -65 nm for maximum velocity of shortening (v <1450 nm/s). Negligible changes in force and number of attached cross-bridges were seen with 10 % changes in these integration ranges.

#### 4. *Mathematica code for solving systems of linear equations*

Drug

*a) Equilibrium MT <-> MDP <-> $MDP_{Dr}$ <-> $MDP_{Dr2}$ and basal ATPase*

```
In[ ]:= LinearSolve[{{-k3p, k3m, 0, 0}, {k3p, -(k3m + kdr * Drug), kdrm, 0},
   {0, kdr * Drug, -(kdrm + kdr2), kdr2m}, {0, 0, kdr2, -kdr2m},
   {1, 1, 1, 1}}, {0, 0, 0, 0, 1}]
```

Out[ ]=

$$\left\{\frac{k3m\ kdr2m\ kdrm}{Drug\ k3p\ kdr\ kdr2 + Drug\ k3p\ kdr\ kdr2m + k3m\ kdr2m\ kdrm + k3p\ kdr2m\ kdrm},\right.$$
$$\frac{k3p\ kdr2m\ kdrm}{Drug\ k3p\ kdr\ kdr2 + Drug\ k3p\ kdr\ kdr2m + k3m\ kdr2m\ kdrm + k3p\ kdr2m\ kdrm},$$
$$\frac{Drug\ k3p\ kdr\ kdr2m}{Drug\ k3p\ kdr\ kdr2 + Drug\ k3p\ kdr\ kdr2m + k3m\ kdr2m\ kdrm + k3p\ kdr2m\ kdrm},$$
$$\left.\frac{Drug\ k3p\ kdr\ kdr2}{Drug\ k3p\ kdr\ kdr2 + Drug\ k3p\ kdr\ kdr2m + k3m\ kdr2m\ kdrm + k3p\ kdr2m\ kdrm}\right\}$$

This code, slightly modified, is also used for solving the basal ATPase vs drug concentration from the product $k_{basal}$[MDP], assuming that the M and MD states are transient intermediates and that only the MDP state undergoes basal ATP turnover.

*b) Steady-state actin-activated ATPase for the simplified scheme in Fig. S1.*

Symbolically, the input to solve the steastdy-state ATPase is as follows in Mathematica code and the corresponding standard Matrix notation respectively:

```
LinearSolve[{{-koff, 0, 0, klump, 0, 0, klumpesc + klumpcan},
  {koff, -k3p, k3m, 0, 0, 0, 0},
  {0, k3p, -(k3m + kon + kDrp * DR), 0, kDrm, 0, 0},
  {0, 0, kon, -(klump + 0), kon, 0, 0},
  {0, 0, kDrp * DR, 0, -(kDr2p + kDrm + kon), kDr2m, 0},
  {0, 0, 0, 0, kDr2p, -(konDr + kDr2m), 0},
  {0, 0, 0, 0, 0, konDr, -(0 + klumpesc + klumpcan)},
  {1, 1, 1, 1, 1, 1, 1}}, {0, 0, 0, 0, 0, 0, 0, N}]
```

DR: drug concentration and rate constant names adapted from Fig. 6a to fit Mathematica requirments

$$
\begin{pmatrix} 0 \\ 0 \\ 0 \\ 0 \\ 0 \\ 0 \\ 0 \\ 1 \end{pmatrix}
=
\begin{pmatrix}
-k_{off} & 0 & 0 & k_{lump} & 0 & 0 & k_{lump\text{-}can}+k_{lump\text{-}esc} \\
k_{off} & -k_{3+} & k_{3-} & 0 & 0 & 0 & 0 \\
0 & k_{3+} & -(k_{3-}+k_{on}+k_{Dr+}DR) & 0 & k_{Dr-} & 0 & 0 \\
0 & 0 & k_{on} & -k_{lump} & k_{on} & 0 & 0 \\
0 & 0 & k_{Dr+}DR & 0 & -(k_{Dr-}+k_{Dr2}) & k_{Dr2-} & 0 \\
0 & 0 & 0 & 0 & k_{Dr2+} & -(k_{Dr2}+k_{onDr}) & 0 \\
0 & 0 & 0 & 0 & 0 & k_{onDr} & -k_{lumpdr} \\
1 & 1 & 1 & 1 & 1 & 1 & 1
\end{pmatrix}
\begin{pmatrix} [AM] \\ [MT] \\ [MDP] \\ [AMDP_{PP}] \\ [MDP_{Dr}] \\ [MDP_{Dr2}] \\ [AMDP_{PPDr}] \end{pmatrix}
$$

DR: drug concentration

In order to obtain the steady-state ATPase activity or flux, the above linear system of equations were first solved after inserting numerical values of all rate constants. The ATP turnover rates for different drug concentrations were then derived as $k_{off}$ [AM].

### Supplementary Movie Legends

**Supplementary Videos 1-3. Basal single molecule ATPase of myosin S1$^{E}$.**

**Control (Movie 1), 30 µM Mava (Movie 2) and 100 µM OM (Movie 3) conditions.** The first frame is a standard deviation projection of the video obtained from ~30000 frames using ImageJ. The subsequent frames represent a video of 40x40 µm$^2$ size recorded at 1/0.032 frames s$^{-1}$ with playback speed in real-time. Each image frame has been converted to 8-bit and background subtraction has been performed using a rolling ball algorithm (rolling ball radius 5 pixels).

**Supplementary Videos 4-6. Actin-activated single molecule ATPase for myosin S1$^{E}$ cross-linked to actin for Control (Movie 4), 30 µM Mava (Movie 5) and 100 µM OM (Movie 6) conditions.** The first frame is a standard deviation projection of the video obtained from ~30000 frames using ImageJ. The subsequent frames represent a video of 40x40 µm$^2$ size recorded at 1/0.032 frames s$^{-1}$ with playback speed in real-time. Each image frame has been converted to 8-bit and background subtraction has been performed using a rolling ball algorithm (rolling ball radius 5 pixels).

**Supplementary Videos 7-9. Actin-activated single molecule ATPase data for βM-II cross-linked to actin for Control (7), 30 µM Mava (8) and 100 µM OM (9) conditions.** The first frame is a standard deviation projection of the videos obtained from ~30000 frames using ImageJ. The subsequent frames represent a videos of Control 94x37 µm$^2$ (Movie 7), Mava 60x15 µm$^2$ (Movie 8), OM 71x27 µm$^2$ (Movie 9) size recorded at 1/0.032 frames s$^{-1}$ with playback speed in real-time. Each image frame has been converted to 8-bit and background subtraction has been performed using a rolling ball algorithm (rolling ball radius 5 pixels).

### Supplementary References


1 Rahman, M. A., Usaj, M., Rassier, D. E. & Månsson, A. Blebbistatin Effects Expose Hidden Secrets in the Force-Generating Cycle of Actin and Myosin. *Biophys. J.* **115**, 386-397, doi:10.1016/j.bpj.2018.05.037 (2018).

2 Månsson, A. Hypothesis: Single Actomyosin Properties Account for Ensemble Behavior in Active Muscle Shortening and Isometric Contraction. *International journal of molecular sciences* **21**, doi:10.3390/ijms21218399 (2020).

3 Moretto, L. *et al.* Multistep orthophosphate release tunes actomyosin energy transduction. *Nat Commun* **13**, 4575, doi:10.1038/s41467-022-32110-9 (2022).

4 Månsson, A. Mechanistic insights into effects of the cardiac myosin activator omecamtiv mecarbil from mechanokinetic modelling. *Front Physiol* **16**, 1576245, doi:10.3389/fphys.2025.1576245 (2025).

5 Scellini, B. *et al.* Mavacamten has a differential impact on force generation in myofibrils from rabbit psoas and human cardiac muscle. *J Gen Physiol* **153**, doi:10.1085/jgp.202012789 (2021).

6 Mansson, A. Theoretical treatment of tension transients in muscle following sudden changes in orthophosphate concentration: implications for energy transduction. *J. Muscle Res. Cell Motil.* **46**, 193-213, doi:10.1007/s10974-025-09698-8 (2025).

7 Mansson, A. A mechanokinetic actomyosin model predicts different orthophosphate sensitivities of force and ATP turnover rate during isometric muscle contraction. *Front Physiol* **16**, 1659772, doi:10.3389/fphys.2025.1659772 (2025).

8 Auguin, D. *et al.* Omecamtiv mecarbil and Mavacamten target the same myosin pocket despite opposite effects in heart contraction. *Nat Commun* **15**, 4885, doi:10.1038/s41467-024-47587-9 (2024).

9 Woody, M. S. *et al.* Positive cardiac inotrope omecamtiv mecarbil activates muscle despite suppressing the myosin working stroke. *Nature communications* **9**, 3838, doi:10.1038/s41467-018-06193-2 (2018).

10 Liu, Y., White, H. D., Belknap, B., Winkelmann, D. A. & Forgacs, E. Omecamtiv Mecarbil modulates the kinetic and motile properties of porcine beta-cardiac myosin. *Biochemistry* **54**, 1963-1975, doi:10.1021/bi5015166 (2015).

11 Malik, F. I. *et al.* Cardiac myosin activation: a potential therapeutic approach for systolic heart failure. *Science* **331**, 1439-1443, doi:10.1126/science.1200113 (2011).

12 Rohde, J. A., Thomas, D. D. & Muretta, J. M. Heart failure drug changes the mechanoenzymology of the cardiac myosin powerstroke. *Proc Natl Acad Sci U S A* **114**, E1796-E1804, doi:10.1073/pnas.1611698114 (2017).

13 Liu, C., Kawana, M., Song, D., Ruppel, K. M. & Spudich, J. A. Controlling load-dependent kinetics of beta-cardiac myosin at the single-molecule level. *Nat Struct Mol Biol* **25**, 505-514, doi:10.1038/s41594-018-0069-x (2018).

14 Kawas, R. F. *et al.* A small-molecule modulator of cardiac myosin acts on multiple stages of the myosin chemomechanical cycle. *J. Biol. Chem.* **292**, 16571-16577, doi:10.1074/jbc.M117.776815 (2017).

15 Green, E. M. *et al.* A small-molecule inhibitor of sarcomere contractility suppresses hypertrophic cardiomyopathy in mice. *Science* **351**, 617-621, doi:10.1126/science.aad3456 (2016).

16 McMillan, S. N., Pitts, J. R. T., Barua, B., Winkelmann, D. A. & Scarff, C. A. Mavacamten inhibits myosin activity by stabilizing the myosin interacting-heads motif and stalling motor force generation. *Sci Adv* **12**, eaea9335, doi:10.1126/sciadv.aea9335 (2026).

17 Rohde, J. A., Roopnarine, O., Thomas, D. D. & Muretta, J. M. Mavacamten stabilizes an autoinhibited state of two-headed cardiac myosin. *Proc. Natl. Acad. Sci. U. S. A.* **115**, E7486-E7494, doi:10.1073/pnas.1720342115 (2018).

18 Swenson, A. M. *et al.* Omecamtiv Mecarbil Enhances the Duty Ratio of Human beta-Cardiac Myosin Resulting in Increased Calcium Sensitivity and Slowed Force Development in Cardiac Muscle. *J Biol Chem* **292**, 3768-3778, doi:10.1074/jbc.M116.748780 (2017).

19 Berg, A., Velayuthan, L. P., Tagerud, S., Usaj, M. & Månsson, A. Probing actin-activated ATP turnover kinetics of human cardiac myosin II by single molecule fluorescence. *Cytoskeleton (Hoboken)*, doi:10.1002/cm.21858 (2024).
20 Planelles-Herrero, V. J., Hartman, J. J., Robert-Paganin, J., Malik, F. I. & Houdusse, A. Mechanistic and structural basis for activation of cardiac myosin force production by omecamtiv mecarbil. *Nature communications* **8**, 190, doi:10.1038/s41467-017-00176-5 (2017).
21 Llinas, P. *et al.* How actin initiates the motor activity of Myosin. *Dev Cell* **33**, 401-412, doi:10.1016/j.devcel.2015.03.025 (2015).
22 Dantzig, J. A., Goldman, Y. E., Millar, N. C., Lacktis, J. & Homsher, E. Reversal of the cross-bridge force-generating transition by photogeneration of phosphate in rabbit psoas muscle fibres. *J Physiol* **451**, 247-278 (1992).
23 Huxley, A. F. & Simmons, R. M. Proposed mechanism of force generation in striated muscle. *Nature* **233**, 533-538 (1971).
24 Eisenberg, E. & Hill, T. L. A cross-bridge model of muscle contraction. *Prog Biophys Mol Biol* **33**, 55-82, doi:10.1016/0079-6107(79)90025-7 (1978).
25 Månsson, A. Actomyosin-ADP states, inter-head cooperativity and the force-velocity relation of skeletal muscle. *Biophys. J.* **98**, 1237-1246 (2010).
26 Elmqvist, H. *Simnon: An Interactive Simulation Program for Nonlinear Systems : User's Manual*. Vol. 7502 208 (Department of Automatic Control, Lund Institute of Technology, Institutionen för Reglerteknik, 1975).
27 Månsson, A. Comparing models with one versus multiple myosin-binding sites per actin target zone: The power of simplicity. *J. Gen. Physiol.* **151**, 578-592, doi:10.1085/jgp.201812301 (2019).
28 Månsson, A. & Rassier, D. E. Insights into Muscle Contraction Derived from the Effects of Small-Molecular Actomyosin-Modulating Compounds. *International journal of molecular sciences* **23**, doi:10.3390/ijms232012084 (2022).
29 Månsson, A. Theoretical treatment of tension transients in muscle following sudden changes in orthophosphate concentration: implications for energy transduction. *J. Muscle Res. Cell Motil.*, doi:10.1007/s10974-025-09698-8 (2025).
30 Wang, T. *et al.* Cardiac ventricular myosin and slow skeletal myosin exhibit dissimilar chemomechanical properties despite bearing the same myosin heavy chain isoform. *J Biol Chem* **298**, 102070, doi:10.1016/j.jbc.2022.102070 (2022).
31 Scellini, B. *et al.* Myosin Isoform-Dependent Effect of Omecamtiv Mecarbil on the Regulation of Force Generation in Human Cardiac Muscle. *Int J Mol Sci* **25**, doi:10.3390/ijms25189784 (2024).
32 Nagy, L. *et al.* The novel cardiac myosin activator omecamtiv mecarbil increases the calcium sensitivity of force production in isolated cardiomyocytes and skeletal muscle fibres of the rat. *Br J Pharmacol* **172**, 4506-4518, doi:10.1111/bph.13235 (2015).
33 Governali, S. *et al.* Orthophosphate increases the efficiency of slow muscle-myosin isoform in the presence of omecamtiv mecarbil. *Nat Commun* **11**, 3405, doi:10.1038/s41467-020-17143-2 (2020).
34 ter Keurs, H. E. *et al.* Force, sarcomere shortening velocity and ATPase activity. *Adv. Exp. Med. Biol.* **538**, 583-602; discussion 602 (2003).
35 Iwamoto, H., Oiwa, K., Suzuki, T. & Fujisawa, T. X-ray diffraction evidence for the lack of stereospecific protein interactions in highly activated actomyosin complex. *J. Mol. Biol.* **305**, 863-874, doi:10.1006/jmbi.2000.4334 (2001).
36 Huang, Y. P., Kimura, M. & Tawada, K. Covalent crosslinking of myosin subfragment-1 and heavy meromyosin to actin at various molar ratios: different correlations between ATPase activity and crosslinking extent. *J. Muscle Res. Cell Motil.* **11**, 313-322, doi:10.1007/BF01766669 (1990).
37 Duong, A. M. & Reisler, E. Binding of myosin to actin in myofibrils during ATP hydrolysis. *Biochemistry (Mosc).* **28**, 1307-1313 (1989).
38 Glyn, H. & Sleep, J. Dependence of adenosine triphosphatase activity of rabbit psoas muscle fibres and myofibrils on substrate concentration. *J. Physiol. (Lond).* **365**, 259-276 (1985).

39 Arata, T. Chemical crosslinking of myosin subfragment-1 to F-actin in the presence of nucleotide. *J Biochem* **96**, 337-347, doi:10.1093/oxfordjournals.jbchem.a134843 (1984).
40 Brenner, B. & Eisenberg, E. Rate of force generation in muscle: correlation with actomyosin ATPase activity in solution. *Proc Natl Acad Sci U S A* **83**, 3542-3546, doi:10.1073/pnas.83.10.3542 (1986).
41 Biosca, J. A., Greene, L. E. & Eisenberg, E. Binding of ADP and ATP analogs to cross-linked and non-cross-linked acto X S-1. *J. Biol. Chem.* **261**, 9793-9800 (1986).
42 Stein, L. A., Greene, L. E., Chock, P. B. & Eisenberg, E. Rate-limiting step in the actomyosin adenosinetriphosphatase cycle: studies with myosin subfragment 1 cross-linked to actin. *Biochemistry* **24**, 1357-1363 (1985).
43 Mornet, D., Bertrand, R., Pantel, P., Audemard, E. & Kassab, R. Structure of the actin-myosin interface. *Nature* **292**, 301-306, doi:10.1038/292301a0 (1981).
44 Yamamoto, K. Shift of binding site at the interface between actin and myosin. *Biochemistry (Mosc).* **29**, 844-848 (1990).
45 Yamamoto, K. Binding manner of actin to the lysine-rich sequence of myosin subfragment 1 in the presence and absence of ATP. *Biochemistry (Mosc).* **28**, 5573-5577, doi:10.1021/bi00439a035 (1989).
46 Velayuthan, L. P., Moretto, L., Tagerud, S., Usaj, M. & Månsson, A. Virus-free transfection, transient expression, and purification of human cardiac myosin in mammalian muscle cells for biochemical and biophysical assays. *Scientific reports* **13**, 4101, doi:10.1038/s41598-023-30576-1 (2023).
47 Pilagov, M., Heling, L., Walklate, J., Geeves, M. A. & Kad, N. M. Single-molecule imaging reveals how mavacamten and PKA modulate ATP turnover in skeletal muscle myofibrils. *J. Gen. Physiol.* **155**, doi:10.1085/jgp.202213087 (2023).
48 Hwang, Y., Washio, T., Hisada, T., Higuchi, H. & Kaya, M. A reverse stroke characterizes the force generation of cardiac myofilaments, leading to an understanding of heart function. *Proc Natl Acad Sci U S A* **118**, doi:10.1073/pnas.2011659118 (2021).
49 Månsson, A., Usaj, M., Moretto, L. & Rassier, D. E. Do Actomyosin Single-Molecule Mechanics Data Predict Mechanics of Contracting Muscle? *International journal of molecular sciences* **19**, doi:10.3390/ijms19071863 (2018).
50 Rassier, D. E. & Månsson, A. Mechanisms of myosin II force generation. Insights from novel experimental techniques and approaches. *Physiol. Rev.* **105**, doi:10.1152/physrev.00014.2023 (2025).
